# Branched-Chain α-Keto Acid Mitochondrial Uptake Regulates Fatty Acid Metabolism in Adipose Tissue

**DOI:** 10.64898/2026.11.15.766772

**Authors:** Yunxia Liu, Zhao Liu, Mei Li, Liping Su, Jiaen Lin, Geok Lin Chua, Shuxun Ren, Ye Sing Tan, Michaela Chocholouskova, Xuejiao Zhang, Qiuzhong Zhou, Bin Dong, Colin Ming-Xiang Wong, Yun Ka Wong, Gaizhen Liu, Meng Wang, Lei Sun, Federico Torta, Haipeng Sun, David Lawrence Silver, Chen Gao, Yibin Wang

**Affiliations:** Signature Research Programme in Cardiovascular and Metabolic Disorders, Duke-NUS Medical School, Singapore; National Heart Centre Singapore, Singapore; Singapore Lipidomics Incubator (SLING), Life Sciences Institute and Precision Medicine Translational Research Programme and Department of Biochemistry, YLL School of Medicine, National University of Singapore, Singapore; NHC Key Laboratory of Hormones and Development, Center for Cardiovascular Diseases, The Province and Ministry Co-sponsored Collaborative Innovation Center for Medical Epigenetics, Chu Hsien-I Memorial Hospital & Tianjin Institute of Endocrinology, Tianjin Medical University, Tianjin 300134, China; Key Laboratory of Tropical Translational Medicine of Ministry of Education, Hainan Academy of Medical Sciences/International Center for Aging and Cancer, Hainan Medical University, Haikou 571199, China; Department of Pharmacology and Systems Physiology, University of Cincinnati, Ohio, USA

## Abstract

Branched-chain amino acids (BCAAs) are essential amino acids and their homeostasis is tightly regulated by mostly mitochondrial localized catabolic activities where branched-chain α-keto acids (BCKAs) are the first-step downstream products. BCAAs and BCKAs are both readily detected in circulation, necessitating targeted transporters for their uptake across cellular and mitochondrial membranes. Although mitochondrial transporter for BCAA uptake has been identified, the molecular entity for BCKAs mitochondrial uptake remains unknown. In adult mice, we measured circulating BCAA and BCKAs under feeding and fasting status, and found circulating BCKAs were markedly elevated during fasting while BCAAs levels remained constant. By integrating genetic association analysis in human and multi-tissue nutrient-responsive transcriptomic profiling in mice, we identified SLC25A35 as a candidate mitochondrial transporter highly associated with BCKA homeostasis. SLC25A35 expression was highly enriched in brown adipocytes. Genetic inactivation of SLC25A35 in mice specifically blunted BCKA mitochondria uptake and downstream catabolism without significant impact on BCAA homeostasis, as demonstrated by targeted metabolic profiling and isotope tracing in intact animals, in isolated primary adipocytes and in purified mitochondria. SLC25A35 expression directly promoted mitochondrial BCKA uptake, with a preference for KIC and KMV over KIV. Importantly, SLC25A35 mediated BCKA uptake and catabolism was independent from SLC25A44 mediated BCAA mitochondrial transport, and the BCKA specificity for SLC35A35 was further evaluated by structure-guided identification and mutagenesis based validation for conserved potential BCKA interacting residues in SLC25A35 transmembrane domains. Functionally, SLC25A35 expression enhanced lipogenesis by supporting fatty acid incorporation of BCKA-derived carbon and lipogenic gene expression, independent from mitochondrial production of PEP. During cold exposure, SLC25A35-deficient mice exhibited reduced mitochondrial respiratory activity in brown adipose tissue, depletion of short- and long-chain acylcarnitine species, and attenuated cold-responsive metabolic signalling. Together, these findings identify SLC25A35 as a nutrient-responsive mitochondrial BCKA transporter that reveal a previously unrecognized BCKA specific contribution to adipocyte lipid metabolism and thermogenic adaptation.

## INTRODUCTION

In animals, systemic metabolic homeostasis is maintained despite dynamic shifts in the external nutrient environment during feeding and fasting^1,2^. For critical nutrients such as glucose and fatty acids, elaborate regulatory networks have been established to strategically produce, store and release these metabolites in response to feeding or starvation in order to maintain steady state circulating levels^3,4^. For essential amino acids with no *de novo* synthesis capacities in animal cells, their homeostasis regulations are mostly achieved through controlled degradation, i.e., catabolic fluxes^5^. Branched-chain amino acids (BCAAs) are essential amino acids with important functions beyond protein synthesis, including nutrient signaling and cellular homeostasis^6–8^. Indeed, recent studies have uncovered a significant impact of abnormal BCAA catabolism on human physiology and diseases, particularly for chronic disorders such as diabetes, fatty liver and heart failure^9–13^. However, our current understanding of BCAA regulation and related pathogenic mechanism remains very limited.

The current paradigm for BCAA homeostasis centers on the dedicated catabolic activity in mitochondria, particularly, the rate-limiting branched-chain α-keto acid dehydrogenase (BCKDH) complex ^14–17^. BCKDH activity is directly induced by the upstream substrates branched-chain α-keto acids (BCKAs)^18^. This positive feed-forward regulation is viewed as the cornerstone of BCAA homeostasis in response to the fluctuation in environmental BCAA levels. BCKA are the first downstream metabolites of BCAA and can be interconverted into BCAAs through reversible activities of the branched-chain amino acid aminotransferases (BCATs)^19,20^. Therefore, systemic BCAA and BCKA levels are expected to remain tightly coordinated across different nutrient conditions. Although this paradigm is well established at the biochemical level *in vitro*, it has not been directly tested in intact animals. In our current study, we analyzed steady-state plasma levels of BCAAs and BCKAs under feeding and starvation conditions and found that the circulating BCAA levels were unaffected, presumably due to a robust suppression of mitochondrial BCKDH activity due to lower levels of intra-mitochondria BCKA. To our surprise, our data showed the circulating BCKA levels were markedly elevated during starvation. This unexpected uncoupling between circulating BCKA and mitochondrial BCAA catabolic activity during fasting-feeding cycle challenges the current paradigm of a coupled BCAA/BCKA degradation in mitochondria regulated through BCKA mediated modulation of BCKDH activities, and implies the presence of a so-far unknown step for nutrient-sensitive BCKA specific entry into mitochondria

Mitochondria are essential organelles that play central roles in cellular metabolism and homeostasis^21^. The mitochondria-compartmented metabolic activities, such as BCAA catabolism, require well- orchestrated influx and efflux of metabolic substrates and products^22,23^. The solute carrier (SLC) 25 family members consist of mostly transporters located at the mitochondrial inner or outer membrane^24–26^. While the substrates for many SLC25A family members have been identified, there are still numerous transporters with unknown substrates^27^. Identifying their substrates for these so-called “orphan” transporters would be critical to unlock the molecular network governing nutrient utilization and mobilization. In the current study, we systematically analyzed the expression profiles of SLC25A family members in adult mice across multiple tissues under feeding and fasting conditions. Among the nutrient regulated SLC25A transporters, the SLC25A35 was ranked top for nutrient responsiveness with a robust down-regulation under fasting conditions and upregulation following refeeding across different tissues in adult mice. Systems-based analysis also identified a significant association between SLC25A35 and BCKA metabolism in humans. Targeted metabolomic and flux analyses in the wildtype and SLC25A35-deficient mice demonstrated that SLC25A35 expression was necessary for normal mitochondrial BCKA uptake and downstream metabolites without measurable impact on BCAA uptake or metabolism. SLC25A35 inactivation led to deranged systemic homeostasis of BCKA in response to fasting and refeeding, also without significant impact on BCAA. Remarkably, SLC25A35 was detected to be highly enriched in brown adipose tissue, and its expression was necessary for cold-induced adaptation, thermogenesis, and fatty acid metabolism in brown adipose tissue. Therefore, we propose that SLC25A35 functions as a BCKA mitochondrial transporter with a substrate-specific and nutrient status-specific role in BCKA uptake. This finding uncovered a new independent regulatory circuit in BCAA/BCKA catabolism through a mitochondrial transporter dedicated to BCKA catabolism. The functional impact from SLC25A35 deficiency in mice underscores the previously unappreciated role of BCKA in adipose metabolism and thermogenic activity.

## RESULTS

### Nutrient-responsive profiling nominates SLC25A35 as a candidate mitochondrial BCKA transporter

One of the key hallmarks of metabolic homeostasis is to maintain stable internal levels under a dynamically changing environment. Branched-chain amino acids (BCAAs) are essential amino acids that cannot be synthesized *de novo* in animal cells^28^. Therefore, the intra-tissue and systemic free BCAA can only be sourced from external food uptake or internal recycling from degraded proteins^29^. However, how BCAA homeostasis is maintained under different nutrient statuses remains poorly understood. In order to investigate BCAA homeostasis *in vivo,* we measured circulating BCAA levels under both feeding and starvation conditions and found they were not affected by the feeding status (Figure 1A). In contrast, the circulating levels of branched-chain α-keto acids (BCKA), the first downstream catabolic products of BCAA, were markedly elevated under fasting conditions (Figure 1B). This is unexpected uncoupling between the systemic levels of BCKA vs. BCAA during starvation indicated that mitochondrial uptake and degradation of BCKA vs. BCAA involve two independent regulatory circuits.

**Figure 1.**
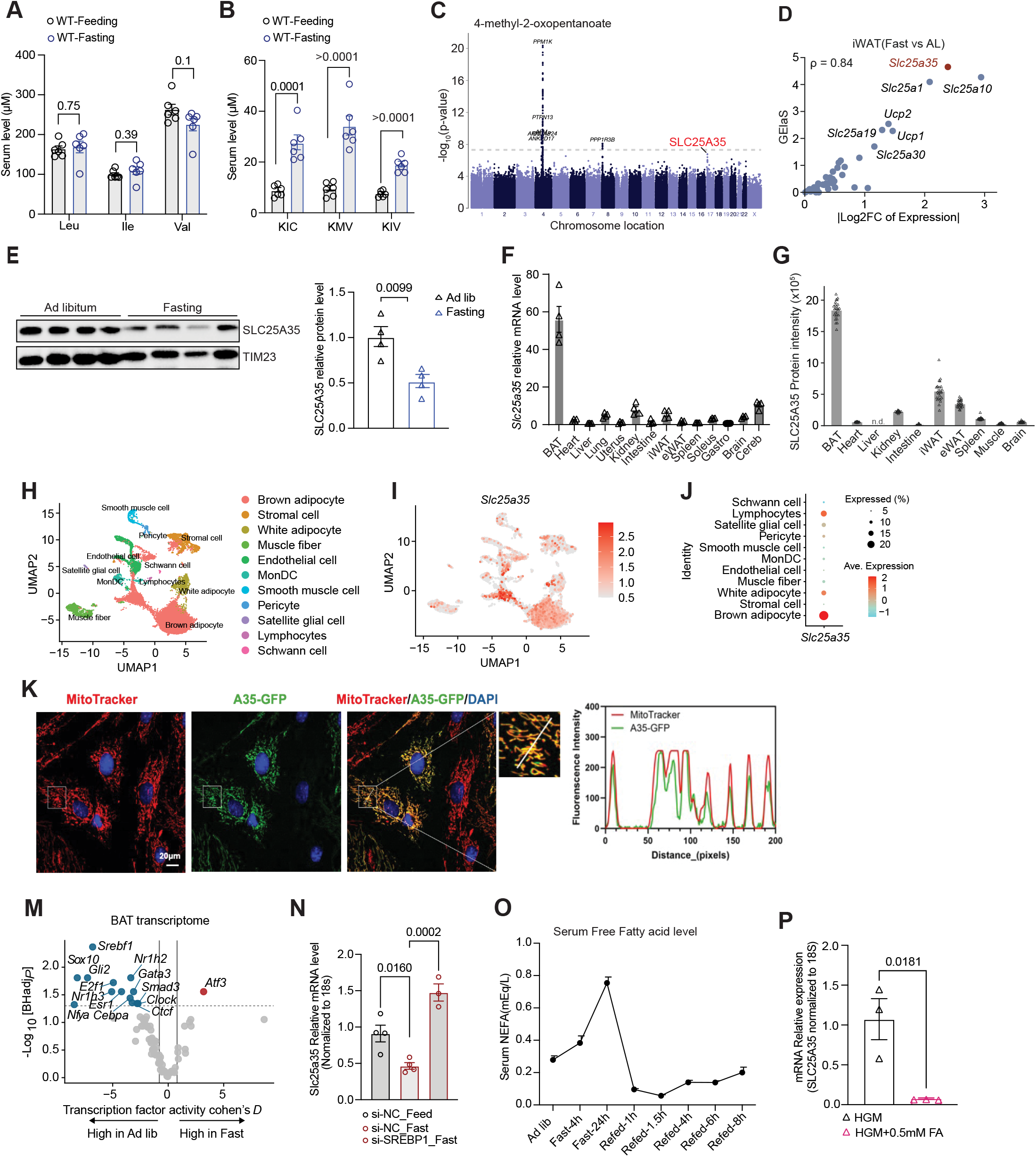
Nutrient-responsive profiling identifies SLC25A35 as a candidate mitochondrial BCKA transporter. (A) Serum levels of branched-chain amino acid (BCAA) in WT mice under fed and overnight-fasted conditions (n = 6). (B) Serum levels of branched-chain keto acid (BCKA) in WT mice under fed and overnight-fasted conditions (n = 6). (C) Manhattan plot of the associations between genetic variants and plasma 4-methyl-2-oxopentanoate (KIC) in the METSIM cohort (https://pheweb.org/metsim-metab). The signal at the *SLC25A35* locus is labeled in red. (D) Gene elasticity score (GElaS) against the absolute log_2_ fold change on fasting versus ad libitum for SLC25 families analyzed in inguinal white adipose tissue (iWAT) of young mice. (E) Immunoblot analysis of mitochondrial SLC25A35 protein isolated from brown adipose tissue (BAT) of mice under ad libitum and fasting conditions (left panel). Quantification of SLC25A35 levels normalized to the inner mitochondrial membrane marker TIM23 is shown in the right panel (n = 4). (F) *Slc25a35* mRNA expression across the indicated organs under ad libitum feeding condition (n = 4). (G) SLC25A35 protein abundance (DIA–MS intensity) in ten tissues of male WT mice from a published multi-organ proteome atlas^35^. Bars show the mean, whiskers ± SEM and triangles individual mice (n = 24). (H) Uniform manifold approximation and projection (UMAP) of single-cell transcriptomes of brown adipose tissue, colored by cell type. Data reanalyzed from Behrens *et al*^45^. (I) *Slc25a35* expression in mouse brown adipose tissue in UMAP. (J) Dot plot of *Slc25a35* expression across cell types showed *Slc25a35* is mainly expressed in brown adipocyte. (K) Fluorescence images of pre-adipocytes expressing GFP-tagged human SLC25A35 (green) and stained with MitoTracker (red) and DAPI (blue). Scale bar, 20 µm. (L) Fluorescence intensity profiles of MitoTracker (red) and A35-GFP (green) along the white line in the magnified region in right panel. (M) Volcano plot of the differences in transcription factor activity between the ad libitum and fasted states in the BAT transcriptome. (N) *Slc25a35* mRNA levels in cultured brown pre-adipocytes transfected with control siRNA (si-NC) or SREBP1 siRNA (si-SREBP1) and cultured under fed or fasting conditions (n = 3-4). (O) Serum free fatty acid (NEFA) levels in wild-type mice under ad libitum, fasting and refeeding conditions at the indicated time points (n = 10). (P) Relative *Slc25a35* mRNA levels in AML12 cells cultured in high-glucose medium without 10% FBS in the absence or presence of 0.5 mM free fatty acids (palmitic acid and oleic acid) (n = 3). Data are presented as mean ± SEM. P values in panels (A and B) were determined by multiple unpaired t tests with false discovery rate (FDR) correction. P values in panels (E and P) were determined by unpaired two-tailed Student’s t tests. P values in panel (N) were determined by one-way ANOVA followed by multiple comparisons test.

To identify candidate mitochondrial BCKA transporters, we interrogated human genetic associations with circulating KIC levels using PheWeb^30,31^. The strongest association mapped to the *PPM1K* locus. PPM1K is a mitochondrial phosphatase that activates the BCKDH complex and a well-established regulator of BCKA oxidation. Notably, we also identified a suggestive association near genetic locus of human SLC25A35 gene (rs199925223, β = 0.81, SE = 0.15, P = 1.7 × 10̅T) (Figure 1C). SLC25A35 is a new characterized member of the mitochondrial SLC25 family, reported to have transport activity for mitochondrial produced phosphoenolpyruvate (PEP). Because metabolites transfer across mitochondrial membranes are carried out by dedicated mitochondrial transporters, many of which are encoded by SLC25 family genes^32^. We next examined the nutrient-dependent regulation of the transporter family. We performed transcriptomic analyses in four major metabolic organs, including inguinal white adipose tissue (iWAT), epididymal white adipose tissue (eWAT), skeletal muscle (SKM), and liver, under ad libitum (AL), 16 hour fasting (F), and followed by 4 hour refeeding (R) conditions in 2-month-old healthy male mice^33^. Based on the changes of mRNA levels during the AL-F-R cycle, we ranked all SLC25 family genes according to their nutrient-dependent gene elasticity using the previously described Gene Elasticity Score (GEIaS) as described previously (Figure 1D and S1A)^33^. and many of these genes have known substrates such as SLC25A1 (for citrate)^34,35^, SLC25A10 (for dicarboxylate acid)^35^, SLC25A19 (for thiamine pyrophosphate)^36^, and SLC25A7 (for proton)^37,38^, each with established roles in lipid and carbohydrate metabolism. Together, the human genetic association and nutrient-responsive expression profile nominated SLC25A35 as a candidate mitochondrial BCKA transporter for functional investigation.

Using an independent cohort of male mice, we further confirmed that *Slc25a35* mRNA expression was significantly reduced under fasting conditions but re-activated upon refeeding across multiple metabolically active organs, such as brown adipose tissue (BAT), iWAT, SKM, liver and heart (Figure S2A and S2B). SLC25A35 protein abundance was reduced in brown adipose tissue during fasting (Figure 1E). At both mRNA and protein levels^39^, SLC25A35 was highly enriched in fat tissues, particularly brown adipose tissues in adult mice (Figure 1F and 1G, S2C). To define the cellular distribution of *Slc25a35* in mouse BAT, we reanalyzed a published single-nucleus RNA-sequencing dataset derived from mouse brown adipose tissue. UMAP analysis identified adipocyte and non-adipocyte populations (Figure 1H). Feature and dot-plot analyses showed that *Slc25a35* was predominantly expressed in adipocytes, with the highest expression in brown adipocytes (Figure 1I and 1J). As expected, the endogenous SLC25A35 was enriched in the mitochondrial fraction from brown adipose tissue (Figure S2D) and the human SLC25A35 protein expressed in preadipocyte was localized to mitochondria based on immunofluorescent microscopy (Figure 1K and 1L). In cultured brown preadipocytes and AML12 hepatocytes, serum starvation markedly reduced the expression of SLC25A35 at mRNA and protein levels (Figure S2E-S2H), whereas serum replenishment markedly increased SLC25A35 expression. These results indicate that SLC25A35 is dynamically regulated in response to nutrient environment, a hallmark of a nutrient sensitive regulator. Transcription factor activity analysis of BAT transcriptomes from different feeding conditions identified *Srebf1* mediated transcription activity was reduced in fasting compared to ad libitum feeding conditions (Figure 1M, S2I and S2J). in cultured adipocytes, SREBP1 knockdown significantly blunted starvation induced *Slc25a35* expression (Figure 1N and S2K). These results suggest that SREBP1 may serve as upstream regulator *Slc25a35* expression during fasting. Consistent with the role of lipid sensitive SREBP1 in SLC25A35 regulation, we found circulating free fatty acid levels increased markedly during fasting and rapidly declined following refeeding (Figure 1O). In cultured brown preadipocytes, high-glucose treatment did not alter Slc25a35 mRNA, SLC25A35 protein abundance, or promoter activity (Figure S2L-S2N). In contrast, fatty acid treatment significantly suppressed SLC25A35 mRNA and protein expression and markedly reduced SLC25A35 promoter activity, indicating that lipid-derived signals are responsible for SLC25A35 suppression during fasting state (Figure 1P, and S2O-S2Q). These findings indicate that elevated fatty acid availability is sufficient to suppress SLC25A35 expression and may contribute to its downregulation during fasting.

### SLC25A35 is associated with and necessary for BCKA homeostasis

To further investigate whether SLC25A35 functions as a BCKA transporter, we conducted an integrated analysis, including gene–module association determination (G-MAD), genome-wide association studies (GWAS) and metabolomics (Figure 2A). Based on G-MAD analysis using 14 publicly available gene expression datasets from mouse adipose tissue^40,41^, *slc25a35* gene expression was significantly associated with genes enriched in the KEGG pathways for TCA cycle and branched-chain amino acid metabolism (Figure 2B). We also performed association analysis between the genomic variants of SLC25A35 and plasma metabolites in humans using the METSIM (Metabolic Syndrome in Man) platform from a dataset of 6,136 Finnish men^30,31^. This analysis identified 4-methyl-2-oxopentanoate, the branched-chain keto acid product from L-leucine, with a significant association (p=1.7e-7) with a minor allele of SLC25A35 gene (G262A; rs199925223) (Figure 2C). Importantly, the same allele also showed trends to be associated with several other metabolites related to BCKAs, including 3-methyl-2-oxovalerate (BCKA product from isoleucine), α-hydroxy-isovalerate (derivative of BCKA product from leucine), 2-ethyl-3-hydroxypropionic acid and 3-hydroxyisobutyrate (derivatives of BCKA product from valine). Therefore, these unbiased systems-based analyses point to a significant association between SLC25A35 and BCAA/BCKA metabolism in humans and mice.

**Figure 2.**
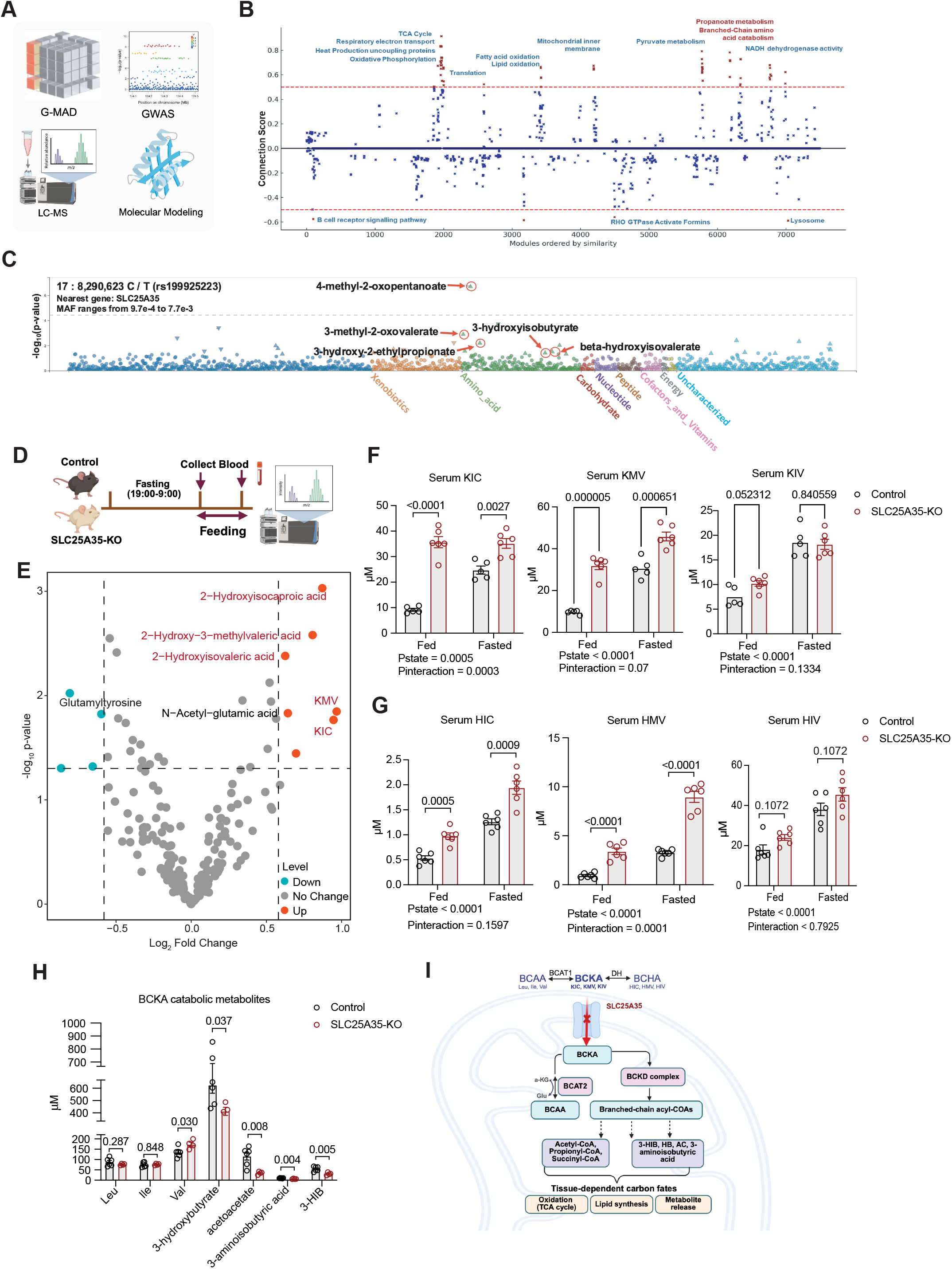
SLC25A35 is associated with and required for systemic BCKA homeostasis. (A) Schematic overview of the integrative analytical strategy used to identify the metabolic substrate and function of SLC25A35, combining gene–module association determination (G-MAD), genome-wide association studies (GWAS), LC–MS–based metabolomics, and molecular modeling. (B) Gene–module association analysis of *Slc25a35* using mouse adipose tissue expression datasets. Dashed lines indicate the correlation threshold (r = 0.5). Gene modules significantly associated with *Slc25a35* are highlighted in red (https://systems-genetics.org/gmad). (C) Stacked PheWeb plots showing associations between the SLC25A35 missense variant rs199925223 and BCKA-related metabolites in the METSIM cohort (https://pheweb.org/metsim-metab/variant/17-8290623-C-T). (D) Experimental schematic illustrating serum collection from control and SLC25A35-KO mice after overnight fasting (7pm-7am) and after refeeding. (E) Volcano plot comparing serum metabolite levels between control and SLC25A35-KO mice under refeeding condition. (n = 6). (F) Serum levels of the KIC, KMV and KIV in control and SLC25A35-KO mice in the fed (2 hours after refeeding) and overnight-fasted states (n = 5–6). (G) Serum levels of the BCKA-derived cytosolic metabolites 2-hydroxyisocaproate (HIC), 2-hydroxy-3-methylvaleric acid (HMV) and 2-hydroxyisovaleric acid (HIV) in control and SLC25A35-KO mice in the fed (2 hours after refeeding) and overnight-fasted states (n = 6). (H) Serum levels of BCAAs (Leu, Ile, Val) and BCKA catabolic metabolites (3-hydroxybutyrate, acetoacetate, 3-aminoisobutyric acid and 3-hydroxyisobutyrate [3-HIB]) in control and SLC25A35-KO mice from the BCKA tolerance-test cohort (n = 3-6). (I) Schematic representation of BCKA metabolism, illustrating SLC25A35-dependent mitochondrial uptake of BCKA, their transamination by BCAT2 and catabolism by the BCKD complex, the tissue- dependent fates of BCKA-derived carbon, and the reversible reduction of BCKA to their reduced metabolites in the cytosol. Data are presented as mean ± SEM. P values in panels (F and G) were determined by two-way ANOVA followed by multiple comparisons test. P values in panel (H) were determined by unpaired two-tailed Student’s t tests.

To define the physiological role of SLC25A35, we deleted exon 2 to 5 of the mouse *slc25a35* gene by breeding *Slc25a35*^flox/flox^ mice with germline cre-expressing mice (Zp3-cre) to generate SLC25A35 global deletion mice (SLC25A35-KO; Figure S3A). The status of SLC25A35 deficiency in the SLC25A35-KO mice was confirmed by a complete loss of mRNA and protein across multiple tissues compared to the wildtype controls (Figure S3B and S3C). The SLC25A35-KO mice developed normally to adulthood compared to the wildtype controls, based on gross morphometric and metabolic phenotype at 8 weeks in male mice, including body weight, fat mass, lean mass, organ weight, food intake, and glucose clearance (Figure S3D-S3G). However, untargeted metabolomic analysis using mass spectrometry in plasma samples from SLC25A35-KO and control mice identified significant shifts in metabolic profiles between the two genotypes under the feeding and fasting conditions based on principal component analysis (PCA) (Figure 2D, S3H and S3I, S3K and S3L). From volcano and heatmap analyses, circulating KIC and KMV levels as well as their cytosolic metabolic by-products, including 2-hydroxyisocaproate (HIC), 2-hydroxy-3-methylvaleric acid (HMV), and 2-hydroxyisovaleric acid (HIV), were significantly elevated in the SLC25A35-KO mice compared to the wildtype controls (Figure 2E, S3J, S3M and S3N). In contrast, no differences were identified for BCAAs. These results are consistent with impaired BCKA handling in the absence of SLC25A35 and support a role for SLC25A35 in the mitochondrial compartmentalization of BCKA metabolism.

Extending our findings from untargeted metabolomics, we performed additional targeted MS analysis on serum samples collected after overnight fasting and 2 hours of refeeding in mice (Figure 2D). Fasting markedly increased circulating BCKA levels in the wildtype mice as described earlier (Figure 2F). In contrast, SLC25A35 deletion led to an accumulation of circulating BCKAs in both fasting and refeeding conditions compared to the controls, particularly for KIC and KMV levels (Figure 2F). Consequently, fasting triggered BCKA elevation was attenuated significantly for KIC based on genotype x nutritional-state interaction (p = 0.0003), whereas the interactions for KMV and KIV did not reach statistical significance. Consistent with impaired mitochondrial clearance of BCKA in the SLC25A35-KO mice, the cytosolically derived BCKA downstream metabolites, including 2-hydroxyisocaproate (HIC), 2-hydroxy-3-methylvaleric acid (HMV), and 2-hydroxyisovaleric acid (HIV), were significantly elevated in circulation (Figure 2G). Given impaired BCKAs uptake in SLC25A35-KO mice, we next examined whether their mitochondrial localized downstream catabolism was altered in circulation. Leucine and isoleucine levels remained unchanged, whereas valine was modestly elevated in the KO mice (Figure 2H). Despite unchanged leucine abundance, its downstream ketogenic products, acetoacetate and 3-hydroxybutyrate, were significantly reduced; the valine-derived metabolite 3-HIB and 3-aminoisobutyric acid were also decreased (Figure 2H). Together with the accumulation of circulating BCKAs, these changes are consistent with impaired mitochondrial BCKA utilization and remodelling of downstream BCAA catabolism following SLC25A35 inactivation (Figure 2I).

### SLC25A35 is required for mitochondrial BCKA import and catabolism

To unambiguously determine whether SLC25A35 is specifically involved in BCKA mitochondrial transport and clearance, we conducted stable isotope tracing study in SLC25A35-KO and control mice by analyzing the serum samples collected at 0, 3, 8, 15, and 25 minutes following the administration of a single bolus intraperitoneal injection of ^13^C-KIC (M+6) and unlabeled ^12^C-KMV and ^12^C-KIV as illustrated (Figure 3A and S4A). In the wildtype control mice, the labeled KIC reached peak values within 3 minutes and was cleared after 15 minutes (Figure 3B). Concurrently, the circulating leucine produced from the labeled KIC (via cytosolic and mitochondrial BCAT activities) also reached its peak and then subsided within the same time frame (Figure 3C). These changes imply a robust catabolic flux and fast BCKA to BCAA conversion at the systemic level in mice. Loss of SLC25A35 markedly impaired the systemic clearance of circulating ^13^C-KIC (M+6) and significantly reduced the generation of ^13^C-Leu (M+6) compared with control mice (Figure 3B and 3C). In parallel, the cytosolic reduction product ^13^C-HIC (M+6), derived from ^13^C-KIC (M+6), was accumulated to higher levels in SLC25A35-KO mice (Figure 3D). In SLC25A35-KO mice, however, the endogenous KIC (M+0) and KMV (M+0), but not KIV (M+0) showed significantly higher levels across all time points post-injection (Figure S4B). Consistently, the unlabeled cytosolic BCKA-derived metabolites HIC (M+0) and HMV (M+0) also were accumulated to higher levels in SLC25A35-KO mice compared with controls (Figure S4C). Importantly, the endogenous BCAA (Leu (M+0), Ile (M+0) and Val (M+0)) were not altered between SLC25A35-KO and control mice (Figure S4D). Together, these isotope-resolved flux analyses demonstrate that SLC25A35 is required for efficient uptake and metabolic flux of BCKA *in vivo*, supporting its role as a mitochondrial transporter of BCKA (Figure S4E).

**Figure 3.**
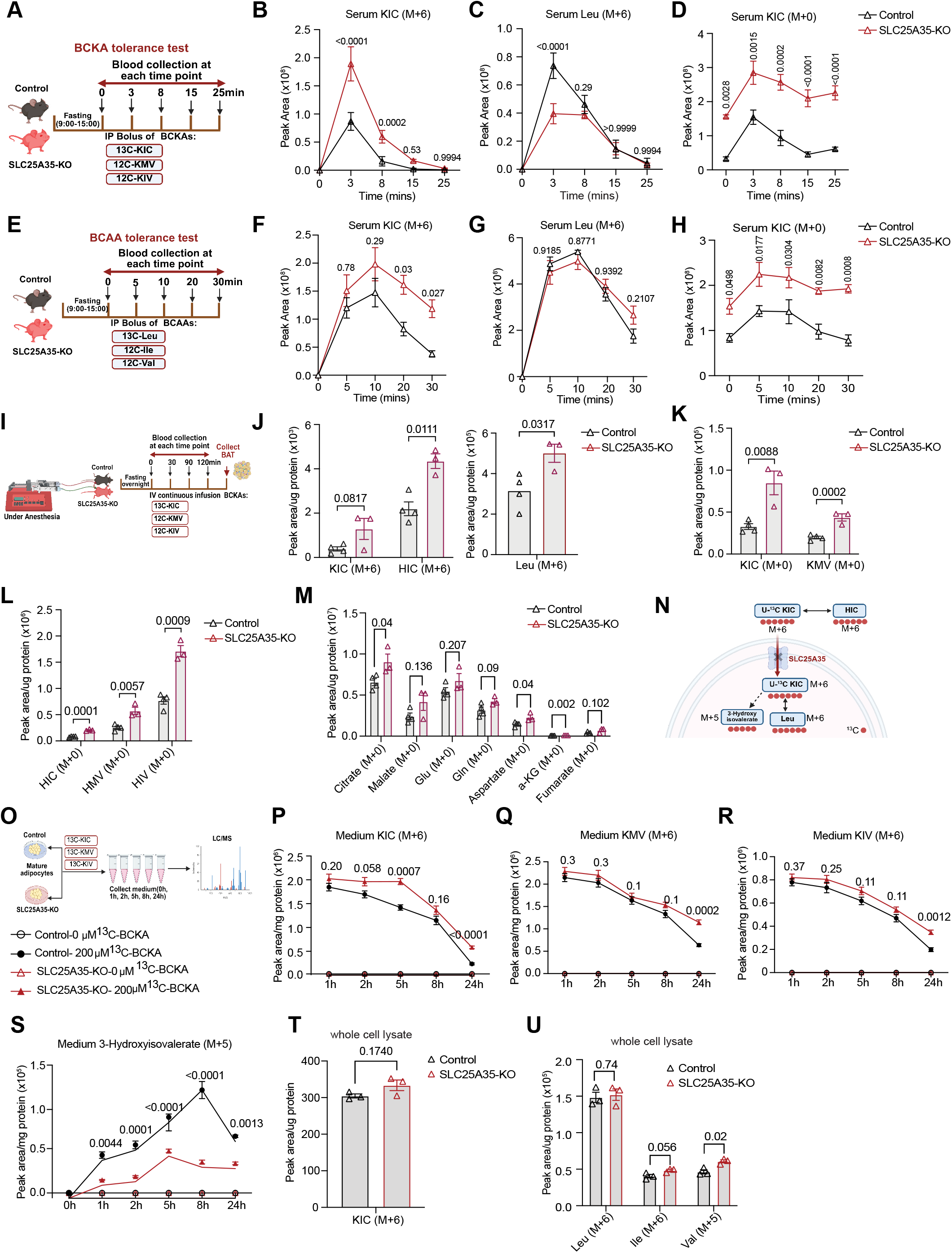
SLC25A35 is required for mitochondrial BCKA uptake and catabolism. (A) Experimental schematic of the BCKA tolerance test, in which control and SLC25A35-KO mice received an intraperitoneal (IP) bolus of uniformly ^13^C-labeled KIC together with unlabeled KMV and KIV, followed by serum collection at 0, 3, 8, 15, and 25 min for metabolite analysis. (B–D) Serum levels of ^13^C-KIC (M+6) (B), ^13^C-KIC–derived Leu (M+6) (C), and unlabeled KIC (M+0) (D) measured at the indicated time points in control and SLC25A35-KO mice (n = 4). (E) Experimental schematic of the BCAA tolerance test, in which control and SLC25A35-KO mice received an IP bolus of uniformly ^13^C-labeled leucine together with unlabeled isoleucine and valine, followed by serum collection at 0, 5, 10, 20, and 30 min for metabolite analysis. (F–H) Serum levels of ^13^C-KIC (M+6) (F), ^13^C-Leu (M+6) (G) and unlabeled KIC (M+0) (H) measured at the indicated time points in control and SLC25A35-KO mice after the ^13^C-Leu bolus (n = 3). (I) Experimental schematic of *in vivo* steady-state intravenous infusion of uniformly ^13^C-labeled BCKA in control and SLC25A35-KO mice, with serum collected at 0, 30, 90, and 120 min and BAT collected at the end of the infusion for metabolite analysis. (J) BAT levels of ^13^C-KIC (M+6) and ^13^C-HIC (M+6) (left) and of ^13^C-Leu (M+6) (right) at the end of the infusion (n = 3-4). (K) BAT levels of unlabeled KIC (M+0) and KMV (M+0) during steady-state infusion (n=3-4). (L) BAT levels of the unlabeled cytosol-derived BCKA metabolites HIC (M+0), HMV (M+0) and HIV (M+0) during steady-state infusion (n = 3-4). (M) BAT levels of citrate, malate, glutamate (Glu), glutamine (Gln), aspartate, α-ketoglutarate (α-KG) and fumarate (M+0) during steady-state infusion (n = 3-4). (N) Schematic illustrating the fate of uniformly ^13^C-labeled KIC: mitochondrial uptake through SLC25A35, transamination to ^13^C-Leu (M+6), catabolism to ^13^C-3-hydroxyisovalerate (M+5), and cytosolic reduction to ^13^C-HIC (M+6). (O) Experimental schematic of medium collection, illustrating sampling at 1, 2, 5, 8, and 24 h from mature adipocytes differentiated from control and SLC25A35-KO preadipocytes. (P–S) Medium levels of ^13^C-labeled BCKA, including ^13^C-KIC (M+6) (P), ^13^C-KMV (M+6) (Q), ^13^C-KIV (M+5) (R), and ^13^C-KIC–derived metabolite ^13^C-3-hydroxyisovalerate (M+5) (S), measured in mature adipocytes differentiated from control and SLC25A35-KO preadipocytes with or without 200 µM ^13^C-BCKA treatment (n = 4). (T–U) Peak areas of intracellular ^13^C-labeled metabolites, normalized to total cellular protein content, measured in whole-cell extracts from control and SLC25A35-KO mature adipocytes after 24 h incubation with uniformly ^13^C-labeled BCKA, including ^13^C-KIC (M+6) (T) and ^13^C-labeled BCAA (U) (n = 3). Data are presented as mean ± SEM. P values in panels (B–D, F–H and P–S) were determined by two-way ANOVA followed by Tukey’s multiple comparisons test. P values in panels (J, L and T) were determined by unpaired two-tailed Student’s t tests. P values in panels (K, M and U) were determined by multiple unpaired t tests with false discovery rate (FDR) correction.

In contrast to the ^13^C-KIC (M+6) tracing results (Figure 3E and S4F), no significant differences in labeled Leucine accumulation were observed between the SLC25A35-KO mice and the wildtype controls when ^13^C-Leu (M+6) was used as tracer (Figure 3G). Intriguingly, the first catabolic product ^13^C-KIC (M+6) was significantly higher in the SLC25A35-KO mice after the peak accumulation time (Figure 3F), again supporting the necessary role of SLC25A35 for BCKA influx but not for their efflux across mitochondria. The cytosolic reduction product of ^13^C-KIC (M+6), ^13^C-HIC (M+6) exhibited a modest increase at later time points following BCAA stimulation (Figure 3H). The unlabeled BCKA, including KIC (M+0) and KMV (M+0), but not KIV (M+0), and the reduction product HIC (M+0) remained highly elevated in SLC25A35-KO mice compared to WT mice (Figure S4G and S4H). There were also no significant differences in unlabeled BCAA, Ile (M+0), and Val (M+0), between the two groups (Figure S4I), highlighting the specificity of SLC25A35 for BCKA transport instead of BCAA (Figure S4J).

To further assess BCKA clearance in vivo, continuing administration of ^13^C-KIC (M+6), ^13^C-KMV (M+6), and ^13^C-KIV (M+5) was performed in anesthetized SLC25A35-KO and control mice for 2 hours. Blood samples were collected at 0, 30, 90, and 120 minutes and analyzed by LC-MS (Figure S5A). The results showed a progressively higher accumulation of labeled and unlabeled KIC and KMV, but not KIV, in the SLC25A35-KO mice compared to the control mice (Figure S5B). Consistent with impaired mitochondrial uptake, the accumulation of labeled cytosolic metabolites, including labeled and unlabeled HMV but not HIC, were significantly elevated in the SLC25A35-KO plasma compared to the controls (Figure S5C). Once again, no changes in labeled and unlabeled BCAA levels were affected by SLC25A35 inactivation (Figure S5D). These steady-state infusion data indicate that SLC25A35 preferentially mediates mitochondrial uptake of KIC and KMV, with limited contribution to KIV transport, without affecting systemic BCAA metabolism under physiological conditions.

To determine how SLC25A35 deficiency affects BCKA metabolism within BAT, We further measured tissue metabolites in BAT under ^13^C-KIC (M+6) infusion (Figure 3I). Following infusion, BAT levels of ^13^C-KIC (M+6) showed a trend of increase in the SLC25A35-KO mice compared with controls (Figure 3J). While ^13^C-Leu (M+6) derived from ^13^C-KIC were increased, whereas its cytosolic derived metabolite ^13^C-HIC (M+6) was markedly accumulated in the SLC25A35-KO mice compared with controls (Figure 3J). Consistent with the tracer-based observations, unlabelled KIC (M+0) and KMV (M+0), exhibited significant accumulation in BAT from SLC25A35-KO mice (Figure 3K). Again, the cytosol-derived BCKA reduction products—including HIC, HMV and HIV (M+0) —were significantly elevated in the BAT tissue of SLC25A35-KO mice compared with controls (Figure 3L). These findings indicate that SLC25A35 loss promotes intracellular BCKA accumulation and diverts BCKA toward cytosolic reduction pathways, consistent with impaired mitochondrial BCKA uptake. SLC25A35 deficiency was also associated with remodeling of central carbon metabolism in BAT. Citrate and aspartate were significantly elevated, whereas a-ketoglutarate was reduced in KO BAT (Figure 3M). Malate, glutamine and fumarate showed nonsignificant upward trends, while glutamate remained unchanged. These changes suggest that impaired mitochondrial BCKA uptake alters the coupling BCKA metabolism, amino acid transamination and the TCA cycle (Figure 3N).

Given the enrichment of SLC25A35 in brown adipocytes, we next examined BCKA uptake and metabolism in differentiated SLC25A35-KO and control adipocytes. Fully differentiated adipocytes were subsequently treated with 100 µM uniformly ^13^C-labeled BCKA, followed by time-resolved metabolomic analysis (Figure 3O and S5E). Extracellular labeled KIC, KMV, and KIV accumulated to significantly higher levels over time in SLC25A35-KO adipocytes than in control cells, consistent with impaired cellular BCKA uptake or clearance (Figures 3P-3R). Conversely, the ^13^C-KIC-derived catabolic product 3-hydroxyisovalerate (M+5) was markedly reduced in the medium of KO adipocytes (Figure 3S). Unlabeled KIC, KMV, and KIV showed similar time-dependent accumulation, accompanied by reduced production of unlabeled 3-hydroxyisovalerate (Figures S5F-S5I). In contrast, labeled leucine (M+6), isoleucine (M+6), and valine (M+5) did not differ between the two groups, indicating that BCAT-mediated BCKA-to-BCAA reamination remained intact (Figures S5J-S5L). These *in vitro* findings recapitulate the *in vivo* metabolic phenotype and further supported a direct role for SLC25A35 in adipocyte BCKA uptake and metabolism, rather than in BCAA transport. In adipocytes, the total concentrations of ^13^C-KIC (M+6), Leu (M+6), Ile (M+6), and Val (M+5) were the same in the SLC25A35 KO and wildtype control cells following labelled BCKA incubation (Figure 3T and 3U). This suggests that SLC25A35 loss alters BCKA uptake and metabolic flux without substantially changing the endpoint intracellular pools of labeled BCKA or BCAA.

### SLC25A35 restores systemic BCKA clearance and directly promotes mitochondrial BCKA uptake

Having established that SLC25A35 deficiency impairs BCKA clearance and promotes accumulation of cytosolic BCKA-derived metabolites, we next determined whether ectopic expression of SLC25A35 was sufficient to restore systemic BCKA utilization. An AAV9 vector expressing SLC25A35 under the control of the CMV promoter was administered in the SLC25A35-KO mice in order to drive SLC25A35 expression across multiple tissues. Following a single intraperitoneal bolus of BCKAs injection (Figure S6A), circulating levels of KIC, KMV, and KIV were significantly lower in the KO mice received AAV9-CMV-*Slc25a35*than the untreated KO mice (Figure S6C). Their corresponding cytosolic reduction products—HIC, HMV, and HIV—were also reduced following SLC25A35 re-expression (Figure S6D). Circulating leucine, isoleucine, and valine were similarly decreased (Figure S6E), consistent with the expected activation of the rate-limiting BCKDH complex by elevated mitochondrial influx of BCKA..

Liver has little BCAT2 activity so it is not a significant organ to catabolise BCAA directly yet may serve as a major organ for systemic BCKA clearance. However, SLC25A35 expression in liver is very low comparing to fat tissues. By administration of an AAV8 vector expressing SLC25A35 under albumin promoter (AAV8-albumin-SLC25A35), we achieved high level of SLC25A35 expression specifically in liver in the SLC25A35 KO mice Figure 4A). Liver-specific SLC25A35 expression significantly reduced circulating KIC in the KO mice under feeding condition but not during fasting, while circulating KMV levels were markedly reduced under both fed and fasted conditions.. In contrast, KIV concentrations did not differ significantly among the three groups under either nutritional conditions (Figure 4B). Interestingly, the elevated serum levels of HIC, HMV, and HIV due to *Slc25a35* deficiency were not corrected by hepatic expression of SLC25A35 (Figure 4C). In the meantime, circulating leucine, isoleucine, and valine were largely unchanged among all the experimental groups under both fed and fasted conditions, except for modest reduction in isoleucine in fasted mice with hepatic SLC25A35 expression (Figure 4D). Together, these findings indicate that hepatic SLC25A35 expression preferentially improves KIC and KMV clearance without broadly altering systemic BCAA pools or fully normalizing their cytosolic reduction products.

**Figure 4.**
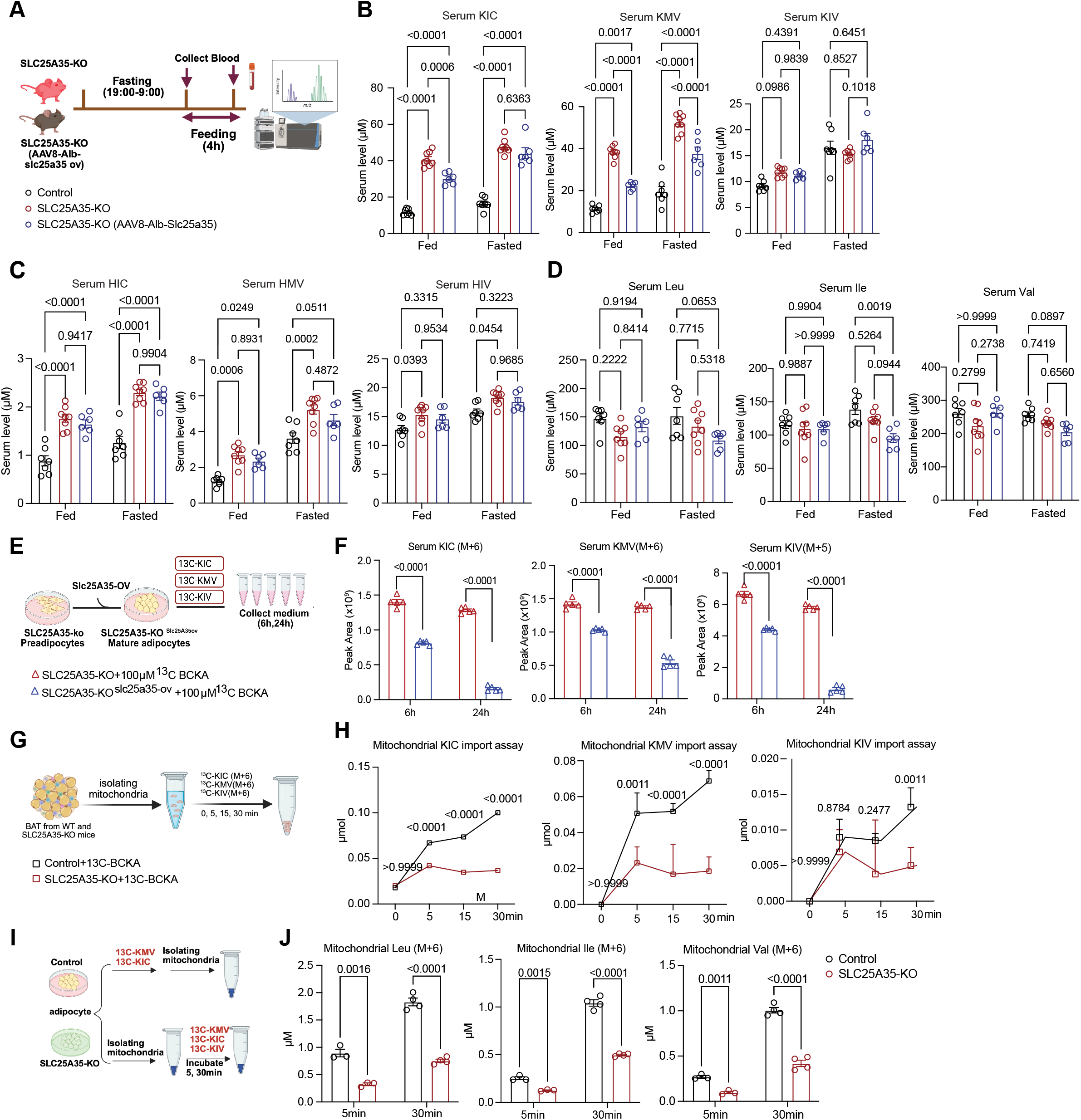
SLC25A35 re-expression restores systemic BCKA clearance and mitochondrial uptake. (A) Experimental schematic illustrating serum collection from control, SLC25A35-KO and SLC25A35-KO mice with liver-specific re-expression of SLC25A35 (AAV8-Alb-*Slc25a35*) after overnight fasting (7PM-7AM) and after 4 hours of feeding. (B) Serum levels of KIC, KMV and KIV in the fed and fasted states in control, SLC25A35-KO and SLC25A35-KO (AAV8-Alb-*Slc25a35*) mice (n = 6-8). (C) Serum levels of HIC, HMV and HIV in the mice described in (B) (n = 6-8). (D) Serum levels of Leu, Ile and Val in the mice described in (B) (n = 6-8). (E) Experimental schematic of the in vitro rescue experiment, in which SLC25A35-KO preadipocytes and SLC25A35-KO preadipocytes re-expressing SLC25A35 (SLC25A35-^KOSlc25a35-ov^) were differentiated into mature adipocytes and incubated with uniformly ^13^C-labeled KIC, KMV and KIV; medium was collected after 6 h and 24 h. (F) Medium levels of ^13^C-KIC (M+6), ^13^C-KMV (M+6) and ^13^C-KIV (M+5) measured in SLC25A35-KO and SLC25A35-rescued mature adipocytes after 6- and 24-hours incubation with 100 µM uniformly ^13^C-labeled BCKA (n = 5). (G) Experimental schematic of the mitochondrial uptake assay, in which mitochondria isolated from BAT of wild-type and SLC25A35-KO mice were incubated with ^13^C-KIC (M+6), ^13^C-KMV (M+6) and ^13^C-KIV (M+5) and sampled at 0, 5, 15 and 30 min for downstream measurements. (H) Time course of the import of ^13^C-KIC, ^13^C-KMV and ^13^C-KIV into BAT mitochondria from control and SLC25A35-KO mice (n = 4). (I) Experimental schematic of the mitochondrial uptake assay in mature adipocytes, in which mitochondria isolated from control and SLC25A35-KO adipocytes were incubated with uniformly ^13^C-labeled BCKA for 5 and 30 min. (J) Mitochondrial levels of ^13^C-Leu (M+6) (J), ^13^C-Ile (M+6) (K) and ^13^C-Val (M+5) (L) after 5- and 30- min incubation of isolated adipocyte mitochondria with ^13^C-BCKA in control and SLC25A35-KO samples (n = 3-4). Data are presented as mean ± SEM. P values in panels (B–D) were determined by two-way ANOVA followed by Tukey’s multiple comparisons test. P values in panel (F) were determined by one-way ANOVA followed by Dunnett’s multiple comparisons test. P values in panel (H) were determined by two-way ANOVA followed by Tukey’s multiple comparisons test. P values in panels (J–L) were determined by multiple unpaired t tests with false discovery rate (FDR) correction.

To compare the contribution of mitochondrial BCKA transport in mitochondrial oxidation, we next examined liver-specific *Bckdk*-knockout mice (Figure S6F and S6G). BCKDK deletion activates the BCKD complex by loss of E1a subunit phosphorylation; Yet hepatic BCKDK deletion did not significantly reduce circulating BCKA or BCAA levels under fasted 6 hours conditions (Figure S6H and S6I). These findings demonstrate that mitochondrial transport of BCKA, rather than BCKDH activity alone, exerts rate-limiting control over hepatic BCKA catabolic flux.

We directly demonstrate SLC25A35 mediated BCKA catabolism in cells, we re-expressed human SLC25A35 in the primary adipocytes isolated from SLC25A35-KO mice (SLC25A35-KO^slc25a35,^ Figure S6J and S6K), followed by incubation with 100 µM of uniformly ^13^C labeled BCKA (Figure 4E and S6L). Human SLC25A35 accelerated the time-dependent disappearance of labeled BCKA from culture medium and increased the production of their corresponding labeled BCAA (Figure 4F and 4G). Consistent with a restored BCKA uptake activity, we also observed a significant decrease in unlabeled KIC (M+0), KMV (M+0), and KIV (M+0) levels in the media from SLC25A35 expressing adipocytes compared to the KO group at 24 hours post-incubation (Figure S6M). As expected from increased BCAA catabolic flux from elevated mitochondrial BCKA, the unlabeled Leu (M+0), Ile (M+0), and Val (M+0) were also reduced in the media over time along with unlabeled BCKA (Figure S6N). Together, these rescue experiments support a conserved role for human and mouse SLC25A35 in mitochondrial BCKA transport and metabolism.

To directly assess mitochondrial BCKA uptake, mitochondria were isolated from BAT of wildtype control and SLC25A35-KO mice and then incubated with uniformly ^13^C-labeled KIC, KMV and KIV for 0, 5, 15 and 30 minutes (Figure 4S), control mitochondria displayed a progressive accumulation of labeled KIC and KMV, whereas their accumulation was markedly blunted in SLC25A35-deficient mitochondria (Figure 4T). KIV accumulation was comparable between groups at the earlier time points but was significantly reduced in KO mitochondria after 30 minutes. We next measured the corresponding labeled BCAA generated during the mitochondrial BCKA import assay. Control mitochondria showed time-dependent accumulation of ^13^C-leucine (M+6), ^13^C-isoleucine (M+6), and ^13^C-valine (M+5) (Figure S6O). Their accumulation was generally reduced in SLC25A35-KO mitochondria, particularly for isoleucine and valine. Together, these findings support a direct role for SLC25A35 in mitochondrial BCKA import, with a stronger contribution to the rapid uptake of KIC and KMV and a more limited or slower contribution to KIV transport.

It is possible that SLC25A35 mediated BCKA uptake is indirectly contributed by BCAA uptake as they are coupled in mitochondria. To investigate this question, we studied genetic interaction between SLC25A35 with SLC25A44, a recently^42,43^. We generated SLC25A44 single KO (SLC25A44-KO) and SLC25A35/A44 double-knockout mice (SLC25A35/A44-DKO) (Figure S7A and S7B) and measured the serum metabolites of BCKA through metabolomic analysis under fasting and refeeding conditions (Figure S7D-S7F). The results showed SLC25A44 KO had no significant impact on circulating BCKA levels under either feeding or fasting conditions, in wildtype or SLC25A35 KO background, implicating that SLC35A35 mediated BCKA regulation is independent from SLC25A44 mediated BCAA uptake during nutrient dependent responses.

### Structural determinants of SLC25A35-mediated BCKA transport

Phylogenetic analysis identified SLC25A35 orthologs across diverse vertebrate lineages, including mammals, birds, reptiles, amphibians, and fish (Figure S8A). Multiple-sequence alignment of human, canine, bovine, mouse, rat, and zebrafish SLC25A35 revealed strong conservation, particularly within the membrane-spanning carrier core (Figure S8B), suggesting a conserved transport function. We performed in-silico substrate-binding analysis of human SLC25A35 by screening 303 mitochondrial metabolites for predicted interactions using Boltz-2^44^. KIC, was ranked fourth among the potential metabolites, with a confidence score of 0.621 compared with a median score of 0.34, after oxaloacetate, pyruvate and hydroxypyruvate (Figure 5A). The predicted model positioned KIC within the central cavity formed by the six transmembrane helices, approximately 2.5-4.3 Å from Gln73, Arg175, Met223, and Arg276 (Figures 5B and 5C). Guided by the predicted binding pose, evolutionary conservation and human genetic evidence, we generated variants at predicted KIC-interacting residues (R175A and Q73A/R262A), a conserved arginine (R232A), and Gly262, the position corresponding to the human p.Gly262Asp variant associated with circulating KIC levels (G262A), and expressed these variants in SLC25A35-KO adipocytes (Figure S8C and S8D). Based on BCKA isotope labelling, R175A, R232A, and G262A variants showed different degrees of impairment compared to the wild-type SLC25A35 for clearance of ^13^C-KIC (M+6) and ^13^C-KIV (M+5) from the culture medium. Clearance of ^13^C-KMV (M+6) was impaired by R232A and G262A, but not by R175A, whereas the Q73A/R276A double variant behaved comparably to wild-type SLC25A35 in the cell-based assay (Figures 5E–5G). In isolated mitochondria expressing wild-type or corresponding mutant SLC25A35 incubated with the ^13^C-labeled BCKA for 5 or 30 minutes. At 5 minutes time point, the formation of ^13^C-leucine (M+6), ^13^C-isoleucine (M+6), and ^13^C-valine (M+5) was comparable between wild-type and mutant mitochondria. At 30 minutes time point, however, mitochondria expressing R175A, R232A, or Q73A/R276A were detected with significantly lower amounts of the corresponding labeled BCAAs than those expressing wild-type SLC25A35, whereas G262A showed only modest effects (Figures 5H–5J). Overall, these results provided additional corroborated evidence to support SLC25A35 as a BCKA transporter.

**Figure 5.**
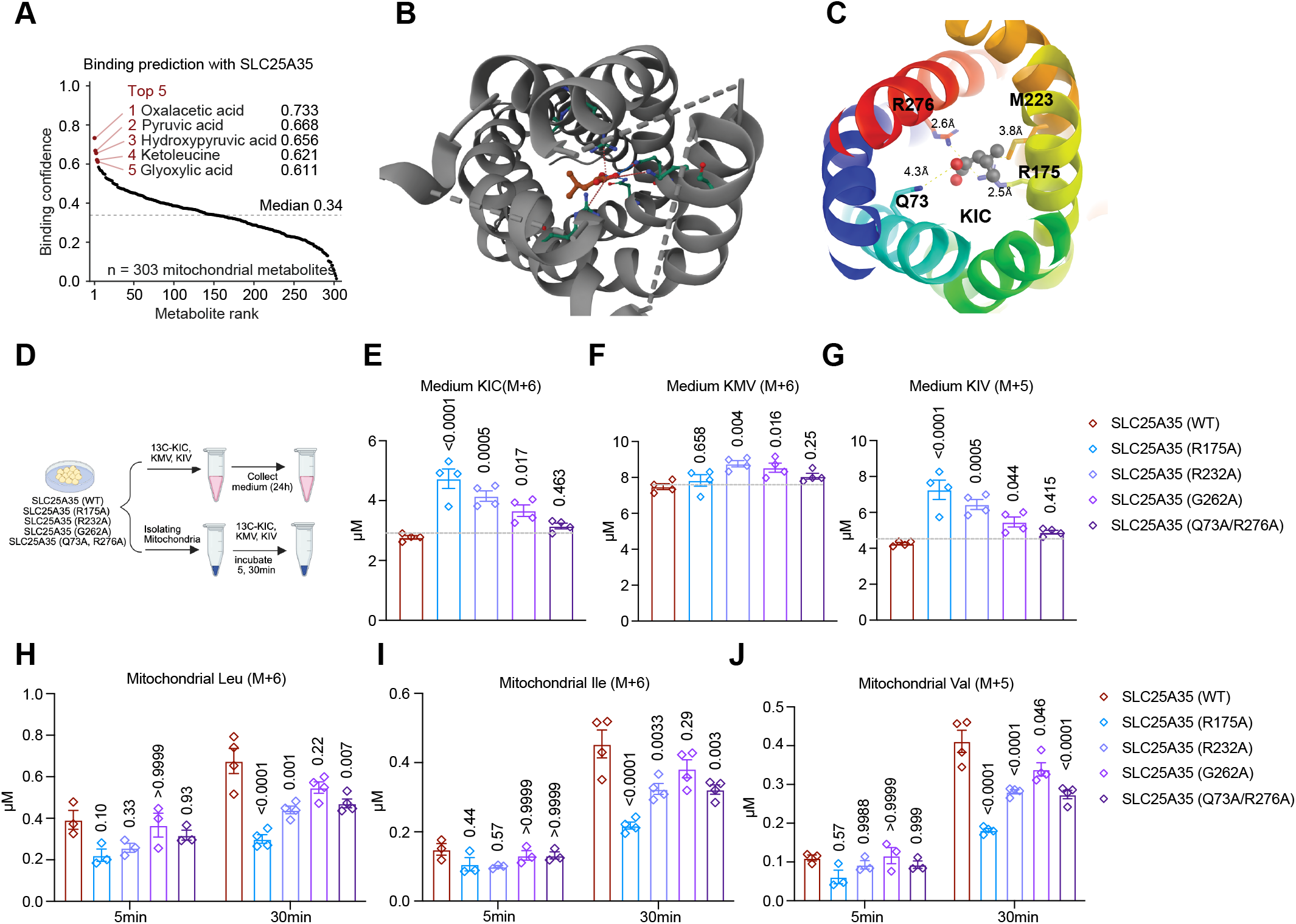
Structural determinants of SLC25A35-dependent BCKA handling. (A) Binding confidence predicted by Boltz-2^44^ for 303 mitochondrial metabolites with human SLC25A35 protein. The five top-ranked metabolites and their binding confidence are indicated. (B) AlphaFold 3-predicted structure^67^ of human SLC25A35 represented in cartoon using PyMOL 3.1. (C) Top view of the predicted KIC-binding conformation. The transmembrane (TM) helices of SLC25A35 are colored using the PyMOL rainbow preset with TM1 in blue to TM6 in red. KIC and the residues within 5 Å are displayed in sticks and labeled, with distances in Å. (D) Experimental schematic. Cells expressing wild-type SLC25A35 (WT) or the indicated variants (R175A, R232A, G262A or Q73A/R276A) were incubated with uniformly ^13^C-labeled KIC, KMV and KIV, and medium was collected after 24 hours (E–G) or mitochondria were isolated and incubated with the tracers for 5 and 30 mins (H–J). (E–G) Medium levels of ^13^C-KIC (M+6) (E), ^13^C-KMV (M+6) (F) and ^13^C-KIV (M+5) (G) after 24 hours incubation with ^13^C-BCKA in cells expressing wild-type SLC25A35 or the indicated variants (n = 4). Dashed lines indicate the mean level in cells expressing wild-type SLC25A35; P values are for the comparison with wild-type SLC25A35. (H–J) Mitochondrial levels of ^13^C-Leu (M+6) (H), ^13^C-Ile (M+6) (I) and ^13^C-Val (M+5) (J) after 5 and 30 mins incubation of isolated mitochondria with ^13^C-BCKA (n =3-4). P values are for the comparison with wild-type SLC25A35. Data are presented as mean ± SEM. P values in panels (E–G) were determined by one-way ANOVA followed by Dunnett’s multiple comparisons test. P values in panels (H–J) were determined by two-way ANOVA followed by Dunnett’s multiple comparisons test.

### SLC25A35 couples BCKA transport to adipocyte lipid storage

Having established that SLC25A35 is enriched in adipose tissue and mediates mitochondrial BCKA uptake, we next investigated whether this transport activity contributes to adipocyte lipid metabolism. SLC25A35 expression was markedly increased during human adipogenesis at both the transcript and protein levels (Figure 6A), paralleling its induction during mouse brown-adipocyte differentiation (Figures S9B and S9C). Across the AL–F–R cycle, *Slc25a35* ranked among the top 20 genes in iWAT according to GEIaS, alongside genes with established roles in lipid metabolism, including *Fasn*, *Scd1*, *Acly*, and *Dgat2* (Figure S9A). Pearson correlation analysis for lipogenesis vs. lipolysis scores showed Slc25a35 expression was positively associated with lipogenesis score but not with the lipolysis score (Figure 6B). Analysing a public single-cell gene expression dataset from fat tissue, SLC25A35 was identified to be enriched in brown and white adipocytes, parallel with similar enrichment profiles as Pck1 and BCAA metabolic score (Figure 6C). Consistent with this single-cell analysis, fractionation of adipose tissue into mature adipocytes and the stromal vascular fraction (SVF) showed that *Slc25a35* and *Pck1* were enriched in mature adipocytes, whereas *Pck2* was expressed predominantly in the SVF (Figure S9D-S9G). Overexpression of SLC25A35 in wildtype preadipocytes led to increased lipid accumulation (Figure 6D), accompanied by the induction of genes involved in adipogenesis, lipogenesis, and lipid handling, including *Pparg*, *Fasn*, *Dgat1*, *Dgat2*, *Cd36*, *Lpl*, and *Atgl*. SLC25A35 overexpression also significantly increased cytosolic PEP-generating enzyme PCK1, as well as *Bckdk* and *Bckdha*, which participate in the regulation and oxidation of BCKAs, respectively (Figure 6E). Consistently, re-expression of SLC25A35 in SLC25A35-KO preadipocytes restored lipid accumulation (Figure 6F) and increased the abundance of fatty acid species, with the exception of several very-long-chain and polyunsaturated fatty acids (Figure 6G). Results from BCKA tracing showedSLC25A35 expression also increased the incorporation of BCKA-derived carbon into the monomethyl branched-chain fatty acids FA 15:0 (M+3), FA 17:0 (M+3), and iso/anteiso FA 17:0 (M+5), with a similar trend observed for iso/anteiso FA 16:0 (M+4) (Figure 6H). These findings connect SLC25A35-mediated BCKA transport with the utilization of BCKA-derived carbon for adipocyte fatty acid synthesis. These findings connect SLC25A35-mediated BCKA transport with the utilization of BCKA-derived carbon for adipocyte fatty acid synthesis. Consistent with impaired BCKA transport, the SLC25A35 mutants (R175A and R232A) were substantially less effective to restore lipid accumulation in SLC25A35 KO adipocytes (Figure 6I and 6J), These results directly link the BCKA-transport function of SLC25A35 to its effects on adipocyte lipid accumulation and fatty acid composition.

**Figure 6.**
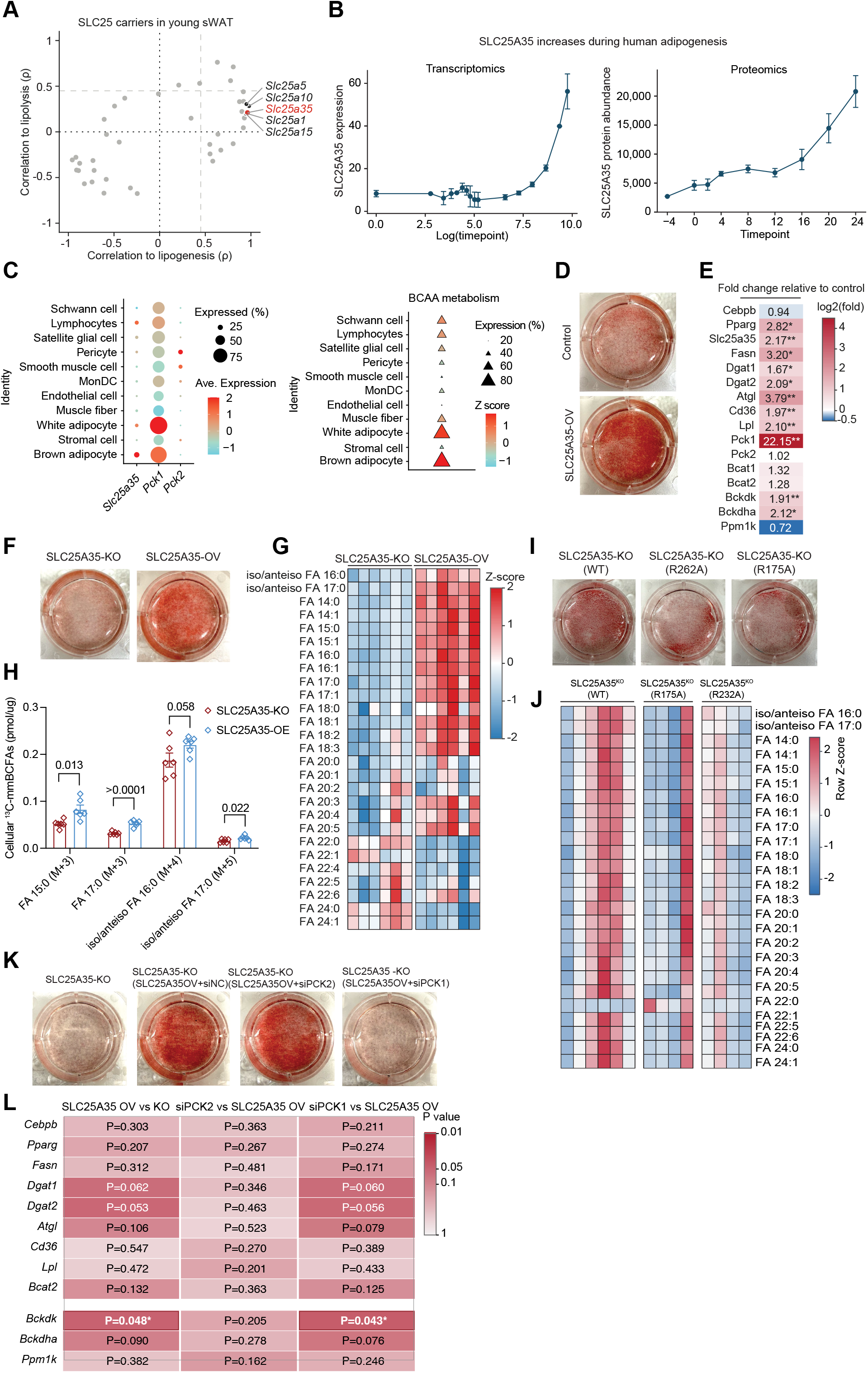
SLC25A35 couples mitochondrial BCKA handling to adipocyte lipid storage. (A) *SLC25A35* mRNA (left) and SLC25A35 protein (right) abundance during human adipogenesis (https://adiposetissue.org/). (B) Pearson correlation of each SLC25 family gene with a lipogenesis score (x axis) and a lipolysis score (y axis) across young subcutaneous white adipose tissue (sWAT) of mice in the ad libitum, fasted and refed states from published bulk RNA-seq data. Lipogenesis score: *Acly*, *Acaca*, *Fasn* and *Scd1*^68^; lipolysis score: *Pnpla2*, *Lipe*, *Mgll* and *Abhd5*^69^. (C) *Slc25a35*, *Pck1* and *Pck2* mRNA level (left panel) and the BCAA metabolism pathway activity (right panel) across the cell types of mouse brown adipose tissue. (D) Oil Red O staining of control and SLC25A35-overexpressing (SLC25A35-OV) adipocytes after differentiation. (E) Heatmap of the fold change in the expression of the indicated genes in SLC25A35-OV adipocytes relative to control adipocytes, measured by RT-qPCR. Numbers show the fold change; *P < 0.05, **P < 0.01. (F) Oil Red O staining of SLC25A35-KO and SLC25A35-overexpressing (SLC25A35-OV) adipocytes. (G) Heatmap of the fatty acid species in total lipid extracts of SLC25A35-KO and SLC25A35-OV adipocytes (n = 6). (H) Incorporation of ^13^C-labeled BCKA into monomethyl branched-chain fatty acids (mmBCFAs): FA 15:0 (M+3), FA 17:0 (M+3), iso/anteiso FA 16:0 (M+4) and iso/anteiso FA 17:0 (M+5) in SLC25A35-KO and SLC25A35-overexpressing (SLC25A35-OV) adipocytes (n = 6). (I) Oil Red O staining of SLC25A35-KO adipocytes re-expressing wild-type SLC25A35 (WT) or the R175A or G262A variant. (J) Heatmap of the fatty acid species in total lipid extracts of SLC25A35-KO adipocytes re-expressing wild-type SLC25A35 (WT) or the R175A or R232A variant (n = 4-6). (K) Oil Red O staining of SLC25A35-KO adipocytes and of SLC25A35-KO adipocytes overexpressing SLC25A35 and transfected with control siRNA (siNC), *Pck2* siRNA (siPCK2) or *Pck1* siRNA (siPCK1). (L) Heatmap of the P values for the comparison of gene expression between SLC25A35-overexpressing and SLC25A35-KO cells (SLC25A35-OV vs KO) and between siPCK2- or siPCK1-transfected and untransfected SLC25A35-overexpressing cells; *P < 0.05. Data are presented as mean ± SEM. P values in panel (H) were determined by unpaired two-tailed Student’s t tests. P values in panels (E and L) were determined by unpaired two-tailed Student’s t tests.

Among the metabolic genes examined, *Pck1* showed the strongest induction following SLC25A35 overexpression in adipocytes suggesting a functional connection between SLC25A35-mediated BCKA metabolism and PCK1-dependent gluconeogenesis. Consistent with this hypothesis, knockdown of *Pck1* significantly reduced the lipid accumulation induced by SLC25A35 overexpression, whereas *Pck2* knockdown had no detectable effect (Figure 6K). *Pck1* knockdown also blunted gene expression changes induced by SLC25A35 expression (Figure 6L). *Pck1* expression increased during adipocyte differentiation and was enriched in mature adipocytes, whereas *Pck2* expression decreased during differentiation and was detected predominantly in the SVF (Figure S9I). Moreover, *Pck2* knockdown did not alter either lipid accumulation or the transcriptional response to SLC25A35 overexpression (Figures S9D–S9J).

Together, these findings support a model in which SLC25A35 promotes adipocyte lipid storage through two complementary mechanisms: incorporation of BCKA-derived carbon into fatty-acid acyl chains and activation of a PCK1-dependent glyceroneogenic pathway that supports glycerol-backbone production and fatty acid esterification. Thus, PCK1, but not PCK2, appears to be the principal metabolic effector associated with SLC25A35-mediated BCKA transport and triglyceride accumulation.

### SLC25A35 deficiency attenuates cold-responsive metabolic signalling in BAT

Given that SLC25A35 is a nutrient-responsive gene, we next examined whether its loss affected whole-body adaptation to feeding–fasting transitions. Locomotor activity, food intake, and heat production were comparable between SLC25A35-KO and control mice (Figure 7A). However, KO mice showed a transiently higher RER during early fasting, suggesting a delayed shift from carbohydrate to fatty acid utilization. This altered metabolic flexibility prompted us to examine their response to cold-induced thermogenic stress.

**Figure 7.**
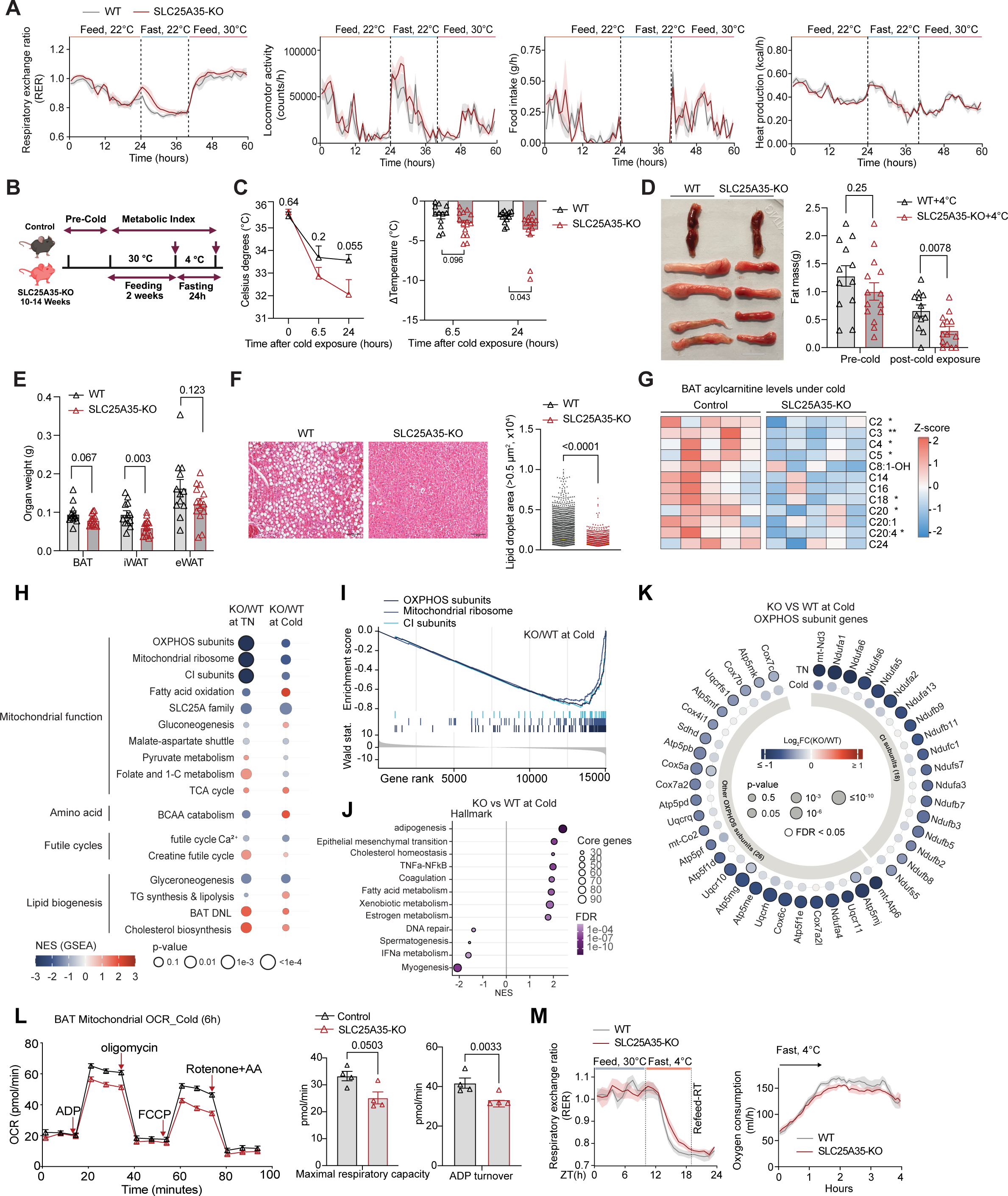
SLC25A35 deficiency impairs mitochondrial and thermogenic adaptation in BAT. (A) Respiratory exchange ratio (RER), locomotor activity, food intake and heat production of control and SLC25A35-KO mice measured by indirect calorimetry. Mice were fed at 22 °C, fasted at 22 °C and refed at 30 °C, as indicated (n = 4-5). Lines show the mean and shading the SEM. (B) Experimental schematic illustrating the cold-challenge protocol, in which control and SLC25A35-KO mice (10–14 weeks old) were housed at thermoneutrality (30 °C) for 2 weeks after a defined pre-cold period and then fasted and exposed to an acute 24 h cold challenge (4 °C). (C) Core body temperature of control and SLC25A35-KO mice at 0, 6.5 and 24 hours of cold exposure (4 °C) (n = 12-14). Right panel showed the body temperature changes to baseline at 6.5 and 24 hours of cold exposure (n = 12-14) (D) Representative images of BAT, iWAT, and eWAT from control and SLC25A35-KO mice after cold exposure (4 °C). Fat mass composition before and after cold exposure (4 °C) in control and SLC25A35- KO mice (right panel, n= 12-14) (E) Weights of brown adipose tissue (BAT), inguinal white adipose tissue (iWAT), and epididymal white adipose tissue (eWAT) from control and SLC25A35-KO mice following cold exposure (n = 12-14). (F) Representative hematoxylin and eosin (H&E)–stained sections of BAT from control and SLC25A35- KO mice following cold exposure. Scale bars, 50 µm. Quantification of adipocyte lipid droplet area from BAT. Analysis was performed using three histological sections per mouse, with entire fields quantified per section (right panel). (G) Heatmap of acylcarnitine abundance in BAT from control and SLC25A35-KO mice after cold exposure, highlighting broad reductions in short- and long-chain acylcarnitines in SLC25A35-deficient BAT (n = 5). *P < 0.05, **P < 0.01. (H) Metabolic pathway enrichment analysis in BAT of SLC25A35-KO relative to wild-type mice at thermoneutrality (TN) and in the cold (n = 6). (I) Running enrichment scores (top) and ranked Wald statistic (bottom) of the MitoCarta3.0 OXPHOS subunit, mitochondrial ribosome and complex I subunit gene sets for KO versus wild-type BAT in the cold. (J) Hallmark gene sets enriched in SLC25A35-KO versus wild-type BAT in the cold. (K) OXPHOS subunit genes (MitoCarta3.0)^70^ with FDR < 0.05 for KO versus wild type at TN or in the cold, grouped as complex I subunits (18 genes) and other OXPHOS subunits (26 genes). (L) Seahorse analysis of mitochondrial oxygen consumption rates (OCR) in mitochondria isolated from BAT of control and SLC25A35-KO mice following 6 hours of cold exposure (left panel). Quantification of maximal respiratory capacity and ADP turnover derived from the Seahorse assay (right panel, n = 4). (M) RER and oxygen consumption of control and SLC25A35-KO mice during feeding at 30 °C, fasting at 4 °C and refeeding at room temperature (RT) (n = 4-5). Data are presented as mean ± SEM. P values in panel (C, left) were determined by two-way ANOVA followed by Tukey’s multiple comparisons test. P values in panels (C, right and D) were determined by one-way ANOVA followed by Dunnett’s multiple comparisons test. P values in panels (F and S–T) were determined by unpaired two-tailed Student’s t tests. P values in panel (E) were determined by multiple unpaired t tests with false discovery rate (FDR) correction. P values in panel (G) were determined by unpaired two-tailed Student’s t tests. P values in panels (H-J) are described in the STAR Methods.

SLC25A35 exhibits the highest expression in BAT relative to other organs, a pattern closely mirroring the core genes involved in BCAA catabolism (Figure 1F and 6C). BAT is a central organ for adaptive thermogenesis, where mitochondrial metabolic flexibility is essential for efficient heat production in response to cold exposure^47,48^. Previous studies have demonstrated that disruption of BCAA catabolism in BAT profoundly impair cold-induced thermogenesis^49,50^, despite the relatively modest contribution of BCAA oxidation to total energy expenditure. These observations support the idea that BCAA-derived metabolites play a permissive role in supporting thermogenic function^51^. So, we subjected the SLC25A35-KO and control mice to an acute cold challenge (Figure 7B). Following two weeks of thermoneutral housing (Figure S10A), no differences were detected between control and SLC25A35- KO mice in body weight, body composition, fat mass percentage, organ weights, or BAT adipocyte size, indicating comparable basal metabolic and adipose tissue status (Figure S10B-S10E). Cold exposure regulated SLC25A35 expression in a tissue and nutritional-state-dependent manner, reducing its mRNA in skeletal muscle under fed conditions and in iWAT during fasting. In BAT, SLC25A35 mRNA remained unchanged, whereas SLC25A35 protein abundance appeared lower during fasting (Figure S10F-S10H). However, during cold exposure, SLC25A35-KO mice exhibited a greater decline in core body temperature compared with controls, indicating impaired cold-induced thermogenesis (Figure 7C). This defect was accompanied by a significant reduction in fat content following cold challenge in SLC25A35-KO mice measured by MRI (Figure 7D). In contrast, total body weight and lean mass were not significantly different between genotypes (Figure S10I and S10J). Consistent with the preferential involvement of adipose tissue in cold-induced thermogenesis, BAT and iWAT weights were significantly reduced in SLC25A35-KO mice following cold exposure compared to the controls, whereas eWAT exhibited a trend of decrease (Figure 7D and 7E). In contrast, weights of non-adipose organs, including liver, heart, and skeletal muscle, were not changed between SLC25A35-KO mice and controls, indicating that SLC25A35 deficiency selectively affects adipose tissue responses to cold (Figure S10K). Histological examination further revealed profound alterations in lipid content in the brown adipose tissue of SLC25A35-KO mice after cold exposure. Hematoxylin and eosin (H&E) staining showed a marked depletion of intracellular lipid droplets in BAT from SLC25A35-KO mice compared with controls (Figure 7F). Quantitative analysis confirmed a significant reduction in lipid droplet size and total lipid droplet area in the SLC25A35-KO BAT tissue (Figure 7F). Concordantly, circulating levels of non-esterified fatty acids (NEFAs) were significantly reduced in SLC25A35-KO mice following cold challenge (Figure S10L), suggesting diminished availability of fatty acids to support thermogenesis. Because sustained cold-induced thermogenesis requires the coordinated delivery and oxidation of fatty acid- and amino acid-derived substrates, we next asked whether mitochondrial carbon handling was altered in SLC25A35-KO BAT. We therefore assessed mitochondrial acyl-group buffering and fatty acid oxidative metabolism by profiling the acylcarnitine pool. Cold-exposed SLC25A35-KO BAT exhibited a broad reduction in acylcarnitine species, including a significant decrease in acetylcarnitine (C2), which reflects mitochondrial acetyl-group availability and buffering (Figure 7G). Short-chain acylcarnitines, including C3, C4, and C5 species that can arise from BCAA/BCKA catabolism, were also decreased, consistent with reduced utilization of BCKA-derived carbon. Long-chain acylcarnitines were similarly reduced, suggesting diminished delivery or oxidation of fatty acid-derived acyl groups during thermogenic activation.

To determine whether these metabolic alterations were accompanied by transcriptional remodelling, we performed bulk RNA-sequencing of BAT from wild-type and SLC25A35-KO mice housed at thermoneutrality (TN) or exposed to cold. Principal component analysis separated samples primarily by temperature along PC1, with an additional genotype-dependent separation between wild-type and SLC25A35-KO BAT at TN (Figure S11A). Relative to wild-type controls, SLC25A35-KO BAT exhibited 882 differentially expressed genes at TN (482 upregulated and 400 downregulated) and 407 during cold exposure (202 upregulated and 205 downregulated) (Figure S11B and S11C). Apart from *Slc25a35*, most members of the *Slc25* family remained unchanged, although its closest paralogue, *Slc25a34*, was markedly reduced in KO BAT at TN (Figure S11D). At TN, GSEA revealed coordinated suppression of oxidative phosphorylation and enrichment of cell-cycle, inflammatory-response, and epithelial– mesenchymal transition programs in SLC25A35-KO BAT (Figure S11E and S11F). Consistently, analysis of MitoCarta3.0 pathways identified OXPHOS subunits and complex I subunits as the most strongly depleted gene sets, with reduced expression broadly distributed across OXPHOS subunits (Figure 7H and 7I). During cold exposure, these mitochondrial gene sets retained negative enrichment scores but were no longer significant, suggesting partial attenuation of the transcriptional defect (Figures 7H and 7K). Conversely, fatty acid oxidation and BCAA catabolism showed directionally positive enrichment in KO BAT. Adipogenesis was the most positively enriched Hallmark pathway, accompanied by enrichment of cholesterol homeostasis and fatty acid metabolism (Figure 7H and 7J). These changes may represent a compensatory transcriptional response to impaired mitochondrial substrate utilization during cold stress.

We next assessed whether this transcriptional signature was accompanied by functional impairment of mitochondrial respiration. Following 6 hours of cold exposure, mitochondria isolated from SLC25A35-KO BAT exhibited reduced ADP turnover and a trend toward lower maximal respiratory capacity (Figure 7L). Maximal respiration was also reduced in SLC25A35-KO brown adipocytes following norepinephrine stimulation (Figure S10M). At the whole-body level, fasted KO mice exposed to 4 °C tended to maintain a higher RER and exhibited lower heat production than controls (Figure 7M). Together, these transcriptomic and functional findings indicate that SLC25A35 deficiency compromises mitochondrial oxidative capacity in BAT, while cold-induced activation of lipid- and BCAA-metabolic programs provides only partial compensation.

To determine whether the metabolic phenotype was accompanied by coordinated remodeling of canonical fuel-utilization pathways, we examined transcriptional and protein-level regulators of lipid and glucose metabolism in BAT. Consistent with the RNA-seq results, expression of the mitochondrial respiratory-chain subunit genes Cox8b and Cox7a1 was reduced in SLC25A35-KO BAT at thermoneutrality, together with a significant reduction in the BCAA-catabolic gene Bcat2. During cold exposure, Ucp1 and pparr expression was significantly reduced in KO BAT, whereas most other genes involved in fatty acid synthesis, lipid handling, fatty acid oxidation, thermogenesis, glucose oxidation, and BCAA catabolism remained unchanged (Figure S11G). Across these pathways, most transcripts remained unchanged or showed only modest trends, indicating a lack of coordinated transcriptional remodeling among the targeted metabolic genes during cold exposure. Given these limited transcriptional changes, we next asked whether SLC25A35 deficiency impaired thermogenic activation at the level of protein signaling. Phosphorylation of acetyl-CoA carboxylase (ACC) at Ser79, a canonical downstream readout of AMPK activation, was reduced in SLC25A35-KO BAT, consistent with attenuated AMPK-mediated energy-stress signaling^52,53^ (Figure S11H). Phosphorylation of ATP-citrate lyase at Ser455, an AKT-associated activating site, was also reduced, suggesting diminished nutrient-responsive metabolic signaling^54^ (Figure S11I). Immunoblot analysis further revealed attenuated β-adrenergic–PKA pathway activation in KO BAT, accompanied by reduced protein abundance of the lipolytic enzymes ATGL and HSL ^55,56^(Figure S11H). Together, these findings suggest that SLC25A35 deficiency primarily disrupts the signaling and protein-level mechanisms coordinating lipid mobilization, metabolic remodeling, and mitochondrial fuel supply during cold-induced thermogenesis, rather than inducing a broad transcriptional adaptation of canonical metabolic genes. To determine whether these alterations were accompanied by changes in BCKA metabolism, we next profiled BCKAs and their downstream metabolites in serum and BAT (Figure S12A). During cold exposure, SLC25A35-KO mice exhibited elevated circulating KIC and reduced serum concentrations of the ketone bodies acetoacetate and β-hydroxybutyrate, together with HIC accumulation in BAT (Figures S12B and S12C). The concurrent accumulation of KIC and its reduction product HIC, combined with decreased ketone-body and acylcarnitine pools, is consistent with impaired mitochondrial disposal of BCKA-derived carbon and its increased diversion toward cytosolic reduction. Collectively, these findings link defective SLC25A35-mediated BCKA transport to altered BCKA partitioning, mitochondrial dysfunction, and broader disruption of fatty acid and ketone metabolism during cold-induced thermogenesis.

## DISCUSSION

Here, we identify SLC25A35 as a nutrient-responsive mitochondrial carrier that regulates BCKA compartmentation and catabolism. This conclusion is supported by convergent evidence from human genetic associations, nutrient-responsive transcriptomics, mouse genetics, isotope tracing, gain- and loss-of-function experiments, cultured adipocytes, and isolated-mitochondrial uptake assays. Although the current paradigm for BCAA/BCKA homeostasis mainly focuses on their catabolic activities via intra- mitochondrial enzyme cascades^17–19,57^. our study reveals for the first time an additional regulatory component, involving BCKA uptake and compartmentation across the mitochondrial membrane via a highly specific mitochondrial transporter.

In our study, we find systemic levels of BCAA remain largely unchanged under starvation conditions; however, circulating BCKA levels are significantly elevated. BCKA are the obligatory catabolic intermediates of BCAA, the observed uncoupling between BCAA and BCKA circulating levels during fasting-refeeding cycle is very much unexpected. However, this phenomenon can be explained if BCAA and BCKA are compartmented each through their dedicated transporters. We find SLC25A35 expression is markedly suppressed during starvation across all the tissues analyzed, and SLC25A35 deficiency significantly increased circulating BCKA levels at basal feeding state. Therefore, it is more plausible that BCKA catabolism is reduced under fasting, contributed mainly by suppressed SLC25A35 expression and mitochondrial uptake. In our study, we have observed the impact of SLC25A35 expression on systemic BCKA clearance via global gene inactivation and multi-tissue re-expression approach. However, Hepatic SLC25A35 re-expression partially reduced circulating KIC and KMV but did not fully normalize their concentrations. In comparison, liver-specific *Bckdk* deletion, which relieves inhibitory phosphorylation of BCKDH, did not substantially alter circulating BCKAs or BCAAs following fasting. These observations further support the notion that BCAA and BCKA regulation is not just contributed by mitochondrial catabolic capacity (such as BCKDH) but also BCAA/BCKA mitochondrial flux through dedicated transporters.

Because BCKA are the obligate catabolic intermediates of BCAA, the observed uncoupling between BCAA and BCKA circulating levels during fasting-refeeding cycle is very much unexpected. However, this phenomenon can be explained if BCAA and BCKA are compartmented each through their dedicated transporters. Indeed, loss of SLC25A35 leads to pronounced accumulation of BCKA in the cytosol and extracellular space without significant changes in BCAA levels. In contrast, inactivation the known BCAA transporter, SLC25A44^42,43^, results in BCAA accumulation without significant impact on BCKA. It is interesting to note that SLC25A35 demonstrates different substrate activity towards different BCKA species, with KIC and KMV preferred over KIV. The molecular basis and the functional relevance of this biased substrate specificity remain to be explored. However, this hierarchy mirrors the known substrate preference based on biochemical assay and molecular docking analysis for BCKDH complex^16^. suggesting that SLC25A35 may be tuned to align mitochondrial transport capacity with downstream enzyme activities in terms of substrate specificity.

Yamamuro *et al* has recently reported that SLC25A35 is an exporter for phosphoenolpyruvate, a high energy metabolite for lipogenesis and gluconeogenesis^46^. In their study, lower mitochondrial levels of KIC and KIV were also observed following SLC25A35 inactivation, consistent with its role as a BCKA mitochondria importer. Although direct measurements using a cell-free SLC25A35 loaded liposome did not detect transporter activity for BCKA substrates in their study. In the metabolomic analysis from BAT, we did not observe a significant change in PEP levels in the SLC25A35 mice. Importantly, we showed here the cytosolic derived PEP by pck1 has a significant role in lipogenesis rather than the mitochondrial derived PEP by pck2. In addition, AI-driven structural binding predictions revealed that SLC25A35 shows a much higher binding confidence for KIC^44^. Since each member of the SLC25A family genes often possesses transporter activities for multiple substrates, it is an intriguing possibility that SLC25A35 is an anti-transporter for both PED efflux and BCKA influx across mitochondrial membrane.

Previous studies have demonstrated that BCAA metabolic flux differs markedly among different tissues, despite relatively constant BCAA levels in circulation^29^. SLC25A35 is highly enriched in BAT, and its expression increases with adipocyte differentiation and BCAA catabolic acitivity^47,48^. Although Loss of SLC25A35 has minimal impact on basal lipid accumulation under thermoneutral conditions, gain of function experiments revealed that SLC25A35 promotes adipocyte lipid accumulation and fatty acid remodeling. This is consistent with previous evidence that BCAA catabolism supplies lipogenic acetyl- CoA and branched acyl-CoAs for fatty-acid synthesis^58,59^. Consistently, SLC25A35 expression increased lipid accumulation and promoted incorporation of BCKA-derived carbon into odd-chain and monomethyl branched-chain fatty acids. Transport-impaired variants were less effective in restoring these lipid phenotypes, linking BCKA handling to fatty-acid remodeling. The requirement for PCK1, but not PCK2, further suggests that SLC25A35 supports lipid storage through both BCKA-derived acyl-chain synthesis and PCK1-dependent glyceroneogenesis. The relative contributions of these pathways are likely to depend on adipocyte type and nutrient state. However, during cold exposure, when thermogenic demand is high, SLC25A35 deficiency results in a profound impairment of heat production. Notably, this thermogenic defect does not arise from limited fuel availability. Instead, lipid mobilization is enhanced as evidenced by accelerated lipid droplet depletion, reduced circulating non-esterified fatty acids, and pronounced loss of fat mass. In addition, mitochondrial respiration is markedly reduced in adipocytes lacking SLC25A35, indicating that BCKA catabolic activity has a potential impact on mitochondrial oxidation. At the metabolic level, loss of SLC25A35 is associated with depletion of short- and medium-chain acyl-carnitines, including branched-chain acyl-carnitine species. Together, these findings support a model in which SLC25A35 provides permissive metabolic inputs required to sustain fatty acid–driven thermogenesis^60,61^. In the absence of SLC25A35, lipid mobilization proceeds, but mitochondrial oxidative competence becomes constrained, leading to uncoupled heat production despite of adequate substrates. While SLC25A35 mediated PEP export and lipogenesis may also contribute to this outcome, other mechanisms, including BCAA/BCKA mediated signaling and their direct contribution to the futile cycle of downstream products, such as branched fatty acids may also be important in energy homeostasis and thermogenesis^60^. The exact mechanisms involved in BCKA and SLC25A35 mediated regulation of thermogenic activity in BAT should be further dissected in future studies.

Collectively, our findings identify SLC25A35 as a selective, nutrient-regulated mitochondrial BCKA transporter that operates independently of mitochondrial BCAA import and is required for adaptive thermogenesis. By uncovering a previously unrecognized mitochondrial transport step, this work highlights metabolite transport as an active regulatory checkpoint and positions mitochondrial BCKA entry as a flux-limiting determinant that supports tissue-dependent metabolism of BCKA/BCAA.

### Limitations of the study

Although genetic, metabolomic, isotope-tracing, and isolated-mitochondrial experiments consistently support a requirement for SLC25A35 in mitochondrial BCKA handling, direct transport of purified BCKAs by reconstituted SLC25A35 was not demonstrated. This is particularly important given the reported transport of PEP-but not KIC or KIV-in a purified proteoliposome system; Consequently, whether SLC25A35 transports BCKAs directly, requires additional cofactors or exchange substrates, or regulates their entry indirectly remains unresolved. In addition, the principal physiological studies used global SLC25A35-KO mice, making it difficult to distinguish adipocyte-autonomous effects from contributions of other tissues; adipocyte specific and inducible models, together with studies under chronic metabolic challenges, will be required. Finally, the mechanisms linking altered BCKA compartmentation to mitochondrial dysfunction and thermogenic signaling remain incompletely defined, and the suggestive human genetic association requires independent replication and direct functional evaluation of the naturally occurring variant.

## ACKNOWLEDGMENTS

This work was supported by Singapore National Medical Research Council grants OF-LCG24may-0020 and OFLCG22may-0010 (Y.W.) and PR191670 (C.G., Y.W. and Z.L.). This work was also supported by National Medical Research Council Singapore of OFYIRG24jan-0037 (Y.L.), National Key Research and Development Program of China (2025YFE0126700) (H.S.), Ministry of Education MOET2EP30123 and MOET2EP30224 (D.L.S.), A*STAR Industry Alignment Fund–Industry Collaboration Projects (IAF-ICP) Grant I2501E0055 (F.T.).

## AUTHOR CONTRIBUTIONS

Y.W., Y.L., and C.G. conceived, designed, and supervised the study. D.L.S. and H.S. contributed to data discussion and conceptual development. Y.L., Z.L., G.L.C., L.S., S.R., D.D., and C.M.W. performed the experiments and collected the data. M.L., J.L., Q.Z., D.L.S., F.T., S.L., W.W., and F.H. conducted the data analysis. M.L., J.L., Y.K.W., G.L., and M.W., contributed to the experiments and manuscript editing. All authors reviewed, edited, and approved the final version of the manuscript.

## EXPERIMENTAL MODEL AND STUDY PARTICIPANT DETIALS

### Animals

All animal experiments were performed in accordance with protocols approved by the SingHealth Institutional Animal Care and Use Committee (IACUC) at Duke–NUS Medical School, Singapore. Mice were housed in a specific pathogen–free animal facility under a 12 h light–dark cycle at 22 °C and 50% relative humidity, with ad libitum access to water and a standard irradiated rodent chow diet. For thermoneutral conditions, mice were housed in temperature-controlled incubators set to 30 °C. For cold exposure experiments, mice were transferred to incubators maintained at 4 °C. Cold exposure was initiated when mice were 10–14 weeks of age and was conducted for 24 h, during which food was withdrawn unless otherwise specified. Wild-type mice (strain #000664) and FVB/N-Tg (Zp3-cre) 3Mrt/J mice (strain #003377) were obtained from the Jackson Laboratory. *Slc25a35*^flox/flox^ and *Slc25a44*^flox/flox^ mice were generated at the Transgenic and Reproductive Technology Centre at the university. Global *Slc25a35* and *Slc25a44* knockout mice were generated by crossing the respective floxed mice with Zp3-cre transgenic mice. *Slc25a35*/a44 double-knockout (DKO) mice were generated by crossing *Slc25a35* knockout mice with *Slc25a44* knockout mice. Genotypes were confirmed by PCR analysis of genomic DNA. Core body temperature was measured using an animal electronic thermometer (Alcott Biotech, Shanghai, China). Body weight and food intake were monitored throughout the experiments, and body composition was assessed by magnetic resonance imaging (EchoMRI). Tissues including inguinal white adipose tissue (iWAT), epididymal white adipose tissue (eWAT), brown adipose tissue (BAT), liver, and other organs were excised, weighed, rapidly frozen in liquid nitrogen, and stored at –80 °C until further processing. For metabolite administration experiments, mice were randomly assigned to treatment groups. Experiments included wild-type, *Slc25a35*^flox/flox^, *Slc25a44*^flox/flox^, global *Slc25a35* knockout, global *Slc25a44* knockout, and *Slc25a35/a44* double-knockout mice. All experiments were performed using male littermate mice that were randomly assigned to experimental groups.

### Cold Tolerance Experiment

Mice were pre-exposed to cold in a temperature-controlled incubator at 4 °C for 20 h, followed by transfer to thermoneutral conditions (30 °C) for 14 days to induce thermoneutral adaptation. Mice were then returned to a 4 °C incubator for an additional 24 h. At the end of the experiment, mice were euthanized, and blood, brown adipose tissue (BAT), and other organs were collected for downstream analyses. Core body temperature was recorded every 2 h using an animal electronic thermometer (Alcott Biotech, Shanghai, China).

### Cell Lines

HEK293T, AML12, and HUH7 cells were obtained from the American Type Culture Collection (ATCC). All cell lines were maintained in Dulbecco’s Modified Eagle’s Medium (DMEM) supplemented with 10% fetal bovine serum (FBS) and 1% penicillin–streptomycin (P/S) and cultured in a humidified incubator at 37 °C with 5% CO,.

### Isolation and differentiation of adipose-derived stromal vascular fraction (SVF) cells

BAT and iWAT were collected from 3–4-week-old mice, manually cleared of visible connective tissue, and finely chopped. Tissue fragments were subjected to enzymatic dissociation in phosphate-buffered saline (PBS, pH 7.4) containing 1% bovine serum albumin (BSA) and collagenase type IV (10 mg/mL; Gibco, 17104) at 37 °C for 25–30 min with gentle shaking (500 rpm) in a thermomixer. Enzymatic digestion was terminated by adding two volumes of complete growth medium (DMEM supplemented with 10% fetal bovine serum and 1% penicillin–streptomycin). The resulting cell suspension was filtered through 70- or 100-µm cell strainers to eliminate undigested tissue remnants. Stromal vascular fraction (SVF) cells were collected by centrifugation at 2,000 rpm for 10 min, resuspended in fresh growth medium, and seeded for culture. Cells were maintained at 37 °C in a humidified incubator with 5% CO,.

For adipocytes differentiation, SVF cells were plated in 6-well plates and expanded to full confluence in DMEM containing 10% FBS. Two days after confluence (defined as day 0), differentiation was initiated by switching to induction medium composed of DMEM supplemented with 0.5 mM 3-isobutyl-1- methylxanthine (IBMX; Sigma, I7018), 1 µM dexamethasone (Sigma, D4902), 10 µg/mL insulin (Sigma, I5500), and 10 µM rosiglitazone (Cayman Chemical, 71740). After 48 h, cells were transitioned to maintenance medium (DMEM containing 10% FBS and 10 µg/mL insulin), which was subsequently replaced at 2-day intervals. Mature brown adipocytes (day 6–7) were washed with PBS and fixed in 4% paraformaldehyde for 10 min at room temperature. Lipid accumulation was visualized by staining fixed cells with Oil Red O solution (Sigma) prepared in isopropanol for 1 h. Images were captured using an Olympus microscope equipped with a digital imaging system.

### Lentiviral production and transduction

Lentiviral constructs encoding a scrambled control and wild-type human *SLC25A35* were obtained from VectorBuilder. For lentivirus production, HEK293FT cells at logarithmic growth phase were co-transfected with the expression plasmids together with the packaging vectors pMDLg/pRRE, pRSV-Rev, and pCMV-VSV-G (Addgene). Transfections were performed using X-tremeGENE 9 DNA transfection reagent (Roche) according to the manufacturer’s recommendations. Culture supernatants containing lentiviral particles were harvested 48 h post-transfection, clarified by centrifugation if necessary, and passed through 0.45-µm cellulose acetate filters to remove cellular debris. Primary adipocyte precursors were seeded in 10-cm culture dishes and infected at approximately 30–40% confluence by incubation with lentiviral supernatants supplemented with 8 µg/mL polybrene for 24 h. Following infection, cells were washed and replated in fresh growth medium prior to initiation of adipocytes differentiation. Expression of wild-type human SLC25A35 was verified by quantitative PCR and/or immunoblot analysis.

### Glucose Tolerance Test

Glucose tolerance tests were performed following established procedures with minor modifications. Briefly, SLC25A35-KO and control mice were fasted for 6 h (09:00–15:00) with free access to water. Mice were then administered an intraperitoneal injection of D-glucose (1.5 g/kg body weight; Sigma). Blood glucose levels were measured from tail vein blood at baseline (0 min) and at 15, 30, 60, and 120 min after glucose administration using a handheld glucometer (ACCU-CHEK, Singapore).

### Isolation of mitochondria from mouse tissues

For mitochondrial isolation from mouse tissues^62^, mice were fasted overnight (19:00–09:00) prior to tissue collection. Mice were euthanized by cervical dislocation, and mouse tissues were rapidly excised and processed immediately for mitochondrial isolation. For liver mitochondria isolation, tissues were homogenized in IBc buffer containing 10 mM Tris–MOPS (pH 7.4), 5 mM EGTA/Tris, and 200 mM sucrose. For BAT and other organs mitochondria isolation, tissues were homogenized in IBm1 buffer containing 50 mM Tris–HCl (pH 7.4), 10 mM EDTA, 67 mM sucrose, 50 mM KCl, and 0.2% (w/v) bovine serum albumin (BSA). Excised tissues were minced into small pieces using scissors and homogenized with a Potter–Elvehjem Dounce homogenizer. BAT tissues were homogenized with 30 strokes, whereas liver tissues were homogenized with 10 strokes. The homogenates were centrifuged at 700 × g for 10 min at 4 °C to remove nuclei and cellular debris. The resulting supernatants were collected and centrifuged at 7,000 × g for 10 min at 4 °C to pellet mitochondria. Mitochondrial pellets were resuspended in the corresponding isolation buffers and kept on ice. Mitochondrial protein concentrations were determined using the BCA protein assay kit (Pierce, Thermo Fisher Scientific). Isolated mitochondrial samples were snap-frozen in liquid nitrogen and stored at –80 °C until further analysis by liquid chromatography–mass spectrometry or western blotting.

### Mitochondria extraction for cells

Mitochondria were isolated from cultured cells based on a previously described protocol^63^, with minor modifications. Briefly, confluent cells were washed once with phosphate-buffered saline (PBS) and incubated for 30 min on ice in an isotonic buffer containing 100 mM sucrose, 10 mM MOPS (pH 7.2), 1 mM EGTA, and 0.1% (w/v) fatty acid–free bovine serum albumin (BSA; Sigma-Aldrich). Cells were then harvested and mechanically disrupted using a Potter–Elvehjem homogenizer with a PTFE pestle and glass tube (Sigma-Aldrich) for five strokes, followed by additional disruption by passing the suspension five times through a 25-gauge needle. The homogenate was centrifuged at 700 × g for 5 min at 4 °C to remove nuclei and unbroken cells, and the supernatant was collected. The pellet was subsequently resuspended in isolation buffer containing 210 mM mannitol, 70 mM sucrose, 10 mM MOPS (pH 7.2), 1 mM EGTA, and 0.1% fatty acid–free BSA, homogenized with five additional strokes, and passed through a 25-gauge needle five times. The resulting supernatant containing mitochondria was centrifuged at 12,000 × g for 5 min at 4 °C to pellet mitochondria.

### Acute BCAA and BCKA Tolerance Tests

BCAA and BCKA solutions were freshly prepared in sterile 0.9% saline. The BCAA solution contained 82.5 mM leucine, 37.5 mM isoleucine, and 60 mM valine, corresponding to a molar ratio of 2:1:1.5 (Leu: Ile: Val)^64^. The BCKA solution contained 5 mM α-ketoisocaproate (KIC), 5 mM α-keto-β-methylvalerate (KMV), and 5 mM α-ketoisovalerate (KIV), mixed at a 1:1:1 molar ratio. Control and SLC25A35-KO mice aged 10–14 weeks were fasted for 6 h (09:00–15:00) with free access to water. Following the fasting period, mice received an intraperitoneal injection of either the BCAA or BCKA solution at a volume of 8 µL per gram of body weight. Blood samples were collected via retro-orbital bleeding at 0, 30, and 60 min after injection. Blood samples were allowed to clot on ice for 30 min and then centrifuged at 10,000 × g for 10 min at 4 °C. The resulting serum was collected and subjected to liquid chromatography–mass spectrometry (LC–MS) analysis.

### Single-bolus ^13^C-Leu and ^13^C-KIC tracing following intraperitoneal injection

^13^C-labeled Leu and KIC tracer solutions were freshly prepared in sterile 0.9% saline. The ^13^C-Leu solution consisted of 41.25 mM uniformly ^13^C-labeled leucine (Leu, M+6), 18.7 mM unlabeled isoleucine, and 30 mM unlabeled valine, mixed at a molar ratio of 1:0.5:0.75 (^13^C-Leu (M+6): Ile: Val). The ^13^C-KIC solution contained 5 mM uniformly ^13^C-KIC (M+6), 5 mM unlabeled KMV, and 5 mM unlabeled KIV, mixed at a 1:1:1 molar ratio (^13^C-KIC (M+6): KMV: KIV). Control and SLC25A35-KO mice aged 10–14 weeks were fasted for 6 h (09:00–15:00) with free access to water, followed by a single intraperitoneal injection of either the ^13^C-KIC (M+6) or ^13^C-Leu (M+6) tracer solution at a volume of 8 µL per gram of body weight. For ^13^C-Leu (M+6) tracing, blood samples were collected via retro-orbital bleeding at 0, 5, 10, 20, and 30 min after injection. For ^13^C-KIC (M+6) tracing, blood samples were collected at 0, 3, 8, 15, and 25 min following injection. Blood samples were centrifuged at 10,000 × g for 10 min at 4 °C, and the resulting serum (for ^13^C-Leu (M+6) tracing) or serum (for ^13^C-KIC (M+6) tracing) was collected for LC–MS analysis.

### ^13^C-BCKA tracing in mature adipocytes

Uniformly ^13^C-labeled BCKA (200 µM; composition as indicated) were prepared in serum-free DMEM and added to mature adipocytes differentiated from control, SLC25A35-KO, and *Slc25a35*-rescued cells. Cells were incubated with the tracer-containing medium under standard culture conditions. Aliquots of culture medium (50 µL) were collected at 0, 1, 2, 5 or 6, 8, and 24 h after tracer addition. After 24 h of treatment, mature adipocytes were detached using TrypLE Express (Gibco, Thermo Fisher Scientific). Both culture medium and whole-cell samples were collected and processed for liquid chromatography-mass spectrometry (LC-MS) analysis.

### In vivo ^13^C-BCKA infusion via jugular vein catheterization

For ^13^C-BCKA infusion studies, SLC25A35-KO mice and paired control littermates aged 10–12 weeks were fasted overnight (19:00–09:00) with free access to water. Mice were anesthetized with isoflurane, and a catheter was surgically implanted into the jugular vein for tracer infusion^29^. Prior to tracer administration, approximately 50 µL of blood was collected via retro-orbital bleeding to obtain baseline samples. SLC25A35-KO mice and their paired control littermates were infused simultaneously with a ^13^C-BCKA tracer solution consisting of 25 mM ^13^C-KIC (M+6), ^13^C-KMV (M+6), and ^13^C-KIV (M+6) prepared in sterile saline. The tracer mixture was delivered through the jugular vein catheter at a constant infusion rate of 0.0836 µL per gram body weight per minute, corresponding to an individual tracer infusion rate of 25 nmol/min for each ^13^C-BCKA species. During the infusion, approximately 30 µL of blood was collected from the tail vein at 30, 90, and 120 min, placed on ice for 30 min, and centrifuged at 10,000 × g for 10 min at 4 °C to separate serum. At the conclusion of the experiment, mice were euthanized by cervical dislocation. Brown adipose tissue (BAT) and additional organs were rapidly excised, snap-frozen in liquid nitrogen, and stored at –80 °C until analysis. Serum and tissue samples were subsequently analyzed by LC-MS. All experiments were performed using one SLC25A35-KO mouse and one paired control mouse per day under identical experimental conditions. Data from Four independent pairs collected on separate days were combined for analysis.

### ^13^C-BCKA mitochondrial uptake assay

Mitochondria were isolated from mouse tissues or mature adipocytes as described above. Equal amounts of mitochondrial protein were resuspended in incubation buffer containing 210 mM mannitol, 70 mM sucrose, 10 mM MOPS, 1 mM EGTA, and 0.1% (w/v) BSA (pH 7.2). Mitochondrial suspensions were incubated with 50–100 µM uniformly ^13^C-labeled BCKA for 5 or 15 min at 37 °C with gentle agitation (500 rpm) using a thermomixer. At the indicated time points, mitochondria were pelleted by centrifugation at 12,000 × g for 5 min at 4 °C. The supernatant was carefully removed, and mitochondrial pellets were washed once with ice-cold incubation buffer followed by a second centrifugation at 12,000 × g for 5 min at 4 °C. The final mitochondrial pellets were snap-frozen in liquid nitrogen, stored at –80 °C,and subsequently analysed for intracellular metabolites by LC-MS.

### Untargeted metabolomics sample preparation and LC-MS analysis

**Sample preparation:** 20 µL plasma were extracted with 100 µL acetonitrile/methanol (1:1, v/v), vortexed for 1 min and centrifuged at 14,000 rpm for 8 min at 4°C. The resulting supernatant was dried under vacuum and reconstituted in 50% methanol. After centrifugation, the supernatant was injected for LC-MS analysis with injection volume of 2 µL.

**LC-MS analysis:** untargeted metabolomic analysis was performed using an ultra-performance liquid chromatography-high resolution mass spectrometry (UPLC-HRMS) platform. Metabolites were separated on a C18 column (150 x 4.6 mm, 5 µm, Waters, Milford) at a flow rate of 0.5 mL/min. Mobile phase A was 5 mM ammonium acetate and 0.1% ammonium hydroxide in water and mobile phase B was methanol. The column oven and autosampler tray were held at 30 °C and 6 °C respectively. The chromatographic gradient was as follows: 0-2 min: 2% B; 2-6 min: linear gradient from 2% B to 20% B; 6-8 min: linear gradient from 20% to 50%; 8-10 min: linear gradient from 50%-90%; 10-12 min: held at 90% B, followed by re-equilibration to 2% B. Mass spectrometry was performed on a Q-Exactive Plus orbitrap instrument (Thermo Fisher Scientific) equipped with an electrospray ionization (ESI) source and operated in full scan mode with ddMS2 monitoring in both positive and negative ionization modes. Instrument parameters were as follows: spray voltage +3.8 kV/–2.8 kV; capillary temperature 350 °C; sheath gas 55 arb; auxiliary gas 15 arb; probe heater temperature 350 °C; scan range 70–1000 m/z; resolution 70,000 for full scan and 15,000 for ddMS2 scan. Data acquisition was performed using Xcalibur 4.1 (Thermo Fisher Scientific). All samples were injected in a randomized order. A pooled quality control (QC) sample was prepared by combining 10 µL of supernatant from each sample and injected periodically throughout the batch to monitor instrument stability. Compound Discoverer v3.3 (Thermo Fisher Scientific) was used for peak alignment and extraction. Peak annotation was conducted with a 10 ppm mass tolerance for precursor and fragment ion by searching against an in-house and the online mzCloud library. Compound identification required 0.5 min tolerance of retention time to the authentic standards run under the same LC conditions and the MS2 spectral matching against spectra in the database. Only identified metabolites with peak area variation less than 30% in the QC sample were retained to generate a list including compound name, m/z, retention time and peak area. Then the peak areas were normalized to the median area across all detected metabolites in each sample to minimize the variability from sample preparation and instrument performance. The normalized value was used for subsequent statistical analysis.

### Targeted metabolomics for serum, total cell and isolated mitochondria

**Sample preparation:** Plasma, cell, and isolated mitochondria sample were stored at −80 °C until analysis. For plasma samples, after thawing on ice then the sample (10 µL) was transferred to a new tube and extracted with 100 µL acetonitrile: methanol (1:1, v/v) containing 5 µM internal standards. The mixtures were vortexed and centrifuged at 14,000 rpm for 10 min. Then the supernatants were transferred to a new tube and evaporated to dryness in vacuum concentrator under 4 °C. Dried extracts were reconstituted in 50 µL 5% acetonitrile in water, vortexed for 2 min, and centrifuged at 14,000 rpm for 10 min to remove the insoluble debris. The clarified supernatants were transferred to the glass vials for LC-MS injection. For cell and mitochondria sample, after thawing on ice then the sample were diluted with 100 µL water and extracted by 400 µL acetonitrile: methanol (1:1, v/v) containing internal standards. The mixtures were vortexed, sonicated for 10 min at 4 °C in a water bath and incubated at −80 °C for 30 min to facilitate protein precipitation. Samples were centrifuged at 14,000 rpm for 10 min and supernatants were dried under vacuum concentrator. Prior to LC-MS analysis, dried extracts were reconstituted in 100 µL 5% acetonitrile in water, centrifuged at 14,000 rpm for 10 min and transferred to glass vials. The injection volume was 10 µL.

**LC-MS analysis:** The LC conditions for the targeted assay were described in our previously published paper^12^. Mass spectrometry parameters were set as follows: heater temperature, 350 °C; sheath gas, 50; auxiliary gas, 15; spray voltage, 3 kV. Capillary temperature was set as 350 °C and S-lens was 50. Data were acquired under the mode of PRM mode with the resolution set at 15,000 (m/z 200) and the isolation window was set at 1.5 m/z. The AGC target value was set to 1×10^6^ with a maximum injection time of 100 ms. Raw data were collected using Xcalibur 4.1 and processed with TraceFinder 4.1.

### LC-MS analysis for Stable isotope tracing

For isotope tracing experiments, LC-MS analysis was performed using the same chromatographic conditions as the untargeted assay with minor modifications. Briefly, data were acquired in full scan mode over the m/z range of 70-1000 at the resolution of 140,000 (m/z 200). Data were processed using TraceFinder 4.1 to extract each isotopologue peak areas. Following raw data extraction, natural isotope abundances correction was performed using the AccuCor (https://github.com/XiaoyangSu/AccuCor).

### Acylcarnitine analysis

Acylcarnitine levels in brown adipose tissue (BAT) were quantified at the Duke–NUS Metabolomics Facility using established mass spectrometry–based methods for acylcarnitine extraction and profiling^13,65^. Briefly, BAT samples were homogenized in 50% acetonitrile containing 0.3% formic acid. For acylcarnitine extraction, 50 µL of tissue homogenate was mixed with methanol, followed by derivatization using 3 M hydrochloric acid in methanol or butanol (Sigma-Aldrich). After derivatization, samples were dried under vacuum and reconstituted in methanol dddddprior to analysis. Acylcarnitine measurements were performed by flow injection analysis coupled with tandem mass spectrometry (FIA–MS/MS) using an Agilent 1290 Infinity LC system (Agilent Technologies, CA, USA) interfaced with a QTRAP 5500 quadrupole–linear ion trap mass spectrometer (AB Sciex, DC, USA). Samples were analyzed at a flow rate of 0.4 mL/min using an 80:20 (v/v) methanol: water mobile phase, with an injection volume of 10 µL. Data acquisition and quantification were carried out using MultiQuant™ software (version 3.0.3; AB Sciex).

### RNA extraction and Quantitative RT-PCR

Total RNA was extracted from 20–40 mg of mouse tissues using TRIzol reagent (Invitrogen) according to the manufacturer’s instructions. One microgram of total RNA was reverse-transcribed into complementary DNA (cDNA) using a reverse transcription kit (Bio-Rad). Quantitative real-time PCR was performed using SYBR Green chemistry (Applied Biosystems, Bio-Rad) on a 7900HT Fast Real-Time PCR System (Applied Biosystems). Relative gene expression levels were calculated using the 2^–ΔΔCt method and normalized to 18S rRNA and GAPDH. Primer sequences are provided in Supplementary Table S1 and Table S2.

### Western blot analysis

Whole-cell or tissue lysates were prepared using RIPA buffer (Axil Scientific, First Base) supplemented with protease and phosphatase inhibitor cocktails (Roche). Protein concentrations were determined using the BCA protein assay kit (Pierce, Thermo Fisher Scientific). Equal amounts of protein (10 µg) were resolved by sodium dodecyl sulfate–polyacrylamide gel electrophoresis (SDS–PAGE) and transferred onto polyvinylidene difluoride (PVDF) membranes (Millipore). Membranes were blocked with 5% non-fat milk in Tris-buffered saline containing 0.1% Tween-20 (TBST) and subsequently incubated with the indicated primary antibodies: SLC25A35 (in-house generated and Novus Biologicals, NBP2-85742), TIM23 (Proteintech, 11123-1-AP), GAPDH (Cell Signaling Technology, 2118S), phospho-HSL (Ser563) (Cell Signaling Technology, 4139S), total HSL (Cell Signaling Technology, 4107S), ATGL (Cell Signaling Technology, 2439S), PKA regulatory subunit IIβ (Santa Cruz Biotechnology, sc-376778), DGAT2 (Santa Cruz Biotechnology, sc-293211), SCD1 (Santa Cruz Biotechnology, sc-58420), phospho-PRKAR2B (Ser114) (Thermo Fisher Scientific, PA5-37787), Fasn (Cell Signaling Technology, 3180), phospho-ACC (Thr389) (Cell Signaling Technology, 11818), ACC-Total (Cell Signaling Technology, 3676), Beta-actin (Cell Signaling Technology, 4967), phospho-ACYL (Cell Signaling Technology, 4331), ACYL-Total (Cell Signaling Technology, 4332), Lamin B1 (Santa Cruz Biotechnology, sc-374015), and total BCKDHA (Abcam, ab126173). Following incubation with the appropriate horseradish peroxidase (HRP)-conjugated secondary antibodies, immunoreactive bands were visualized using a ChemiDoc imaging system (Bio-Rad). Band intensities were quantified using ImageJ software.

### Haematoxylin-eosin staining and adipocyte size counting

Brown adipose tissue (BAT) was harvested from euthanized mice and fixed in 10% paraformaldehyde. Fixed tissues were dehydrated, cleared, and embedded in paraffin using standard histological procedures. Paraffin-embedded sections were stained with hematoxylin and eosin (H&E) to assess tissue morphology. All stained sections were imaged using an EVOS M7000 Imaging System (Thermo Fisher Scientific, Pittsburgh, PA). For adipocyte size analysis, representative images were acquired from each mouse (n = 5 per group). Adipocyte cross-sectional area was quantified using AdipoCount software.

### Serum Non-Esterified Fatty Acid (NEFA) measurement

Serum NEFA levels were measured using the LabAssay™ NEFA kit (LabAssay NEFA FUJIFILM 633-52001, FUJIFILM) according to the manufacturer’s instructions. Briefly, 4 µL of serum from wild-type and SLC25A35-KO mice following 24 h of cold exposure was incubated with color reagent A at 37 °C for 10 min, followed by addition of color reagent B and further incubation at 37 °C for 10 min. The reaction generates a purple-colored product proportional to NEFA concentration. Absorbance was measured at 550 nm using a microplate reader, and NEFA concentrations were calculated based on a standard curve generated from the provided standards.

### Seahorse assay in mature adipocytes

Wild-type and SLC25A35-KO derived pre-adipocytes 8 x 10^3^ were first coated onto Seahorse XF96 Cell Culture microplates, followed by induction of maturation for 6 days. Mitochondria oxygen consumption rates (OCR) was measured in mature adipocytes using Seahorse Bioscience XFe-96 Analyzer (Agilent). Seahorse Mito stress kit (Agilent) using 2 µM Oligomycin, 2.5 µM FCCP and 0.5 µM rotenone/antimycin injected according to Seahorse Mito stress Assay Protocol (Agilent).

### Isolated mitochondria Seahorse assay

For assays using isolated mitochondria^66^, mitochondria were freshly isolated from brown adipose tissue (BAT) of wild-type (WT) and SLC25A35-KO mice. A total of 14 µg mitochondrial protein per well was plated onto Seahorse XF96 microplates in mitochondrial assay buffer. Plates were centrifuged at 2,000 × g for 20 min at 4 °C to facilitate mitochondrial adherence. The mitochondrial assay buffer consisted of 70 mM sucrose, 220 mM mannitol, 10 mM KH,PO4, 5 mM MgCl,, 2 mM HEPES, 1 mM EGTA, and 0.4% (w/v) fatty acid–free BSA. Pyruvate and sodium malate were supplied as respiratory substrates for the isolated mitochondrial Seahorse assay. Mitochondrial respiration was assessed using the Seahorse XF Mito Stress Test protocol with sequential injections of ADP (3 mM), oligomycin (3 µM), FCCP (4 µM), and antimycin A (4 µM). Oxygen consumption rate (OCR) values were normalized to mitochondrial protein content, as indicated in the figure legends.

### Immunofluorescence staining for subcellular localization

HUH7 cells (1 × 10^5^) were seeded onto 8-well glass chamber slides (BioTek, Invitrogen) and cultured for 24 h. Cells were then transfected with a pcDNA3.1 expression vector encoding human *SLC25A35* fused in-frame with a FLAG tag using Lipofectamine 2000 (Life Technologies), according to the manufacturer’s instructions. After 18 h of transfection, cells were washed once with PBS and fixed with 4% formaldehyde, followed by two washes with PBS containing 0.1% Tween-20 (PBST). Fixed cells were blocked with 10% normal goat serum (Invitrogen) for 1 h at room temperature and subsequently washed twice with PBST. Cells were then incubated with primary antibodies against FLAG (mouse anti-FLAG, 1:200; Sigma-Aldrich) and the mitochondrial inner membrane protein TIM23 (rabbit anti-TIM23, 1:200; Proteintech). Following two washes with PBST, cells were incubated with species-specific secondary antibodies conjugated to Alexa Fluor 488 (anti-mouse) or Alexa Fluor 547 (anti-rabbit) to visualize SLC25A35 and TIM23, respectively. Nuclei were counterstained with Hoechst 33342 (Life Technologies). Fluorescent images were acquired using a Zeiss confocal microscope.

### RNAseq data analysis for ad-libitum, fasting and refeeding

Gene expression and GElaS (Gene Elastic Score) in major metabolic organs during the ad libitum– fasting–refeeding (AL–F–R) cycle were obtained from a previous study^33^, and the data were visualized using the R package ggplot2 (v3.5.2, https://ggplot2.tidyverse.org/). Gene expression was scaled by mean and variance for each Slc25 gene. GElaS values were ranked based on the score for the *Slc25* gene family.

### Luciferase reporter assay for the SLC25A35 promoter

AML12 cells were seeded into 12-well plates and cultured for 18–24 h prior to transfection. Each well was co-transfected with 1 µg of a pGL3.1 (Promega) firefly luciferase reporter construct containing 1,000 bp of the genomic sequence upstream of the SLC25A35 5’ untranslated region (5’UTR), together with 20 ng of pRL Renilla luciferase plasmid as an internal control, using X-tremeGENE 9 DNA transfection reagent (Roche) according to the manufacturer’s instructions. Twenty-four hours after transfection, culture medium was replaced with either high-glucose or glucose-free DMEM, with or without 10% fetal bovine serum (FBS), and supplemented as indicated with oleic acid and/or palmitic acid at the specified concentrations for the indicated durations. Cells were then harvested, and luciferase activity was measured using the Dual-Luciferase Reporter Assay System (Promega, Madison, WI) following the manufacturer’s protocol. Luminescence was quantified using a Tecan Spark 10M microplate reader. Firefly luciferase activity was normalized to Renilla luciferase activity and expressed as a firefly/Renilla luminescence ratio.

## QUANTIFICATION AND STATISTICAL ANALYSIS

All experimental data were analyzed using GraphPad Prism version 10.1.2 and are presented as mean ± SEM. Data normality was assessed using the Shapiro–Wilk test where applicable. For normally distributed data, parametric statistical tests were applied. Comparisons between two groups were performed using unpaired two-tailed Student’s t tests. Multiple unpaired t tests with false discovery rate (FDR) correction were applied as appropriate. Comparisons among more than two groups were analyzed using one-way analysis of variance (ANOVA), followed by Dunnett’s multiple comparisons test for comparisons with a control group, as specified in the corresponding figure legends. For experiments involving two independent variables (e.g., time and genotype or treatment), two-way ANOVA was performed, followed by Tukey’s multiple comparisons test, as indicated. A P value < 0.05 was considered statistically significant. Experimental schematics were generated using BioRender.

## SUPPLEMENTARY FIGURE LEGENDS

**Figure S1**

(A) Gene elasticity score (GElaS) against the absolute log_2_ fold change on fasting (fasted vs ad libitum) for SLC25 family genes in epididymal white adipose tissue (eWAT), liver and skeletal muscle of young mice.

**Figure S2**

(A) Relative *Slc25a35* mRNA expression in BAT, iWAT, gastrocnemius muscle (Gastro), heart and liver of control mice under ad libitum feeding, fasting, and refeeding, normalized to the ad libitum condition (n = 4 per group).

(B) Relative *Slc25a35* mRNA expression in eWAT, kidney, soleus muscle (Soleus), lung, spleen, uterus, intestine, brain and cerebellum (Cereb) of control mice under ad libitum feeding, fasting, and refeeding, normalized to the ad libitum condition (n = 4 per group).

(C) Immunoblot (left) and quantification (right) of SLC25A35 in mitochondria isolated from the indicated mouse tissues, normalized to Tim23. GAPDH is used as quality control of isolation.

(D) SLC25A35 protein abundance in cytosol and mitochondria.

(E) *Slc25a35* mRNA levels in wild-type preadipocytes cultured in high-glucose medium (HGM) with or without 10% fetal bovine serum (FBS) (n = 6 per group).

(F) Relative *Slc25a35* mRNA levels in AML12 cells treated with HGM alone, HGM supplemented with 10% FBS for 24 h, or HGM for 12 h followed by HGM + 10% FBS for 12 h (n = 3 per group).

(G) Immunoblot analysis of SLC25A35 in wild-type preadipocytes cultured in high-glucose DMEM with or without 10% FBS (left panel). Quantification of SLC25A35 protein levels normalized to GAPDH is shown in the right panel (n = 3 per group).

(H) Immunoblot analysis of SLC25A35 protein levels in AML12 cells under the conditions shown in (F), with β-actin used as a loading control. Quantification of SLC25A35 protein abundance normalized to β-actin is shown in the right panel (n = 3 per group).

(I–J) Differences in transcription factor activity between nutritional states in the subcutaneous white adipose tissue (sWAT) (I) and epididymal white adipose tissue (eWAT) (J) transcriptomes: ad libitum versus fasted and fasted versus refed. The x axes show Cohen’s d and the y axes show –log_10_ of the Benjamini–Hochberg-adjusted P value (dashed lines, adjusted P = 0.05).

(K) *Srebf1* mRNA levels in brown preadipocytes transfected with control siRNA (si-NC) under fed or fasting-mimicking conditions or with SREBP1 siRNA (si-SREBP1) under fasting-mimicking conditions (n = 4 per group).

(L) Relative *Slc25a35* mRNA levels in AML12 cells cultured in complete medium (10% FBS) under low-glucose (1 g/L) or high-glucose (4.5 g/L) conditions (n = 6 per group).

(M) Relative luciferase activity of the SLC25A35 promoter reporter normalized to Renilla luciferase in complete medium (10% FBS) under low-glucose (1 g/L) or high-glucose (4.5 g/L) conditions (n = 3 per group).

(N) Immunoblot analysis of SLC25A35 protein levels in AML12 cells exposed to increasing glucose concentrations in the absence (left) or presence (right) of FBS. β-actin was used as a loading control.

(O) Relative *Slc25a35* mRNA levels in AML12 cells cultured in high-glucose medium with 10% FBS in the absence or presence of 0.5 mM free fatty acids (palmitic acid and oleic acid) (n = 3 per group).

(P) Relative luciferase activity of the SLC25A35 promoter reporter normalized to Renilla luciferase in high-glucose medium with (left) or without (right) 10% FBS in the absence or presence of free fatty acids (palmitic acid and oleic acid; 450 µM with FBS and 500 µM without FBS) (n = 3 per group).

(Q) Immunoblot analysis (left) and quantification (right) of SLC25A35 protein levels in brown preadipocytes cultured in high-glucose medium (HGM) with 10% FBS, in HGM alone, or in HGM supplemented with 300 or 450 µM palmitic acid and oleic acid (PA+OA). GAPDH was used as a loading control. Quantification of SLC25A35 protein abundance is normalized to GAPDH (n = 3 per group).

Data are presented as mean ± SEM. P values in panels (A, B, F, H and Q) were determined by ordinary one-way ANOVA followed by Dunnett’s multiple comparisons test. P values in panels (E, G, L, M, O and P) were determined by unpaired two-tailed Student’s t tests. P values in panel (K) were determined by one-way ANOVA followed by multiple comparisons test.

**Figure S3**

(A) Schematic illustrating generation of *Slc25a35* global knockout mice, achieved by Zp3-Cre–mediated deletion of *Slc25a35* exons 2–5 flanked by loxP sites during early embryogenesis.

(B) Relative *Slc25a35* mRNA expression across multiple organs from control and SLC25A35-KO mice (left; n = 6 per group) and quantification of SLC25A35 protein levels normalized to the inner mitochondrial membrane protein TIM23 (right; n = 3 per group). Gastro, gastrocnemius muscle; Quads, quadriceps muscle.

(C) Immunoblot analysis of SLC25A35 protein in mitochondria isolated from brown adipose tissue (BAT), heart, liver, quadriceps muscle, and kidney (n = 3 per group).

(D) Body weight, lean mass, and fat mass in control and SLC25A35-KO mice under basal conditions (n = 5 per group).

(E) Food and water intake normalized to body weight in control and SLC25A35-KO mice under basal conditions (n = 3 per group).

(F) Glucose tolerance test (GTT) in control and SLC25A35-KO mice following intraperitoneal glucose injection. Blood glucose levels at the indicated time points are shown (n = 7 control; n = 9 SLC25A35-KO).

(G) Weights of major organs from control and SLC25A35-KO mice under basal condition (n = 5 per group).

(H) Experimental schematic illustrating serum collection from control and SLC25A35-KO mice 2 h after refeeding following an overnight fast.

(I) Principal component analysis (PCA) of serum metabolite profiles from control and SLC25A35-KO mice under refeeding conditions.

(J) Heatmap of selected serum metabolites from individual control and SLC25A35-KO mice in the refed state. Values are shown as row z-scores.

(K) Experimental schematic illustrating serum collection from control and SLC25A35-KO mice under fasting conditions for untargeted metabolomic analysis.

(L) Principal component analysis (PCA) of serum metabolite profiles from control and SLC25A35-KO mice, showing clear separation between genotypes. Percent variance explained by each principal component is indicated.

(M) Heatmap of serum metabolite abundance from individual control and SLC25A35-KO mice under fasting condition. Values are shown as row z-scores.

(N) Volcano plot of serum metabolites comparing SLC25A35-KO and control mice under fasting condition. Red, significantly upregulated (Up) and cyan, significantly downregulated (Down) metabolites in SLC25A35-KO mice; dashed lines denote thresholds of |log_2_ fold change| > 0.585 (1.5-fold) and –log_10_ (P value) > 1.3 (P < 0.05).

Data are presented as mean ± SEM. P values in panels (B, D, E and G) were determined by multiple unpaired t tests with false discovery rate (FDR) correction. P values in panel (F) were determined by one-way ANOVA followed by Dunnett’s multiple comparisons test.

**Figure S4**

(A) Experimental schematic of the BCKA tolerance test, in which control and SLC25A35-KO mice were fasted for 6 h (09:00–15:00) and intraperitoneally injected with a bolus containing ^13^C-KIC (M+6) together with unlabeled KMV and KIV. Serum was collected at the indicated time points (0, 3, 8, 15, and 25 min).

(B) Serum levels of unlabeled BCKA (M+0)—KIC, KMV and KIV—in control and SLC25A35-KO mice during the BCKA tolerance test (n = 4 per group).

(C) Serum levels of the unlabeled BCKA-derived metabolites HIC (M+0) and HMV (M+0) in control and SLC25A35-KO mice during the BCKA tolerance test (n = 4 per group).

(D) Serum levels of unlabeled BCAA (M+0)—Leu, Ile and Val—in control and SLC25A35-KO mice during the BCKA tolerance test (n = 4 per group).

(E) Schematic illustrating the metabolic fate of ^13^C-labeled BCKA carbons, including mitochondrial transamination to BCAA and cytosolic reduction to hydroxy-BCKA derivatives.

(F) Experimental schematic of the BCAA tolerance test, in which control and SLC25A35-KO mice were fasted for 6 h (09:00–15:00) and intraperitoneally injected with a bolus containing ^13^C-labeled Leu together with unlabeled Ile and Val. Serum was collected at the indicated time points (0, 5, 10, 20, and 30 min).

(G) Serum levels of unlabeled BCKA (M+0)—KIC, KMV and KIV—in control and SLC25A35-KO mice during the BCAA tolerance test (n = 3 per group).

(H) Serum levels of unlabeled HIC (M+0) in control and SLC25A35-KO mice during the BCAA tolerance test (n = 3 per group).

(I) Serum levels of unlabeled Ile (M+0) and Val (M+0) in control and SLC25A35-KO mice during the BCAA tolerance test (n = 3 per group).

(J) Schematic illustrating the metabolic fate of ^13^C-labeled BCAA carbons, including mitochondrial transamination to BCKA and cytosolic reduction of BCKA to hydroxy-BCKA derivatives.

Data are presented as mean ± SEM. P values in panels (B–D and G–I) were determined by two-way ANOVA followed by Tukey’s multiple comparisons test.

**Figure S5**

(A) Experimental schematic of in vivo steady-state intravenous infusion of uniformly ^13^C-labeled BCKA in control and SLC25A35-KO mice, with serum collected at 0, 30, 90, and 120 min for metabolite analysis.

(B) Serum levels of ^13^C-labeled BCKA (upper row)—^13^C-KIC (M+6), ^13^C-KMV (M+6) and ^13^C-KIV (M+5)—and of the corresponding unlabeled BCKA (lower row)—KIC (M+0), KMV (M+0) and KIV (M+0)—measured at the indicated time points in control and SLC25A35-KO mice during steady-state infusion (signal intensity normalized to the mean; n = 5 per group).

(C) Serum levels of the ^13^C-labeled downstream metabolites ^13^C-HIC (M+6) and ^13^C-HMV (M+6) (left two plots; n = 5 per group) and of unlabeled HIC (M+0) and HMV (M+0) (right two plots; n = 6 per group), measured at the indicated time points in control and SLC25A35-KO mice during steady-state infusion (signal intensity normalized to the mean).

(D) Serum levels of ^13^C-labeled BCAA (upper row)—^13^C-Leu (M+6), ^13^C-Ile (M+6) and ^13^C-Val (M+5)— and of unlabeled BCAA (lower row)—Leu (M+0), Ile (M+0) and Val (M+0)—measured at the indicated time points in control and SLC25A35-KO mice (signal intensity normalized to the mean; n = 6 per group).

(E) Experimental schematic of medium collection, illustrating sampling at 0, 1, 2, 5, 8, and 24 h from mature adipocytes differentiated from control and SLC25A35-KO preadipocytes.

(F–I) Medium levels of unlabeled BCKA (M+0)—KIC (F), KMV (G) and KIV (H)—and the downstream metabolite 3-hydroxyisovalerate (M+0) (I) in mature adipocytes differentiated from control and SLC25A35-KO preadipocytes (n = 5 per group).

(J–L) Medium levels of ^13^C-labeled BCAA—Leu (M+6) (J), Ile (M+6) (K) and Val (M+5) (L)—in mature adipocytes differentiated from control and SLC25A35-KO preadipocytes (n = 5 per group).

Data are presented as mean ± SEM. P values in panels (B–D and F–L) were determined by two-way ANOVA followed by Tukey’s multiple comparisons test. In panels (F–I), the blue P values indicate the significance of differences between SLC25A35-KO and control groups without ^13^C-labeled BCKA treatment, whereas the black P values indicate the significance of differences between SLC25A35-KO and control groups with ^13^C-labeled BCKA treatment.

**Figure S6**

(A) Experimental schematic of the BCKA tolerance test, in which SLC25A35-KO mice and SLC25A35-KO mice with systemic re-expression of SLC25A35 (AAV9-CMV-Slc25a35) were fasted for 6 h (09:00– 15:00) and received an IP bolus of unlabeled KIC, KMV and KIV, followed by blood collection after 8 min.

(B–D) Serum levels of KIC, KMV and KIV (B), HIC, HMV and HIV (C) and Leu, Ile and Val (D) 8 min after the BCKA bolus in SLC25A35-KO mice and SLC25A35-KO mice re-expressing SLC25A35 (AAV9-CMV-Slc25a35) (n = 7 SLC25A35-KO; n = 8 SLC25A35-KO with AAV9-CMV-Slc25a35).

(E) Immunoblot analysis of SLC25A35 in the liver of control, SLC25A35-KO and SLC25A35-KO (AAV8-Alb-Slc25a35) mice. Tim23 was used as a loading control.

(F) Schematic illustrating the generation of liver-specific *Bckdk* knockout (BCKDK-LKO) mice by Alb-Cre–mediated deletion of the floxed *Bckdk* allele.

(G) Relative *Bckdk* mRNA levels (top; n = 7 per group) and immunoblot analysis of BCKDK (bottom) in the liver of control and BCKDK-LKO mice. GAPDH was used as a loading control.

(H–I) Serum levels of BCKA (H) and BCAA (I) in control and BCKDK-LKO mice (n = 4 control; n = 8 BCKDK-LKO).

(J) Experimental schematic illustrating overexpression of *Slc25a35* in SLC25A35-KO preadipocytes.

(K) Immunoblot analysis of SLC25A35 protein levels in control, SLC25A35-KO and SLC25A35-rescued (hSLC25A35) preadipocytes (left; Tim23 and vinculin, loading controls) and relative *Slc25a35* mRNA expression in SLC25A35-KO and SLC25A35-rescued preadipocytes (right; n = 3 per group).

(L) Experimental schematic of the rescue experiment, illustrating medium collection after 6 and 24 h from mature adipocytes differentiated from SLC25A35-KO and SLC25A35-rescued preadipocytes following incubation with uniformly ^13^C-labeled BCKA.

(M–N) Medium levels of unlabeled metabolites, including KIC (M+0), KMV (M+0) and KIV (M+0) (M) and Leu (M+0), Ile (M+0) and Val (M+0) (N), measured in SLC25A35-KO and SLC25A35-rescued mature adipocytes following 6 h and 24 h incubation with uniformly ^13^C-labeled BCKA (n = 5 per group).

(O) Mitochondrial levels of ^13^C-Leu (M+6), ^13^C-Ile (M+6) and ^13^C-Val (M+5) at 0, 5, 15 and 30 min after incubation of BAT mitochondria from control and SLC25A35-KO mice with 100 µM ^13^C-BCKA.

Data are presented as mean ± SEM. P values in panels (B–D and H–I) were determined by multiple unpaired t tests with false discovery rate (FDR) correction. P values in panels (G and K) were determined by unpaired two-tailed Student’s t tests. P values in panels (M and N) were determined by one-way ANOVA followed by Dunnett’s multiple comparisons test. P values in panel (O) were determined by two-way ANOVA followed by Tukey’s multiple comparisons test.

**Figure S7**

(A) Schematic representation of the genomic loci of *Slc25a35* and *Slc25a44* illustrating the generation of global SLC25A35/A44-DKO mice.

(B) Experimental schematic illustrating serum collection from control, SLC25A35-KO, SLC25A44-KO, and SLC25A35/A44-DKO mice under overnight fasting and 2-h refeeding conditions.

(C–D) Serum levels of BCKA measured in control, SLC25A35-KO, SLC25A44-KO, and SLC25A35/A44-DKO mice following 2 h of refeeding (C) or overnight fasting (D) (n = 6 per group in control and SLC25A35-KO; n = 5 in SLC25A44-KO and SLC25A35/A44-DKO).

(E–F) Serum levels of BCAA measured in control, SLC25A35-KO, SLC25A44-KO, and SLC25A35/A44-DKO mice following 2 h of refeeding (E) or overnight fasting (F) (n = 6 control and SLC25A35-KO; n = 5 SLC25A44-KO and SLC25A35/A44-DKO).

Data are presented as mean ± SEM. P values in panels (C–F) were determined by multiple unpaired t tests with false discovery rate (FDR) correction.

**Figure S8**

(A) Gene tree of SLC25A35 homologs, with collapsed clades indicated, the yeast homolog OAC1 included, and human SLC25A35 highlighted in red.

(B) Multiple sequence alignment of SLC25A35 orthologs from human, dog, cow, mouse, rat and zebrafish.

(C) Predicted structure of human SLC25A35 (cartoon, light blue) with Gln73, Arg175, Arg232, Gly262 and Arg276 shown as sticks. Residues are numbered as in human SLC25A35 (UniProt Q3KQZ1, 300 residues).

(D) Design of the SLC25A35 variants. Candidate residues were selected from ligand-binding, conservation and human-variant evidence: R175A (Arg to Ala; KIC-binding site), R232A (Arg to Ala; conserved site), G262A (Gly to Ala; SNP-associated site) and Q73A/R276A (Gln to Ala and Arg to Ala; KIC-binding sites).

**Figure S9**

(A) Fasting- and refeeding-induced transcriptomic changes in subcutaneous (inguinal) white adipose tissue. Genes that are highly affected by both fasting and refeeding are highlighted by gene symbols.

(B) Schematic illustrating the experimental workflow for isolation of preadipocytes from the stromal vascular fraction (SVF), followed by in vitro differentiation into mature adipocytes and Oil Red O staining (lower panel).

(C) Relative *Slc25a35* mRNA expression (left) and immunoblot analysis of SLC25A35 and BCKDHA protein levels (right) during adipocyte differentiation at days 0, 2, 4, and 6, with GAPDH serving as a loading control.

(D–G) Relative mRNA expression of *Pdgfra* (D), *Slc25a35* (E), *Pck1* (F) and *Pck2* (G) in preadipocytes and mature adipocytes differentiated from the stromal vascular fraction (n = 5–6 per group).

(H) Knockdown efficiency of *Pck2* (left) and *Pck1* (right) in cells transfected with control siRNA (si-NC) or the respective siRNA (n = 3 per group).

(I) Oil Red O staining of control adipocytes, SLC25A35-overexpressing (SLC25A35-OV) adipocytes transfected with control siRNA (siNC), and SLC25A35-OV adipocytes transfected with *Pck2* siRNA (siPCK2).

(J) Heatmap of the fold change in the expression of the indicated genes (RT-qPCR) in SLC25A35-OV versus control cells (left) and in siPCK2-versus si-NC-transfected cells (right). Numbers show the fold change and the P value.

Data are presented as mean ± SEM. P values in panels (D–H) were determined by unpaired two-tailed Student’s t tests. P values in panel (J) were determined by unpaired two-tailed Student’s t tests.

**Figure S10**

(A) Experimental schematic illustrating control and SLC25A35-KO mice were housed at thermoneutrality for 2 weeks following a defined pre-cold acclimation period.

(B) Body weight, lean mass, and fat mass in control and SLC25A35-KO mice under thermoneutral conditions (n = 5 per group).

(C) The percentage of fat mass and lean mass in control and SLC25A35-KO mice under thermoneutral conditions (n = 5 per group).

(D) Weights of major organs from control and SLC25A35-KO mice under thermoneutral condition (n = 5 per group).

(E) Representative hematoxylin and eosin (H&E)–stained sections of BAT from control and SLC25A35-KO mice under thermoneutral condition. Scale bars, 50 µm. (right panel) Quantification of adipocyte lipid droplet area from BAT. Analysis was performed using three histological sections per mouse, with entire fields quantified per section (n = 5 per group).

(F) Relative *Slc25a35* mRNA levels in the indicated tissues of mice that were fed at room temperature (Fed+RT) or fed and exposed to cold (Fed+Cold) (n = 4 per group).

(G) Relative *Slc25a35* mRNA levels in the indicated tissues of mice that were fasted at room temperature (Fast+RT) or fasted and exposed to cold (Fast+Cold) (n = 4 per group).

(H) Immunoblot analysis (left) and quantification (right) of SLC25A35 protein abundance in BAT of mice under the indicated conditions (Fed+TN, Fast+TN, Fed+Cold, Fast+Cold). Tim23 was used as a loading control (n = 3 per group).

(I) Experimental schematic illustrating the cold-challenge protocol, in which control and SLC25A35-KO mice were housed at thermoneutrality for 2 weeks, subjected to a defined pre-cold acclimation period, and then exposed to an acute 24 h cold challenge (4 °C).

(J) Lean mass (left) and body weight (right) of control and SLC25A35-KO mice before (pre-cold) and after (post-cold) cold challenge for 24 h (n = 12 control; n = 14 SLC25A35-KO).

(K) Weights of heart, liver, kidney and skeletal muscle (SKM) of control and SLC25A35-KO mice after cold challenge for 24 h (n = 12 control; n = 14 SLC25A35-KO).

(L) Serum non-esterified fatty acid (NEFA) levels in control and SLC25A35-KO mice after 24 h of cold exposure.

(M) Seahorse analysis of oxygen consumption rates (OCR) in mature brown adipocytes differentiated from BAT-derived preadipocytes isolated from control and SLC25A35-KO mice and stimulated with norepinephrine (NE). Basal respiration, ATP production and maximal respiratory capacity are shown (n = 6–7 control; n = 8 SLC25A35-KO).

Data are presented as mean ± SEM. P values in panels (B–D) were determined by multiple unpaired t tests with false discovery rate (FDR) correction. The P value in panel (E) was determined by an unpaired two-tailed Student’s t test. P values in panels (F–H) were determined by multiple unpaired t tests with false discovery rate (FDR) correction. P values in panel (J) were determined by one-way ANOVA followed by Dunnett’s multiple comparisons test. P values in panel (K) were determined by multiple unpaired t tests with false discovery rate (FDR) correction. The P value in panel (L) was determined by an unpaired two-tailed Student’s t test. P values in panel (M) were determined by unpaired two-tailed Student’s t tests.

**Figure S11**

(A) Principal component analysis of the BAT transcriptome (500 most variable genes, variance-stabilized counts) of wild-type (WT) and SLC25A35-KO (KO) mice at thermoneutrality (TN) and in the cold (n = 3 WT and 4 KO mice at TN; 6 WT and 6 KO mice in the cold). PC1 and PC2 explain 51% and 36% of the variance.

(B–C) Volcano plots of KO versus WT at TN (B) and in the cold (C) on shared axes. Colored points are differentially expressed genes (DEGs; red, up in KO; blue, down in KO); numbers give DEG counts; dashed lines mark ±1.5-fold and the P value corresponding to FDR = 0.05.

(D) The 44 genes of the MitoCarta3.0 SLC25A family. Dots, from the outside: one ring per temperature (outer, TN; inner, cold); dot color, shrunken log_2_ fold change (KO/WT; capped at ±1); dot size, –log_10_ Wald P on one scale for both temperatures (capped at P = 10^−10^); black outline, FDR < 0.05; square, P not available. *Slc25a35* is labeled in red.

(E) Hallmark gene sets enriched in KO versus WT BAT at TN (GSEA NES; color, FDR; size, number of core-enrichment genes).

(F) Running enrichment scores (top) and ranked Wald statistic (bottom) of the MitoCarta3.0 OXPHOS subunit, mitochondrial ribosome and complex I subunit gene sets for KO versus WT BAT at TN.

(G) Heatmap of the relative expression (RT–qPCR) of genes involved in adipogenic regulation, BCAA catabolism, lipid synthesis and uptake, fatty acid oxidation, lipolysis, glucose metabolism and thermogenesis in BAT from control and SLC25A35-KO mice maintained at thermoneutrality (30 °C) for 2 weeks or after cold challenge (4 °C) for 24 h (n = 5 per group).

(H) Immunoblot analysis (top) and corresponding quantification (bottom) of lipogenesis-associated proteins in BAT of control and SLC25A35-KO mice following cold exposure (4 °C). Protein levels were normalized to GAPDH (n = 6 per group).

(I) Immunoblot analysis (left) and corresponding quantification (right) of lipolysis-associated proteins in BAT of control and SLC25A35-KO mice following cold exposure (4 °C). Protein levels were normalized to GAPDH (n = 6 per group).

Data are presented as mean ± SEM. P values in panels (H and I) were determined by multiple unpaired t tests with false discovery rate (FDR) correction. Differential expression and gene-set statistics in panels (A–F) are described in the STAR Methods.

**Figure S12**

(A) Experimental schematic. Control and SLC25A35-KO mice housed at thermoneutrality for 2 weeks were fasted for 6 h at 30 °C or 4 °C, and serum and BAT were collected.

(B) Heatmap of serum KIC, KMV, KIV, HIC, HMV, HIV, Leu, Ile, Val, acetoacetate, 3-hydroxybutyrate (3-HB), 3-aminoisobutyric acid, 3-hydroxy-3-methylglutarate (3-HMG), 3-hydroxyisobutyrate (3-HIB) and 3-hydroxy-3-methylbutyrate (HMB) in control and SLC25A35-KO (A35-KO) mice at 30 °C and 4 °C (row z-scores). Numbers show group means; P values are for the comparison between SLC25A35-KO and control mice at the same temperature.

(C) Heatmap of BAT KIC, KMV, HIC, HMV, HIV, Leu, Ile, Val, acetoacetate, 3-HB, 3-HMG, 3-aminoisobutyric acid, 3-HIB and HMB in control and SLC25A35-KO (A35-KO) mice at 30 °C and 4 °C (row z-scores). Numbers show group means; P values are for the comparison between SLC25A35-KO and control mice at the same temperature.

Data are presented as mean ± SEM. P values were determined by two-way ANOVA followed by multiple comparisons test.

