## Supplementary material for "Branched-Chain α-Keto Acid Mitochondrial Uptake Regulates Fatty Acid Metabolism in Adipose Tissue": Supplementary FIgures.pdf

### Figure S1

**A**

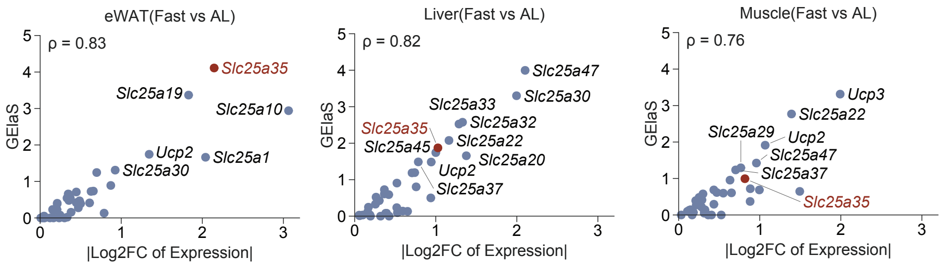
eWAT(Fast vs AL)
ρ = 0.83
Slc25a35
Slc25a19
Slc25a10
Ucp2
Slc25a1
Slc25a30
GElaS
|Log2FC of Expression|
Liver(Fast vs AL)
ρ = 0.82
Slc25a47
Slc25a30
Slc25a33
Slc25a32
Slc25a35
Slc25a22
Slc25a45
Slc25a20
Ucp2
Slc25a37
Muscle(Fast vs AL)
ρ = 0.76
Ucp3
Slc25a22
Slc25a29
Ucp2
Slc25a47
Slc25a37
Slc25a35

### Figure S2

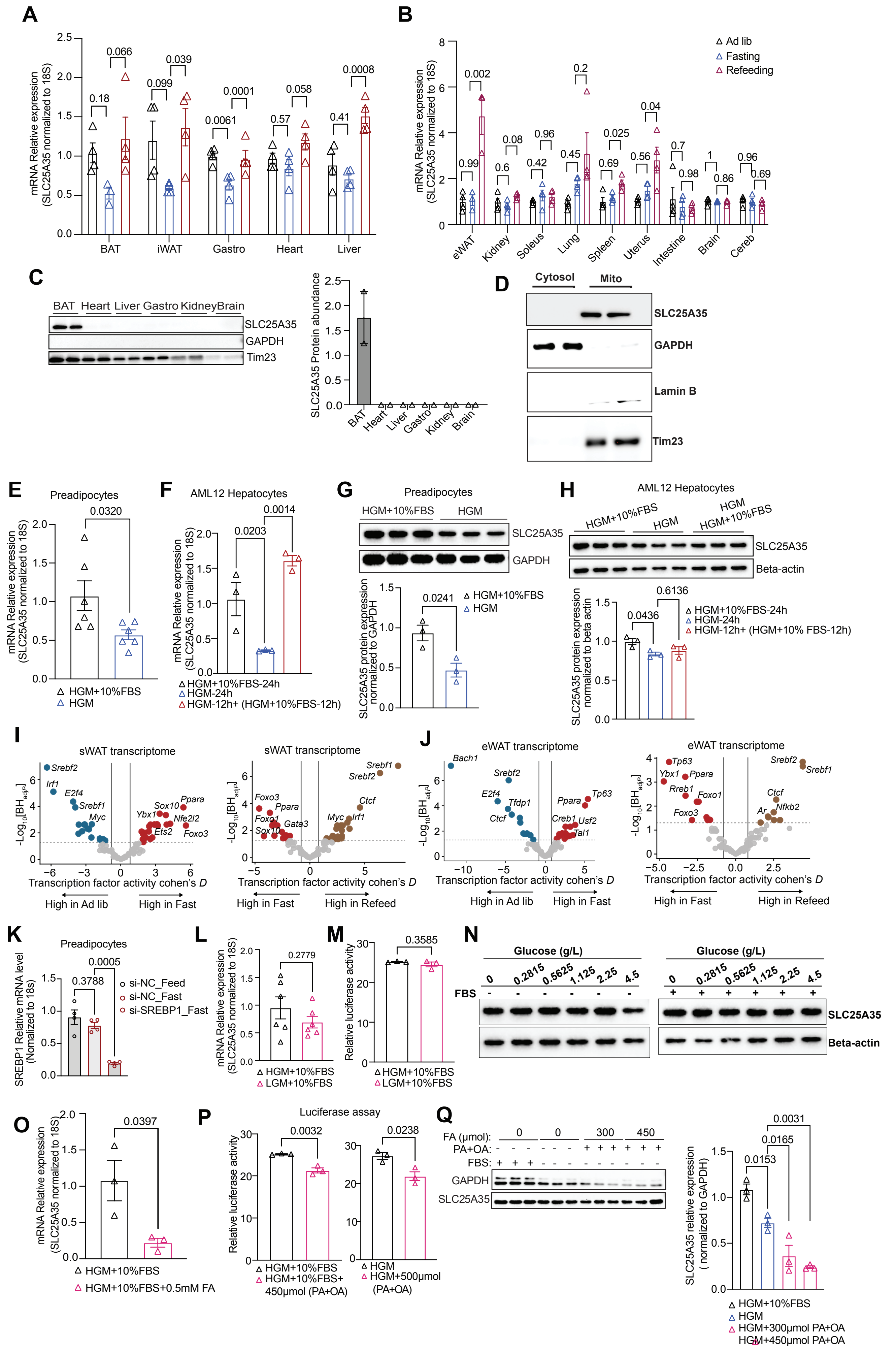
A
mRNA Relative expression (SLC25A35 normalized to 18S)
BAT
iWAT
Gastro
Heart
Liver
B
Ad lib
Fasting
Refeeding
eWAT
Kidney
Soleus
Lung
Spleen
Uterus
Intestine
Brain
Cereb
C
BAT Heart Liver Gastro Kidney Brain
SLC25A35
GAPDH
Tim23
SLC25A35 Protein abundance
D
Cytosol
Mito
SLC25A35
GAPDH
Lamin B
Tim23
E
Preadipocytes
HGM+10%FBS
HGM
F
AML12 Hepatocytes
HGM+10%FBS-24h
HGM-24h
HGM-12h+ (HGM+10%FBS-12h)
G
Preadipocytes
HGM+10%FBS
HGM
SLC25A35
GAPDH
SLC25A35 protein expression normalized to GAPDH
H
AML12 Hepatocytes
HGM+10%FBS
HGM
HGM HGM+10%FBS
SLC25A35
Beta-actin
SLC25A35 protein expression normalized to beta actin
HGM+10%FBS-24h
HGM-24h
HGM-12h+ (HGM+10% FBS-12h)
I
sWAT transcriptome
Transcription factor activity cohen's D
High in Ad lib
High in Fast
High in Refeed
J
eWAT transcriptome
K
Preadipocytes
SREBP1 Relative mRNA level (Nomalized to 18s)
si-NC_Feed
si-NC_Fast
si-SREBP1_Fast
L
HGM+10%FBS
LGM+10%FBS
M
Relative luciferase activity
N
Glucose (g/L)
FBS
SLC25A35
Beta-actin
O
HGM+10%FBS
HGM+10%FBS+0.5mM FA
P
Luciferase assay
HGM+10%FBS+ 450μmol (PA+OA)
HGM+500μmol (PA+OA)
Q
FA (μmol):
PA+OA:
FBS:
GAPDH
SLC25A35
SLC25A35 relative expression ( normalized to GAPDH)
HGM+300μmol PA+OA
HGM+450μmol PA+OA

### Figure S3

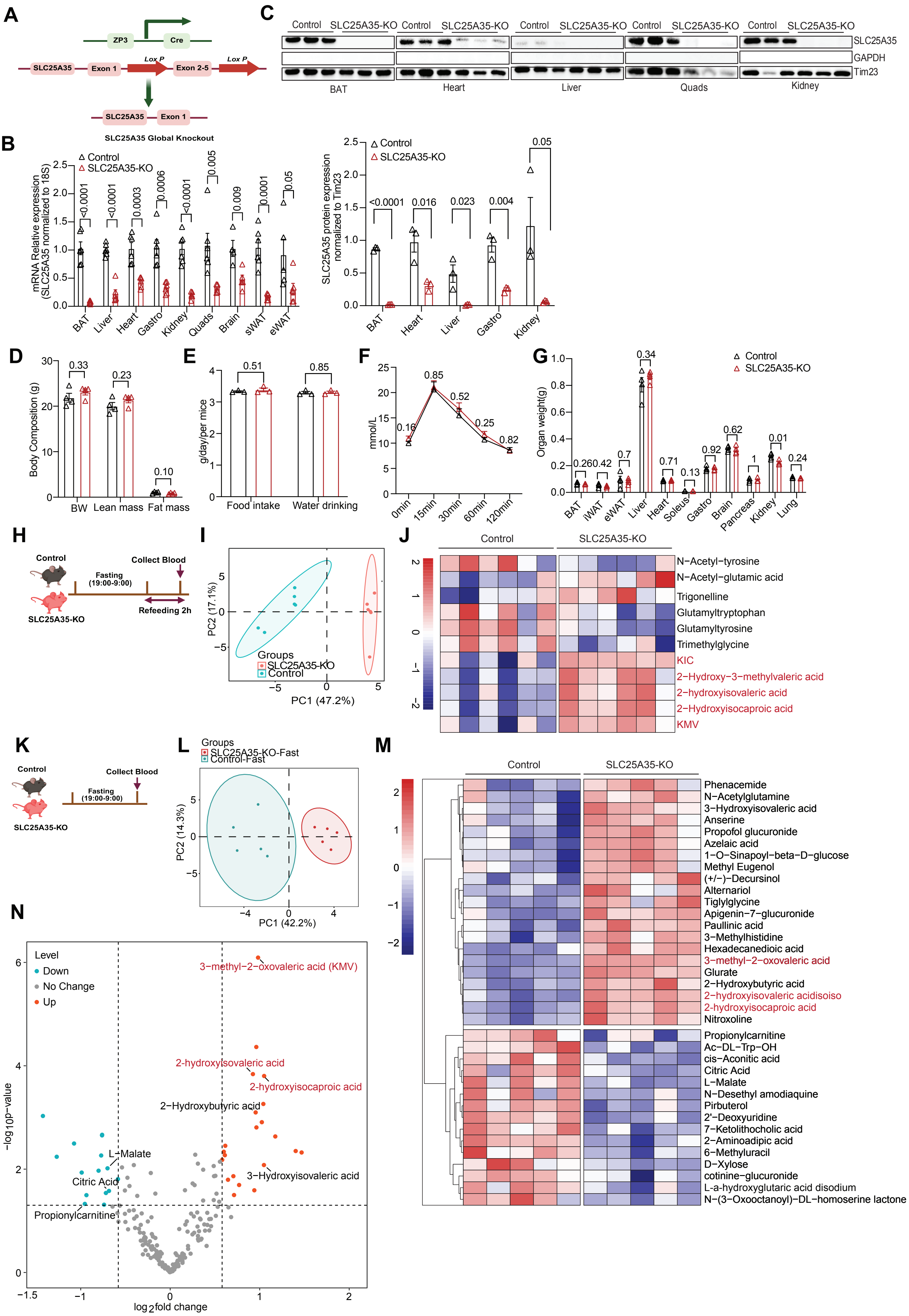
A
ZP3
Cre
SLC25A35
Exon 1
Lox P
Exon 2-5
Lox P
SLC25A35
Exon 1
SLC25A35 Global Knockout
C
Control
SLC25A35-KO
SLC25A35
GAPDH
Tim23
BAT
Heart
Liver
Quads
Kidney
B
Control
SLC25A35-KO
mRNA Relative expression (SLC25A35 normalized to 18S)
BAT
Liver
Heart
Gastro
Kidney
Quads
Brain
sWAT
eWAT
SLC25A35 protein expression normalized to Tim23
D
Body Composition (g)
BW
Lean mass
Fat mass
E
g/day/per mice
Food intake
Water drinking
F
mmol/L
0min
15min
30min
60min
120min
G
Organ weight(g)
Control
SLC25A35-KO
BAT
iWAT
eWAT
Liver
Heart
Soleus
Gastro
Brain
Pancreas
Kidney
Lung
H
Control
Collect Blood
Fasting (19:00-9:00)
Refeeding 2h
SLC25A35-KO
I
PC2 (17.1%)
PC1 (47.2%)
Groups
SLC25A35-KO
Control
J
Control
SLC25A35-KO
N-Acetyl-tyrosine
N-Acetyl-glutamic acid
Trigonelline
Glutamyltryptophan
Glutamyltyrosine
Trimethylglycine
KIC
2-Hydroxy-3-methylvaleric acid
2-hydroxyisovaleric acid
2-Hydroxyisocaproic acid
KMV
K
Control
Collect Blood
Fasting (19:00-9:00)
SLC25A35-KO
L
Groups
SLC25A35-KO-Fast
Control-Fast
PC2 (14.3%)
PC1 (42.2%)
M
Control
SLC25A35-KO
Phenacemide
N-Acetylglutamine
3-Hydroxyisovaleric acid
Anserine
Propofol glucuronide
Azelaic acid
1-O-Sinapoyl-beta-D-glucose
Methyl Eugenol
(+/-)-Decursinol
Alternariol
Tiglylglycine
Apigenin-7-glucuronide
Paullinic acid
3-Methylhistidine
Hexadecanedioic acid
3-methyl-2-oxovaleric acid
Glurate
2-Hydroxybutyric acid
2-hydroxyisovaleric acidisoiso
2-hydroxyisocaproic acid
Nitroxoline
Propionylcarnitine
Ac-DL-Trp-OH
cis-Aconitic acid
Citric Acid
L-Malate
N-Desethyl amodiaquine
Pirbuterol
2'-Deoxyuridine
7-Ketolithocholic acid
2-Aminoadipic acid
6-Methyluracil
D-Xylose
cotinine-glucuronide
L-a-hydroxyglutaric acid disodium
N-(3-Oxooctanoyl)-DL-homoserine lactone
N
Level
Down
No Change
Up
3-methyl-2-oxovaleric acid (KMV)
2-hydroxyisovaleric acid
2-hydroxyisocaproic acid
2-Hydroxybutyric acid
L-Malate
Citric Acid
3-Hydroxyisovaleric acid
Propionylcarnitine
-log10p-value
log2fold change

### Figure S4

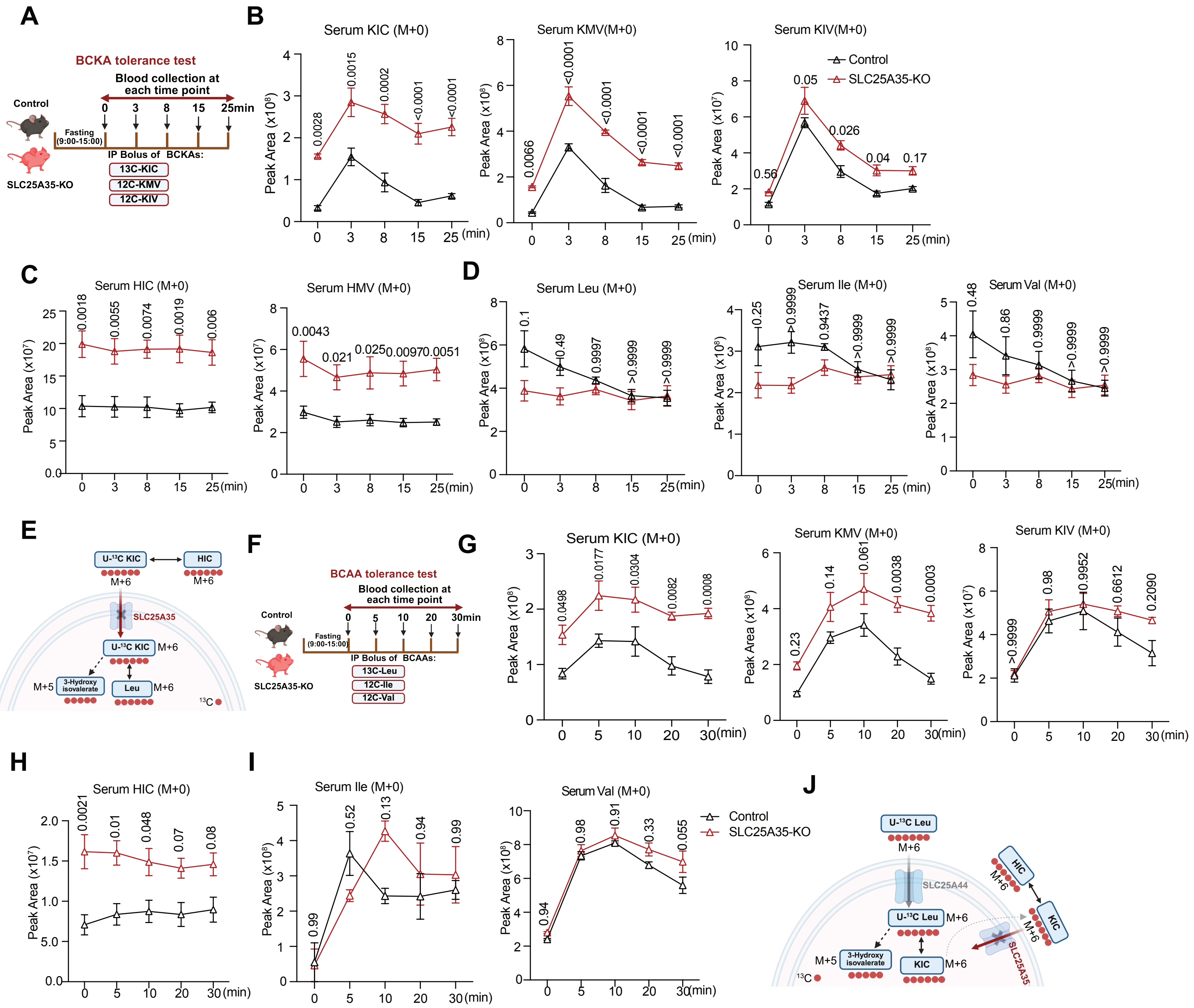
A
BCKA tolerance test
Blood collection at each time point
Control
SLC25A35-KO
Fasting (9:00-15:00)
IP Bolus of BCKAs:
13C-KIC
12C-KMV
12C-KIV
B
Serum KIC (M+0)
Serum KMV(M+0)
Serum KIV(M+0)
Control
SLC25A35-KO
Peak Area
C
Serum HIC (M+0)
Serum HMV (M+0)
D
Serum Leu (M+0)
Serum Ile (M+0)
Serum Val (M+0)
E
U-13C KIC
HIC
M+6
SLC25A35
3-Hydroxy isovalerate
Leu
M+5
F
BCAA tolerance test
IP Bolus of BCAAs:
13C-Leu
12C-Ile
12C-Val
G
Serum KIC (M+0)
Serum KMV (M+0)
Serum KIV (M+0)
H
Serum HIC (M+0)
I
Serum Ile (M+0)
Serum Val (M+0)
J
U-13C Leu
SLC25A44
KIC
HIC

### Figure S5

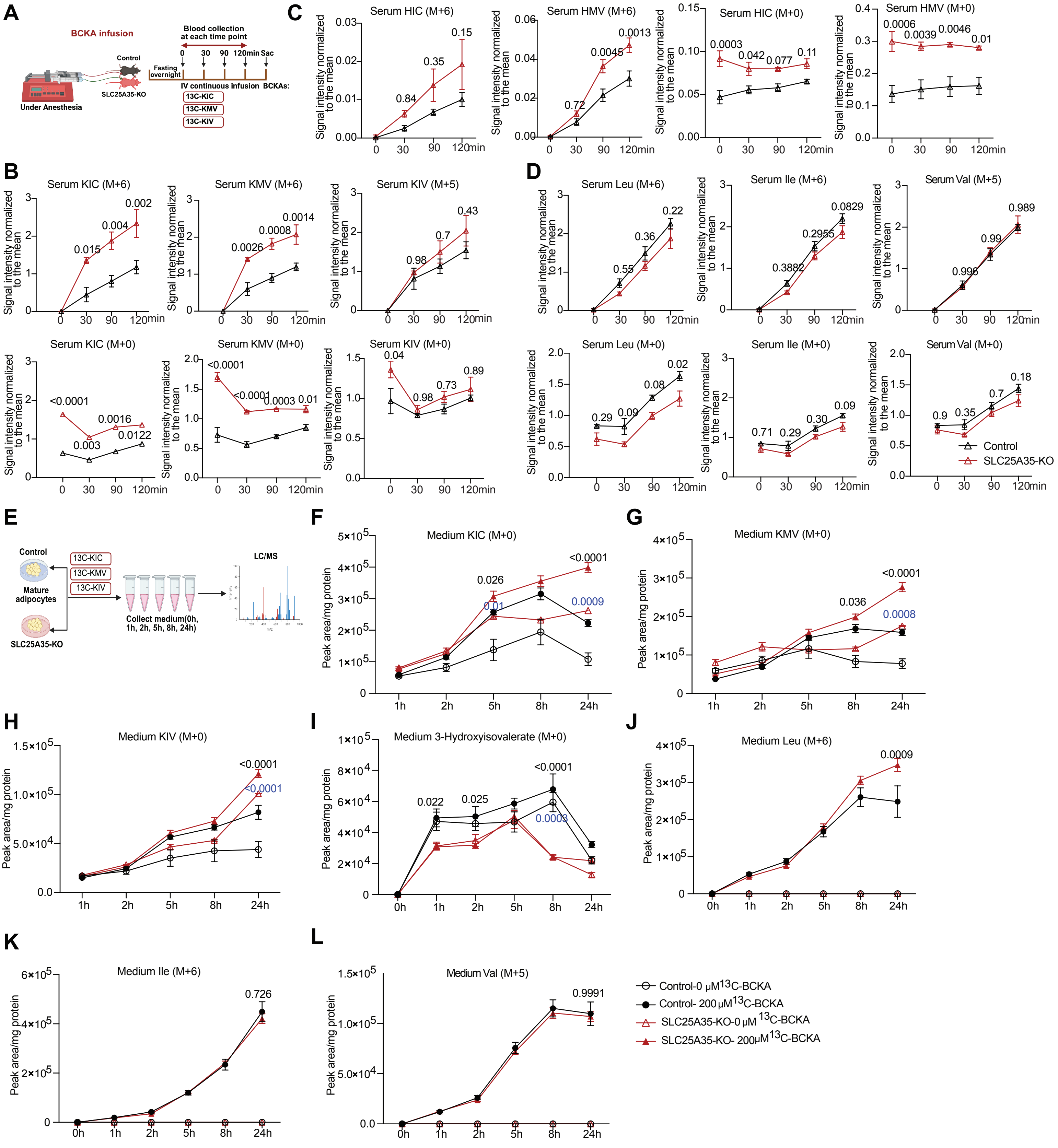

A
BCKA infusion
Blood collection at each time point
Control
SLC25A35-KO
Under Anesthesia
Fasting overnight
0 30 90 120min Sac
IV continuous infusion BCKAs:
13C-KIC
13C-KMV
13C-KIV
B
Serum KIC (M+6)
Serum KMV (M+6)
Serum KIV (M+5)
Serum KIC (M+0)
Serum KMV (M+0)
Serum KIV (M+0)
Signal intensity normalized to the mean
C
Serum HIC (M+6)
Serum HMV (M+6)
Serum HIC (M+0)
Serum HMV (M+0)
D
Serum Leu (M+6)
Serum Ile (M+6)
Serum Val (M+5)
Serum Leu (M+0)
Serum Ile (M+0)
Serum Val (M+0)
Control
SLC25A35-KO
E
Control
Mature adipocytes
SLC25A35-KO
Collect medium(0h, 1h, 2h, 5h, 8h, 24h)
LC/MS
F
Medium KIC (M+0)
G
Medium KMV (M+0)
H
Medium KIV (M+0)
I
Medium 3-Hydroxyisovalerate (M+0)
J
Medium Leu (M+6)
K
Medium Ile (M+6)
L
Medium Val (M+5)
Peak area/mg protein
Control-0 μM 13C-BCKA
Control- 200 μM 13C-BCKA
SLC25A35-KO-0 μM 13C-BCKA
SLC25A35-KO- 200μM 13C-BCKA

### Figure S6

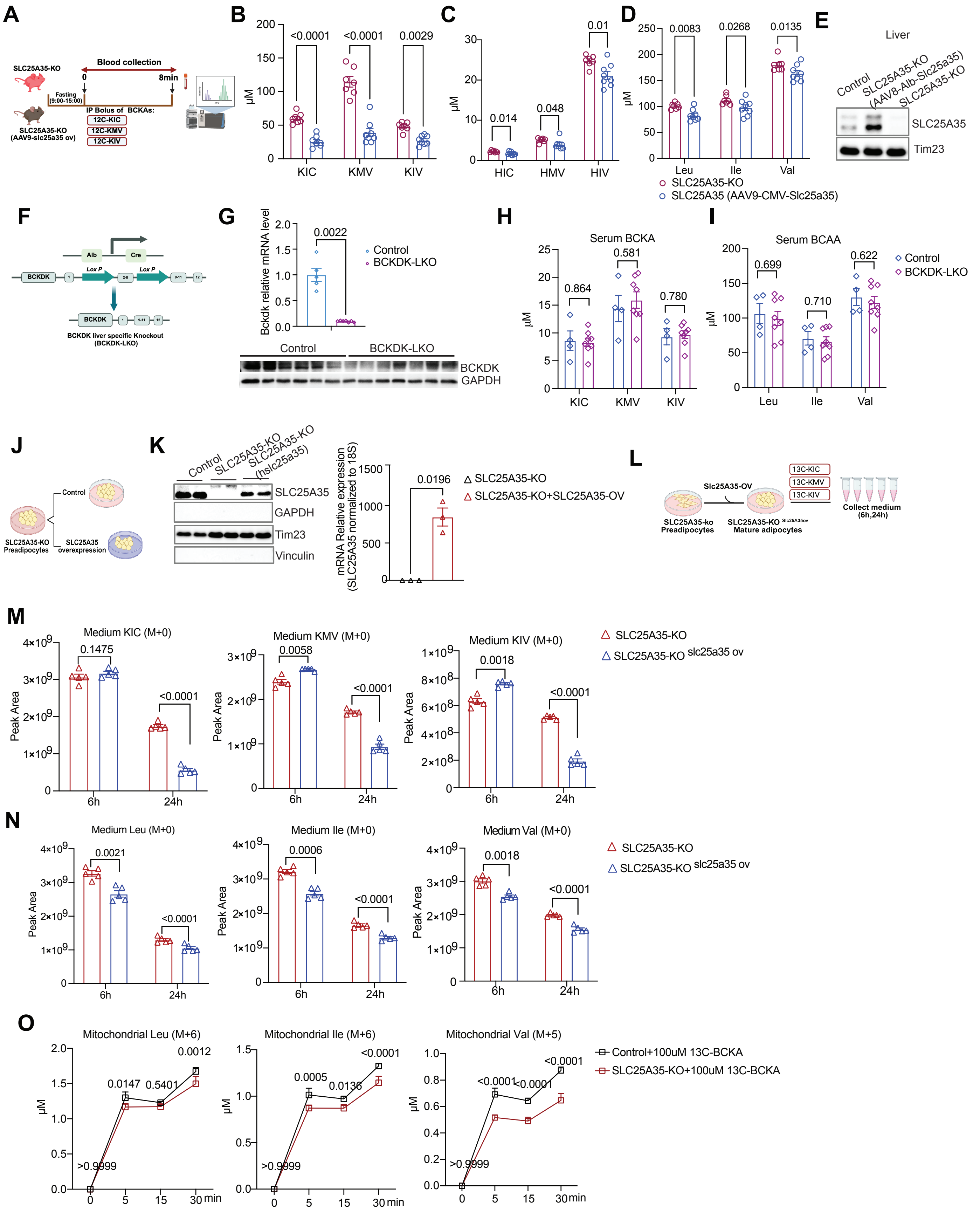

A
SLC25A35-KO
Blood collection
0
8min
Fasting (9:00-15:00)
IP Bolus of BCKAs:
12C-KIC
12C-KMV
12C-KIV
SLC25A35-KO (AAV9-slc25a35 ov)
B
μM
<0.0001
<0.0001
0.0029
KIC
KMV
KIV
C
μM
0.014
0.048
0.01
HIC
HMV
HIV
D
μM
0.0083
0.0268
0.0135
Leu
Ile
Val
SLC25A35-KO
SLC25A35 (AAV9-CMV-Slc25a35)
E
Liver
Control
SLC25A35-KO (AAV8-Alb-Slc25a35)
SLC25A35-KO
SLC25A35
Tim23
F
Alb
Cre
Lox P
Lox P
BCKDK
BCKDK
BCKDK liver specific Knockout (BCKDK-LKO)
G
Bckdk relative mRNA level
0.0022
Control
BCKDK-LKO
Control
BCKDK-LKO
BCKDK
GAPDH
H
Serum BCKA
μM
0.864
0.581
0.780
KIC
KMV
KIV
I
Serum BCAA
μM
0.699
0.710
0.622
Leu
Ile
Val
Control
BCKDK-LKO
J
Control
SLC25A35-KO Preadipocytes
SLC25A35 overexpression
K
Control
SLC25A35-KO
SLC25A35-KO (hslc25a35)
SLC25A35
GAPDH
Tim23
Vinculin
mRNA Relative expression (SLC25A35 normalized to 18S)
0.0196
SLC25A35-KO
SLC25A35-KO+SLC25A35-OV
L
Slc25A35-OV
13C-KIC
13C-KMV
13C-KIV
SLC25A35-ko Preadipocytes
SLC25A35-KO Slc25A35ov Mature adipocytes
Collect medium (6h,24h)
M
Medium KIC (M+0)
Medium KMV (M+0)
Medium KIV (M+0)
Peak Area
0.1475
<0.0001
0.0058
<0.0001
0.0018
<0.0001
6h
24h
SLC25A35-KO
SLC25A35-KO slc25a35 ov
N
Medium Leu (M+0)
Medium Ile (M+0)
Medium Val (M+0)
Peak Area
0.0021
<0.0001
0.0006
<0.0001
0.0018
<0.0001
6h
24h
SLC25A35-KO
SLC25A35-KO slc25a35 ov
O
Mitochondrial Leu (M+6)
Mitochondrial Ile (M+6)
Mitochondrial Val (M+5)
μM
0.0012
0.0147
0.5401
>0.9999
<0.0001
0.0005
0.0136
>0.9999
<0.0001
<0.0001
<0.0001
>0.9999
min
Control+100uM 13C-BCKA
SLC25A35-KO+100uM 13C-BCKA

### Figure S7

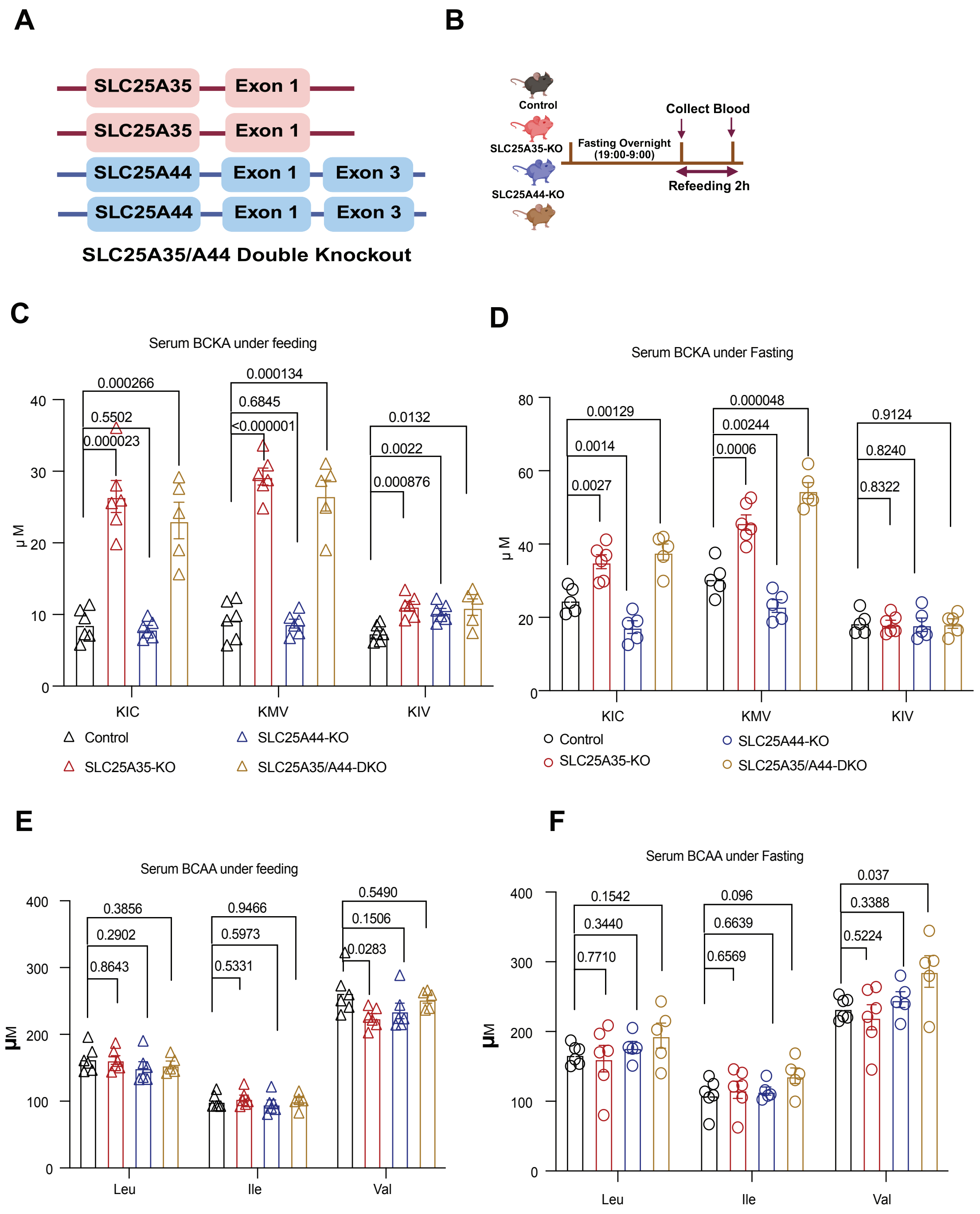
A
SLC25A35
Exon 1
SLC25A44
Exon 3
SLC25A35/A44 Double Knockout
B
Control
SLC25A35-KO
SLC25A44-KO
Fasting Overnight (19:00-9:00)
Collect Blood
Refeeding 2h
C
Serum BCKA under feeding
μM
KIC
KMV
KIV
0.000266
0.5502
0.000023
0.000134
0.6845
<0.000001
0.0132
0.0022
0.000876
Control
SLC25A35-KO
SLC25A44-KO
SLC25A35/A44-DKO
D
Serum BCKA under Fasting
0.00129
0.0014
0.0027
0.000048
0.00244
0.0006
0.9124
0.8240
0.8322
E
Serum BCAA under feeding
Leu
Ile
Val
0.3856
0.2902
0.8643
0.9466
0.5973
0.5331
0.5490
0.1506
0.0283
F
Serum BCAA under Fasting
0.1542
0.3440
0.7710
0.096
0.6639
0.6569
0.037
0.3388
0.5224

### Figure S8

A

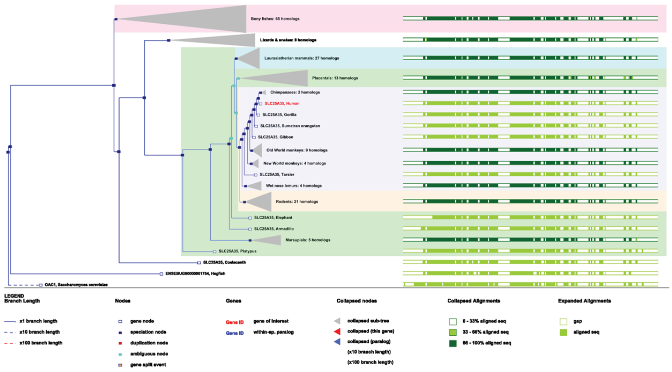

B

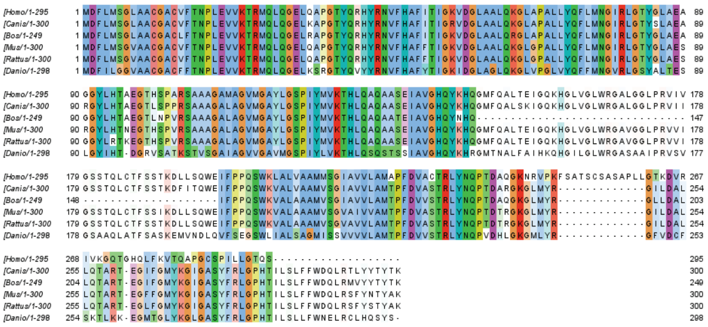

C

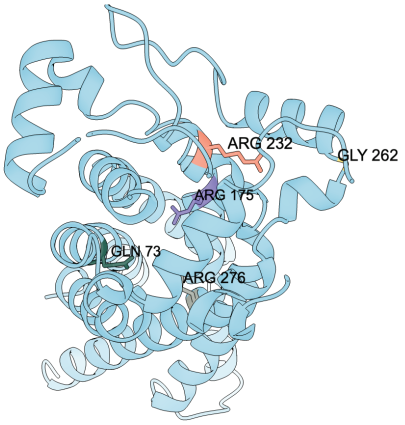


D

SLC25A35 mutagenesis design

Candidate residues selected from ligand-binding, conservation, and human-variant evidence

| Mutation | Wild-type residue | Substitution | Rationale |
| --- | --- | --- | --- |
| R175A | Arg (R) | Ala (A) | ● KIC-binding site |
| R232A | Arg (R) | Ala (A) | ● Conserved site |
| G262A | Gly (G) | Ala (A) | ● SNP-associated site |
| Q73A / R276A | Gln (Q) / Arg (R) | Ala (A) / Ala (A) | ● KIC-binding sites |

### Figure S9

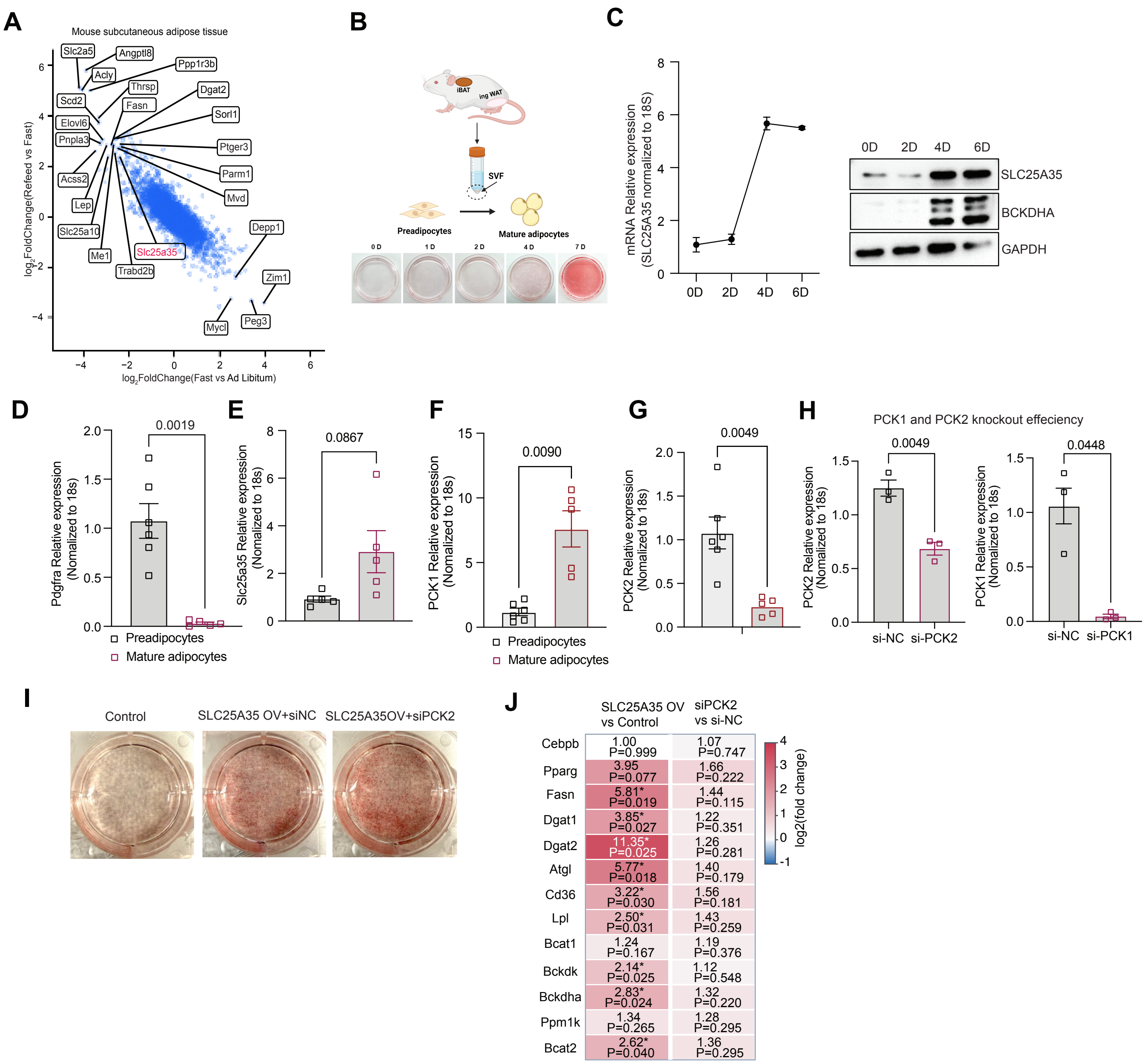

A
Mouse subcutaneous adipose tissue
log2FoldChange(Refeed vs Fast)
log2FoldChange(Fast vs Ad Libitum)
Slc2a5
Angptl8
Ppp1r3b
Acly
Thrsp
Dgat2
Scd2
Fasn
Sorl1
Elovl6
Pnpla3
Ptger3
Parm1
Acss2
Mvd
Lep
Depp1
Slc25a10
Me1
Slc25a35
Trabd2b
Zim1
Peg3
Mycl
B
iBAT
ing WAT
SVF
Preadipocytes
Mature adipocytes
0 D
1 D
2 D
4 D
7 D
C
mRNA Relative expression (SLC25A35 normalized to 18S)
0D
2D
4D
6D
SLC25A35
BCKDHA
GAPDH
D
Pdgfra Relative expression (Nomalized to 18s)
0.0019
Preadipocytes
Mature adipocytes
E
Slc25a35 Relative expression (Nomalized to 18s)
0.0867
F
PCK1 Relative expression (Nomalized to 18s)
0.0090
Preadipocytes
Mature adipocytes
G
PCK2 Relative expression (Nomalized to 18s)
0.0049
H
PCK1 and PCK2 knockout effeciency
PCK2 Relative expression (Nomalized to 18s)
0.0049
si-NC
si-PCK2
PCK1 Relative expression (Nomalized to 18s)
0.0448
si-NC
si-PCK1
I
Control
SLC25A35 OV+siNC
SLC25A35OV+siPCK2
J
SLC25A35 OV vs Control
siPCK2 vs si-NC
log2(fold change)
Cebpb 1.00 P=0.999 1.07 P=0.747
Pparg 3.95 P=0.077 1.66 P=0.222
Fasn 5.81* P=0.019 1.44 P=0.115
Dgat1 3.85* P=0.027 1.22 P=0.351
Dgat2 11.35* P=0.025 1.26 P=0.281
Atgl 5.77* P=0.018 1.40 P=0.179
Cd36 3.22* P=0.030 1.56 P=0.181
Lpl 2.50* P=0.031 1.43 P=0.259
Bcat1 1.24 P=0.167 1.19 P=0.376
Bckdk 2.14* P=0.025 1.12 P=0.548
Bckdha 2.83* P=0.024 1.32 P=0.220
Ppm1k 1.34 P=0.265 1.28 P=0.295
Bcat2 2.62* P=0.040 1.36 P=0.295

### Figure S10

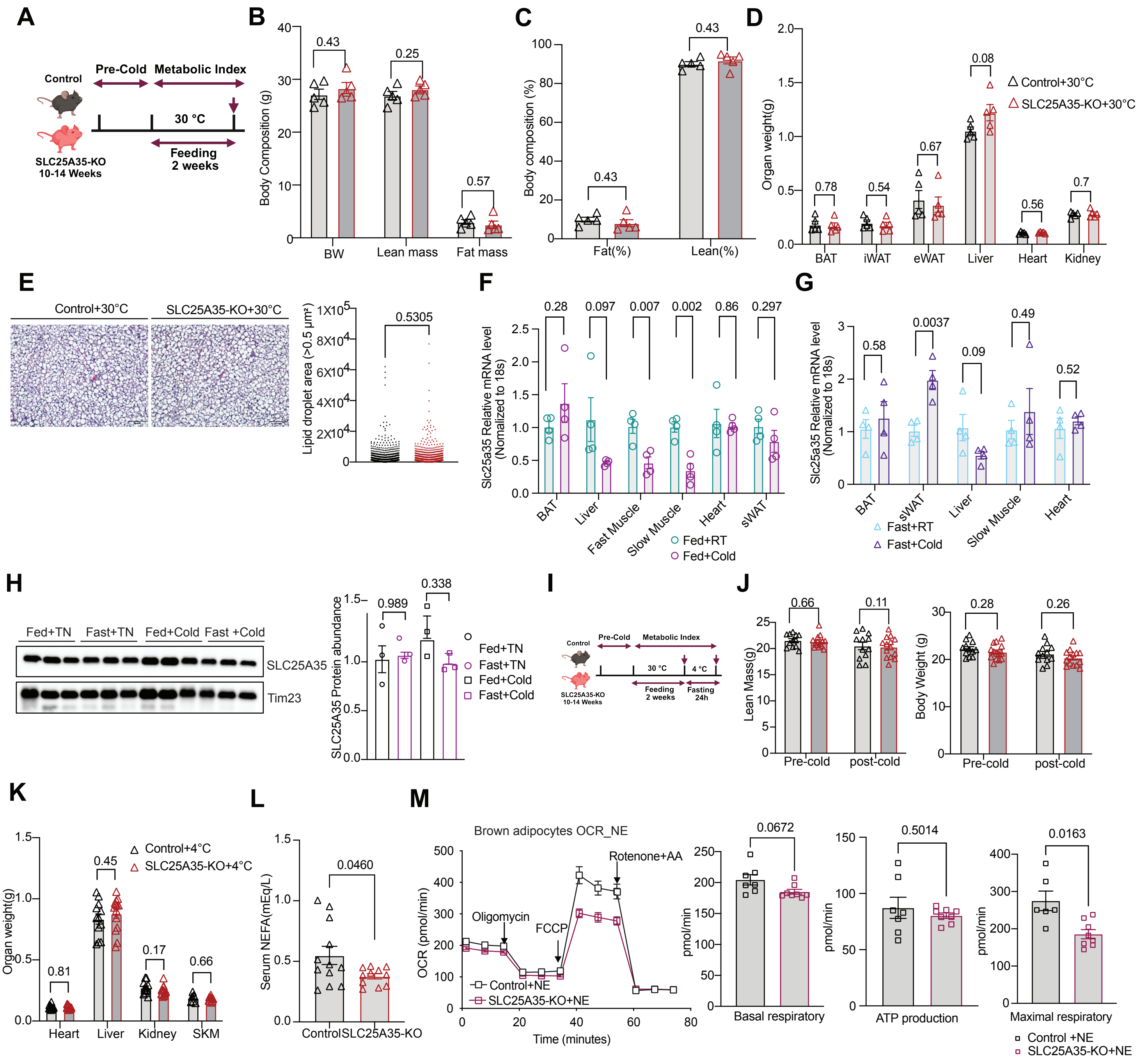

A
Control
Pre-Cold
Metabolic Index
30 °C
Feeding
2 weeks
SLC25A35-KO
10-14 Weeks
B
Body Composition (g)
BW
Lean mass
Fat mass
0.43
0.25
0.57
C
Body composition (%)
Fat(%)
Lean(%)
0.43
0.43
D
Organ weight(g)
Control+30°C
SLC25A35-KO+30°C
BAT
iWAT
eWAT
Liver
Heart
Kidney
0.78
0.54
0.67
0.08
0.56
0.7
E
Control+30°C
SLC25A35-KO+30°C
Lipid droplet area (>0.5 μm²)
0.5305
F
Slc25a35 Relative mRNA level (Nomalized to 18s)
0.28
0.097
0.007
0.002
0.86
0.297
BAT
Liver
Fast Muscle
Slow Muscle
Heart
sWAT
Fed+RT
Fed+Cold
G
Slc25a35 Relative mRNA level (Nomalized to 18s)
0.58
0.0037
0.09
0.49
0.52
BAT
sWAT
Liver
Slow Muscle
Heart
Fast+RT
Fast+Cold
H
Fed+TN
Fast+TN
Fed+Cold
Fast +Cold
SLC25A35
Tim23
SLC25A35 Protein abundance
0.989
0.338
Fed+TN
Fast+TN
Fed+Cold
Fast+Cold
I
Control
Pre-Cold
Metabolic Index
30 °C
4 °C
Feeding
2 weeks
Fasting
24h
SLC25A35-KO
10-14 Weeks
J
Lean Mass(g)
0.66
0.11
Pre-cold
post-cold
Body Weight (g)
0.28
0.26
Pre-cold
post-cold
K
Organ weight(g)
Control+4°C
SLC25A35-KO+4°C
0.81
0.45
0.17
0.66
Heart
Liver
Kidney
SKM
L
Serum NEFA(mEq/L)
0.0460
Control
SLC25A35-KO
M
Brown adipocytes OCR_NE
OCR (pmol/min)
Oligomycin
FCCP
Rotenone+AA
Control+NE
SLC25A35-KO+NE
Time (minutes)
pmol/min
0.0672
Basal respiratory
0.5014
ATP production
0.0163
Maximal respiratory
Control +NE
SLC25A35-KO+NE

### Figure S11

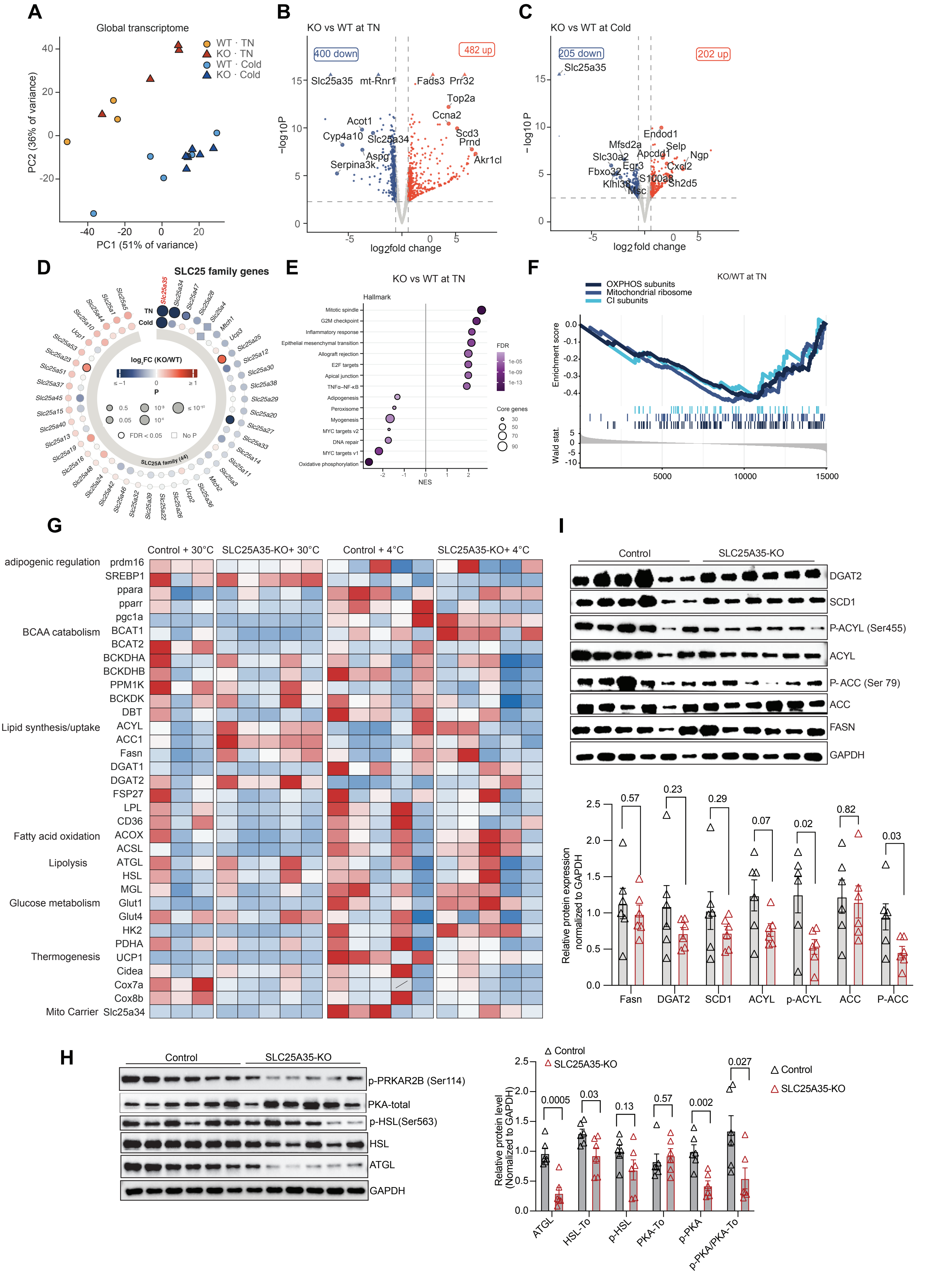
A
Global transcriptome
WT · TN
KO · TN
WT · Cold
KO · Cold
PC2 (36% of variance)
PC1 (51% of variance)
B
KO vs WT at TN
400 down
482 up
Slc25a35
mt-Rnr1
Fads3
Prr32
Top2a
Ccna2
Acot1
Cyp4a10
Slc25a34
Scd3
Prnd
Aspg
Serpina3k
Akr1cl
log2fold change
C
KO vs WT at Cold
205 down
202 up
Slc25a35
Endod1
Mfsd2a
Selp
Apcdd1
Ngp
Slc30a2
Egr3
Cxcl2
Fbxo32
S100a8
Klhl38
Sh2d5
Msc
D
SLC25 family genes
TN
Cold
SLC25A family (44)
E
KO vs WT at TN
Hallmark
Mitotic spindle
G2M checkpoint
Inflammatory response
Epithelial mesenchymal transition
Allograft rejection
E2F targets
Apical junction
TNFα–NF-κB
Adipogenesis
Peroxisome
Myogenesis
MYC targets v2
DNA repair
MYC targets v1
Oxidative phosphorylation
NES
FDR
Core genes
F
KO/WT at TN
OXPHOS subunits
Mitochondrial ribosome
CI subunits
Enrichment score
Wald stat.
G
Control + 30°C
SLC25A35-KO+ 30°C
Control + 4°C
SLC25A35-KO+ 4°C
adipogenic regulation
prdm16
SREBP1
ppara
pparr
pgc1a
BCAA catabolism
BCAT1
BCAT2
BCKDHA
BCKDHB
PPM1K
BCKDK
DBT
Lipid synthesis/uptake
ACYL
ACC1
Fasn
DGAT1
DGAT2
FSP27
LPL
CD36
Fatty acid oxidation
ACOX
ACSL
Lipolysis
ATGL
HSL
MGL
Glucose metabolism
Glut1
Glut4
HK2
PDHA
Thermogenesis
UCP1
Cidea
Cox7a
Cox8b
Mito Carrier
Slc25a34
I
Control
SLC25A35-KO
DGAT2
SCD1
P-ACYL (Ser455)
ACYL
P- ACC (Ser 79)
ACC
FASN
GAPDH
Relative protein expression normalized to GAPDH
Fasn
DGAT2
SCD1
ACYL
p-ACYL
ACC
P-ACC
H
Control
SLC25A35-KO
p-PRKAR2B (Ser114)
PKA-total
p-HSL(Ser563)
HSL
ATGL
GAPDH
Relative protein level (Nomalized to GAPDH)
ATGL
HSL-To
p-HSL
PKA-To
p-PKA
p-PKA/PKA-To

### Figure S12

**A**

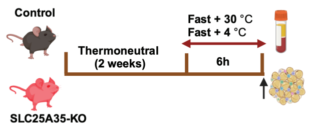


**B**

Serum BCKA metabolites

| | Control + 30°C | A35-KO + 30°C | Control + 4°C | A35-KO + 4°C |
|---|---|---|---|---|
| KIC | 19.35 | 58.34 p=0.0002 | 10.68 | 29.67 p<0.0001 |
| KMV | 22.34 | 65.25 p=0.0019 | 9.43 | 14.22 p=0.175 |
| KIV | 24.48 | 26.32 p=0.930 | 16.85 | 15.31 p=0.503 |
| HIC | 1.32 | 2.88 p=0.0001 | 1.70 | 1.70 p=1.000 |
| HMV | 4.35 | 7.68 p=0.066 | 1.34 | 2.17 p=0.064 |
| HIV | 22.12 | 25.59 p=0.557 | 14.59 | 16.14 p=0.821 |
| Leu | 175.6 | 188.5 p=0.971 | 355.1 | 450.0 p=0.246 |
| Ile | 128.3 | 161.8 p=0.420 | 182.2 | 195.6 p=0.662 |
| Val | 234.6 | 297.5 p=0.687 | 399.3 | 473.0 p=0.419 |
| Acetoacetate | 101.4 | 92.30 p=0.888 | 115.0 | 40.93 p=0.0004 |
| 3-HB | 547.7 | 569.2 p=0.993 | 868.7 | 489.2 p=0.0044 |
| 3-Aminoisobutyric acid | 3.10 | 3.46 p=0.925 | 4.29 | 2.99 p=0.071 |
| 3-HMG | 0.228 | 0.157 p=0.388 | 0.290 | 0.157 p=0.044 |
| 3-HIB | 34.04 | 24.29 p=0.265 | 35.34 | 35.34 p=1.000 |
| HMB | 1.16 | 1.53 p=0.118 | 1.53 | 2.83 p<0.0001 |

Row Z-score: 2, 1, 0, -1, -2

**C**

BAT BCKA metabolites

| | Control + 30°C | A35-KO + 30°C | Control + 4°C | A35-KO + 4°C |
|---|---|---|---|---|
| KIC | 0.572 | 0.659 p=0.891 | 1.33 | 1.42 p=0.761 |
| KMV | 0.215 | 0.265 p=0.797 | 0.468 | 0.600 p=0.743 |
| HIC | 0.195 | 0.334 p=0.127 | 0.182 | 0.577 p=0.0001 |
| HMV | 0.689 | 0.863 p=0.708 | 0.381 | 0.698 p=0.143 |
| HIV | 5.19 | 4.23 p=0.694 | 5.52 | 6.46 p=0.985 |
| Leu | 282.5 | 246.8 p=0.753 | 263.0 | 315.4 p=0.353 |
| Ile | 144.9 | 102.6 p=0.263 | 132.6 | 154.5 p=0.396 |
| Val | 236.9 | 182.3 p=0.481 | 252.2 | 278.2 p=0.514 |
| Acetoacetate | 13.25 | 18.59 p=0.433 | 30.09 | 19.44 p=0.273 |
| 3-HB | 206.0 | 144.7 p=0.390 | 459.0 | 306.8 p=0.129 |
| 3-HMG | 0.036 | 0.019 p=0.198 | 0.431 | 0.226 p=0.192 |
| 3-Aminoisobutyric aci | 0.831 | 0.792 p=0.926 | 1.59 | 1.05 p=0.249 |
| 3-HIB | 6.44 | 3.58 p=0.214 | 11.73 | 8.67 p=0.533 |
| 3-HMB | 0.836 | 0.456 p=0.540 | 1.00 | 2.70 p=0.223 |

Row Z-score: 2, 1, 0, -1, -2
